# AlphaGenome: advancing regulatory variant effect prediction with a unified DNA sequence model

**DOI:** 10.1101/2025.06.25.661532

**Authors:** Žiga Avsec, Natasha Latysheva, Jun Cheng, Guido Novati, Kyle R. Taylor, Tom Ward, Clare Bycroft, Lauren Nicolaisen, Eirini Arvaniti, Joshua Pan, Raina Thomas, Vincent Dutordoir, Matteo Perino, Soham De, Alexander Karollus, Adam Gayoso, Toby Sargeant, Anne Mottram, Lai Hong Wong, Pavol Drotár, Adam Kosiorek, Andrew Senior, Richard Tanburn, Taylor Applebaum, Souradeep Basu, Demis Hassabis, Pushmeet Kohli

**Author notes:** (Z.A.); (P.K.). These authors contributed equally to this work (Žiga Avsec, Natasha Latysheva, Jun Cheng, Guido Novati, Kyle R. Taylor, Tom Ward, Clare Bycroft, Lauren Nicolaisen, Eirini Arvaniti).

## Abstract

Deep learning models that predict functional genomic measurements from DNA sequence are powerful tools for deciphering the genetic regulatory code. Existing methods trade off between input sequence length and prediction resolution, thereby limiting their modality scope and performance. We present AlphaGenome, which takes as input 1 megabase of DNA sequence and predicts thousands of functional genomic tracks up to single base pair resolution across diverse modalities – including gene expression, transcription initiation, chromatin accessibility, histone modifications, transcription factor binding, chromatin contact maps, splice site usage, and splice junction coordinates and strength. Trained on human and mouse genomes, AlphaGenome matches or exceeds the strongest respective available external models on 24 out of 26 evaluations on variant effect prediction. AlphaGenome’s ability to simultaneously score variant effects across all modalities accurately recapitulates the mechanisms of clinically-relevant variants near the *TAL1* oncogene. To facilitate broader use, we provide tools for making genome track and variant effect predictions from sequence.

## Introduction

Interpreting the impact of genome sequence variation remains a central biological challenge. Non-coding variants, which reside outside of protein-coding regions, are particularly challenging to interpret due to the diverse molecular consequences they can elicit. For example, non-coding variants can modulate genome properties such as chromatin accessibility, epigenetic modifications, and 3D chromatin conformation. Variants can further influence messenger RNA (mRNA) availability by altering expression levels or modifying sequence composition through splicing changes. Additionally, variants can exhibit cell type or tissue-specific effects. Given that over 98% of observed genetic variation in humans is non-coding^1^, global characterization of the complex effects of this vast majority of variants remains intractable without computational predictions.

Computational methods can learn patterns from experimental data to predict and explain variant effects. One class of methods, sequence-to-function models^2–6^, takes a DNA sequence as input and predicts *genome tracks* – a data format associating each DNA base pair with a value (representing read coverage, count, or signal) derived from experimental assays performed in cell lines or tissues. Genome tracks span various data *modalities* measuring gene expression (with *output types* comprising RNA-seq, CAGE-seq, PRO-cap), splicing (splice sites, splice site usage, splice junctions), DNA accessibility (DNase-seq, ATAC-seq), histone modification (ChIP-seq), transcription factor binding (TF ChIP-seq), or chromatin conformation (Hi-C/micro-C). Successfully trained sequence-to-function models accurately predict experimental measurements from input sequences. Furthermore, by comparing genome track predictions from an alternative sequence versus a reference sequence, these models can predict the molecular effects of variants.

Currently, deep learning-based sequence-to-function models face two fundamental tradeoffs constraining their ability to predict how variants affect diverse modes of biological regulation. First, often due to computational limitations, models must trade off between capturing long-range genomic interactions and achieving nucleotide-level predictive resolution. While models like SpliceAI^5^, BPNet^7^, ProCapNet^8^, and APARENT2^9^ provide base-resolution predictions, they are restricted to short input sequences (e.g., 10kb or less), thus missing the influence of distal regulatory elements. Conversely, models such as Enformer^2^ and Borzoi^3^ can process longer sequences (∼200kb to ∼500kb) to capture broader context, but at the cost of reducing output resolution (128bp or 32bp bins), which can blur fine-scale regulatory features like splice sites, transcription factor footprints or polyadenylation sites.

A second tradeoff exists between capturing diverse modalities versus specializing in one or a few. Several state-of-the-art (SOTA) models are highly specialized for single modalities, such as SpliceAI^5^ for splice site prediction, ChromBPNet^10^ for local chromatin accessibility, or Orca^4^ for 3D genome architecture. However, specialized models alone are insufficient for capturing the diverse molecular consequences of variants across modalities. Even within a single modality like splicing, specialized models such as SpliceAI^5^ or Pangolin^11^ predict certain aspects (such as splice site prediction) while omitting others (such as splice junction prediction or competition between splice sites). Models like DeepSEA^12^, Basenji^13^, Enformer^2^, Sei^14^, and Borzoi^3^ have demonstrated the utility and practicality of multimodal models. They allow users to employ a single model for several modalities, instead of requiring multiple specialized models. Furthermore, their learned general sequence representation enables them to be readily fine-tuned for new tasks^15,16^. However, these more generalist models can lag behind their specialized counterparts on certain tasks, like splicing, or may lack particular modalities, such as contact maps.

Here we present AlphaGenome, a model that unifies multimodal prediction, long sequence context, and base-pair resolution into a single framework. The model takes 1 megabase (Mb) of DNA sequence as input and predicts a diverse range of genome tracks across numerous cell types. AlphaGenome’s splicing predictions include a novel splice junction prediction approach alongside splice site usage prediction. We evaluated AlphaGenome’s performance using a comprehensive set of benchmarks, covering both its ability to accurately predict genome tracks on previously unseen DNA sequences and its effectiveness in variant effect prediction tasks. AlphaGenome achieved SOTA performance on 22 out of 24 genome track prediction tasks and 24 out of 26 variant effect prediction tasks. We performed extensive ablations of target resolution, sequence length, distillation, and modality combinations to explain AlphaGenome’s performance and inform design choices for future sequence-to-function models. We envisage AlphaGenome will provide a powerful and extensible foundation for analyzing the regulatory code within the genome.

We first present key technical details of the AlphaGenome data and training procedure, alongside a high-level summary of our evaluations (Fig. 1). We then demonstrate high-fidelity genome track prediction performance, a prerequisite for variant effect prediction (Fig. 2). Next, we focus on variant effect prediction with modality-specific deep dives into splicing (Fig. 3), gene expression (Fig. 4), and chromatin accessibility (Fig. 5). Finally, we highlight the model’s utility in cross-modality variant interpretation (Fig. 6) and dissect the impact of modeling choices on AlphaGenome’s performance (Fig. 7).

**Figure 1.**
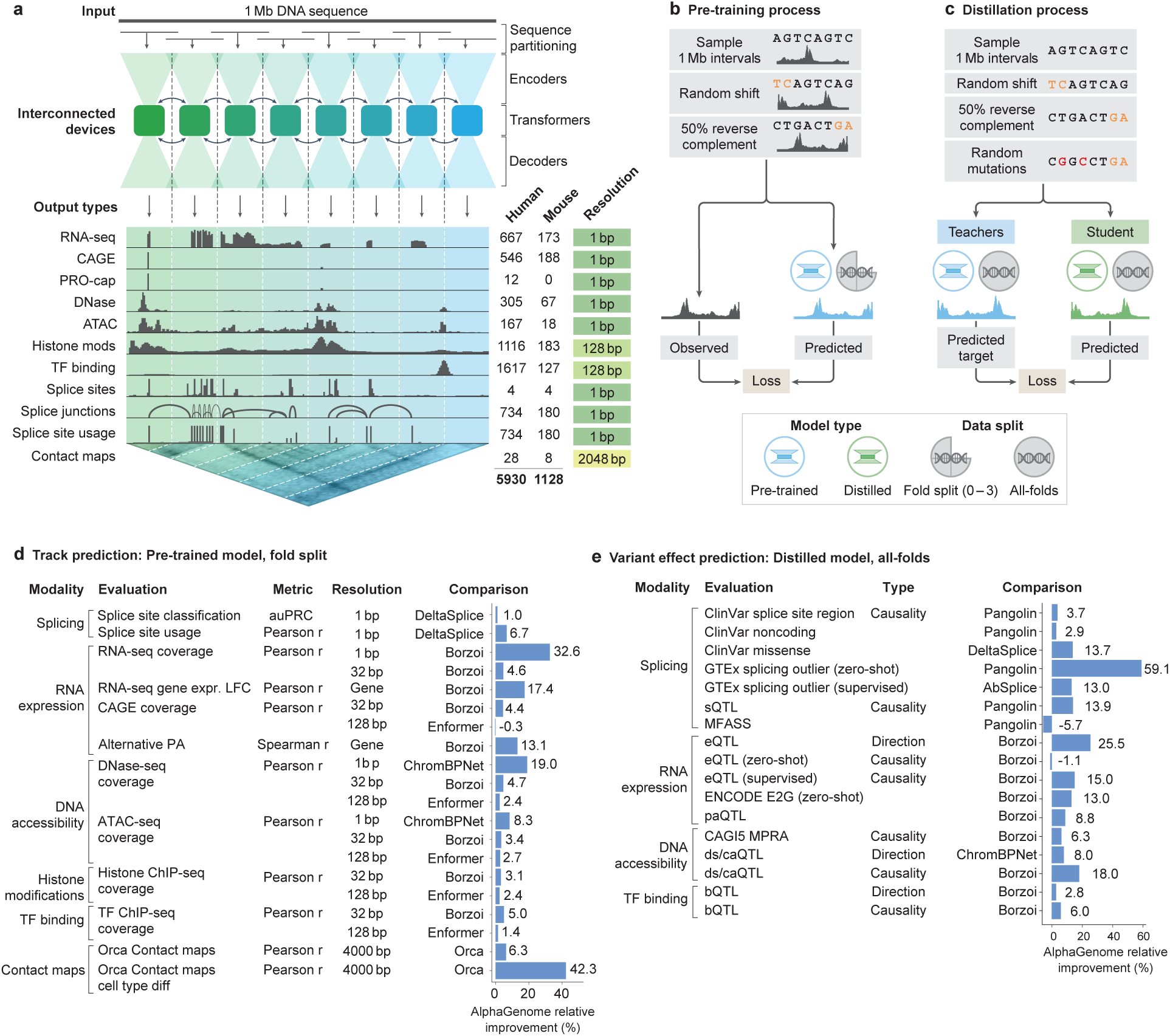
AlphaGenome model architecture, training regimes, and comprehensive evaluation performance. **(a)** Model overview. AlphaGenome processes 1 Mb DNA sequences and species identity (human/mouse) to predict 5,930 human or 1,128 mouse genome tracks across diverse cell types and 11 output types at specific resolutions (indicated far right). Computation leverages sequence parallelism, breaking the 1 Mb DNA sequence into 131 kb chunks processed across devices. The core architecture features a U-Net-style design comprising an encoder (downsampling the sequence), transformers with inter-device communication, and a decoder (upsampling), which feed into task-specific output heads responsible for generating the final predictions at their respective assay-specific resolutions (detailed in Extended Data Fig. 1). **(b)** 1 Mb DNA intervals are sampled from cross-validation folds, augmented (shifted and reverse complemented), and used to train the model against experimental targets, yielding fold-specific and *all-folds* teacher models. **(c)** A student model learns to reproduce predictions from frozen *all-folds* teacher models using augmented and mutationally perturbed input sequences, yielding a single model suitable for variant effect prediction. **(d)** Relative performance improvement (%) of AlphaGenome over the best competing model for a selection of genome track prediction tasks across different modalities and resolutions (Supplementary Table 3). For classification tasks, we adjust the relative improvement to account for the performance of a random classifier (Methods). PA: polyadenylation. **(e)** Relative performance improvement of AlphaGenome over the best competing model for a subset of variant effect prediction tasks (Supplementary Table 4). The ds/caQTL direction (causality) rows represent the average relative improvement across multiple similar datasets (Methods).

**Figure 2.**
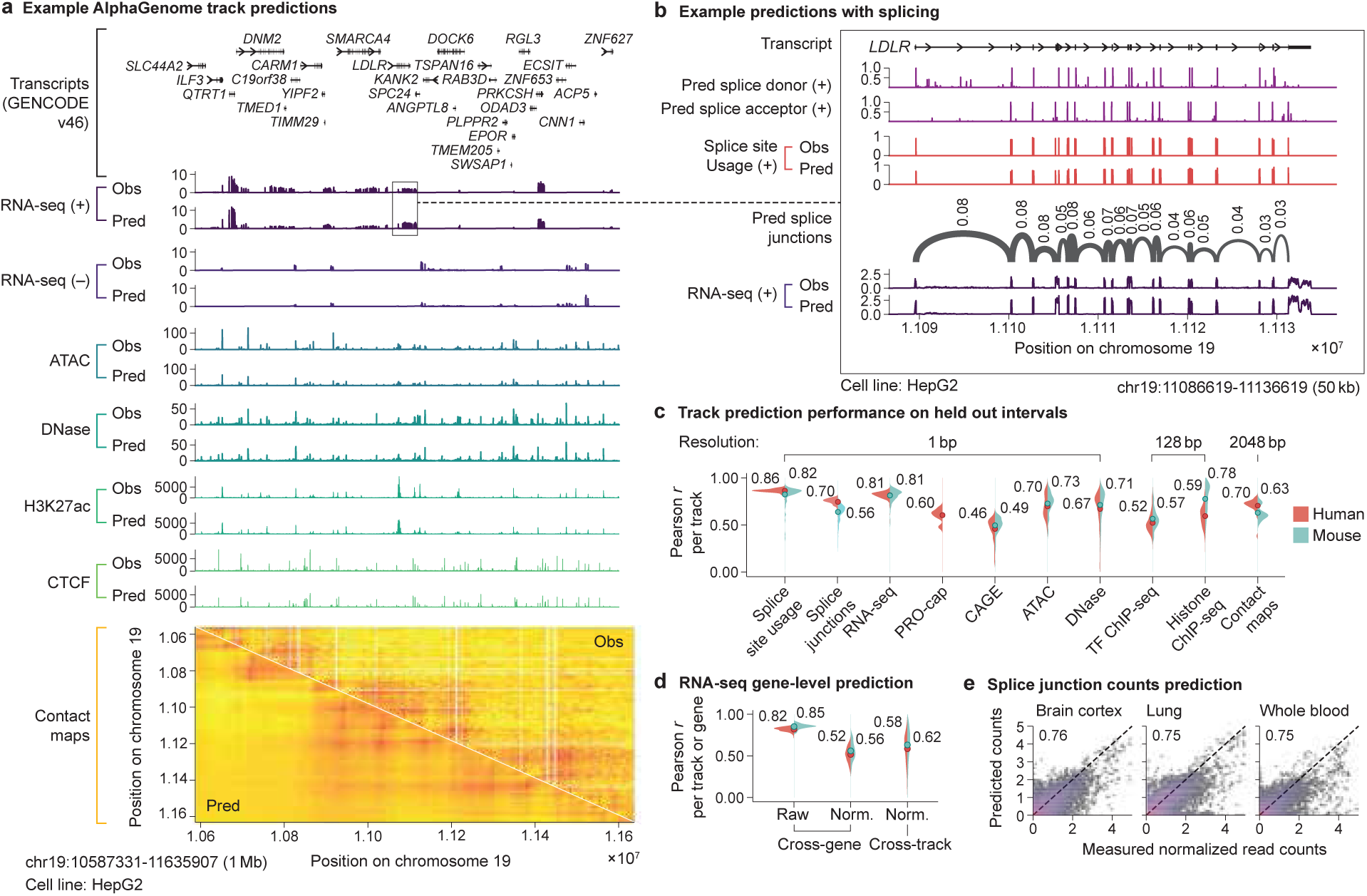
Example AlphaGenome track predictions and detailed performance evaluations. **(a)** Observed (Obs) and AlphaGenome- predicted (Pred) genome tracks within a 1 Mb held-out region of human chromosome 19 (0-based coordinates: 10587331-11635907) in the HepG2 cell line. Y-axis scales for each assay are defined in the Methods section. Strand-specific tracks are denoted as positive (+) or negative (-), while strand-agnostic tracks are shown without a strand symbol. Contact maps are pairwise interaction matrices, therefore both x- and y-axes display genome coordinate positions. RNA-, ATAC- and DNase-seq track predictions are at 1 bp resolution; H3K27ac and CTCF ChIP-seq are at 128 bp; and contact maps are at 2048 bp resolution. **(b)** Basepair resolution AlphaGenome predictions for a 50 kb (kilobase) region highlighting detailed splicing (donor/acceptor sites, splice site usage, splice junctions) and RNA-seq predictions around the *LDLR* gene. (**b**) is a zoomed-in view from (**a**). **(c)** Performance evaluation across different modalities. Violin plots display the distribution of Pearson correlations between predicted and observed tracks evaluated on held-out test intervals. Each violin plot is grouped by modality and split by organism (human in red, mouse in blue). Filled circles with accompanying numerical values indicate the mean Pearson *r* per assay group and organism. Splice junction, RNA-seq, PRO-cap, CAGE, and ChIP-seq tracks were log(1 + 𝑥) transformed, while the remainder were untransformed. **(d)** Evaluation of RNA-seq gene log-expression prediction on held-out test intervals. The left-most panel assesses Pearson correlation between predicted and observed log-expression values across all genes within individual tracks. The middle and right-most panels evaluate the prediction of tissue- or cell type-specificity using quantile-normalized expression values (detailed in Methods); correlations are computed either across genes per track (middle) or across tracks per gene (right). **(e)** Predicted versus observed splice junction read counts (log(1 + 𝑥) transformed, 𝑛 = 1, 344, 738) and Pearson r between them in selected human tissues known for having distinct splicing patterns^20^. Each hexagonal bin is colored by the density of the data points in that bin, with warmer colors corresponding to higher density. The diagonal dotted line indicates perfect agreement (predicted = observed). Additional tissues displayed in Extended Data Fig. 2d.

**Figure 3.**
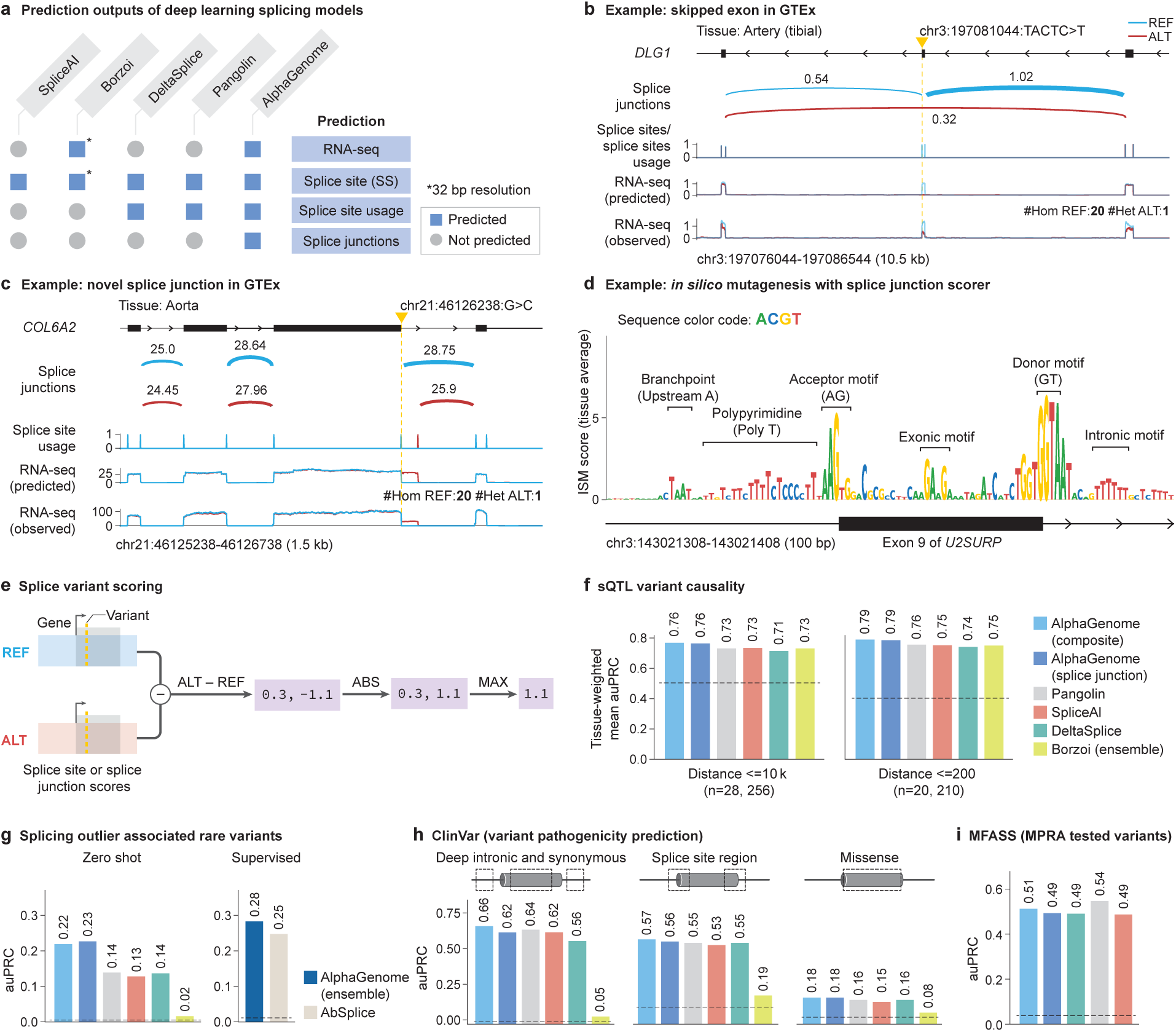
AlphaGenome is a state-of-the-art splicing variant effect prediction model. **(a)** Splicing prediction types across deep learning based models (SpliceAI^5^, Borzoi^3^, DeltaSplice^22^, Pangolin^11^, and AlphaGenome). All models predict at 1 bp resolution, with the exception of Borzoi at 32 bp resolution. All models produce explicit splicing predictions except Borzoi, whose splice site predictions are calculated implicitly through the 32 bp RNA-seq coverage outputs. **(b)** Example of a variant causing exon skipping in the *DLG1* gene. Predicted splice junction, splice site usage, and RNA-seq coverage, as well as the observed RNA-seq coverage for the reference allele (REF, blue) and the alternative allele (ALT, red) are shown. Predicted and observed tracks are from Artery (tibial) tissue. Observed RNA-seq coverage is obtained from GTEx by averaging samples with either the REF or ALT allele. The number of samples per genotype are annotated. #Hom REF: number of individuals homozygous for the REF sequence in GTEx. #Het ALT: number of individual heterozygous for the ALT sequence in GTEx . **(c)** Example of a variant causing alternative splice junction formation in the *COL6A2* gene by creating a new splicing donor and disrupting the extant one. Predicted and observed tracks are from Aorta tissue. **(d)** *In silico* mutagenesis of exon 9 of the *U2SURP* gene and the flanking introns using the mean splice junction score across tissues. Splicing related motifs are highlighted. **(e)** Schema of variant effect prediction with AlphaGenome. The maximum difference between REF and ALT predictions across splice sites or splice junctions is used to score variants (Methods). **(f)** Comparison of AlphaGenome composite and splice junction scorers versus other methods for classifying fine-mapped sQTL variants. Variants are stratified into two groups by distance to the splice site, as done in Borzoi^3^. Tissue-specific scores were used for Borzoi and AlphaGenome. Tissue-specific auPRCs were averaged, weighted by the variant count per tissue. **(g)** Comparison of AlphaGenome versus other methods for predicting whether a rare GTEx variant is associated with a splicing outlier event, as done in AbSplice^29^. AlphaGenome was evaluated in two settings: zero-shot (unsupervised) and supervised. In the latter, AlphaGenome’s scores were used to train an ensemble model, similar to the ensemble model AbSplice. Both ensemble models were trained/evaluated on the same data split of splicing outlier variants (Methods). **(h)** Classifying pathogenic versus benign ClinVar variants based on splicing effects for deep intronic (>6 bp from splice sites) and synonymous (>3 bp from splice sites) variants, variants in the splice site region (within 6 bp intronic or 3 bp exonic or), and missense variants predicted as ‘likely_benign’ by AlphaMissense^30^. **(i)** auPRC on the classification of experimentally validated splice disrupting variants (data from Chong *et al.*^28^). MFASS: Multiplexed Functional Assay of Splicing using Sort-seq. MPRA: Massively Parallel Reporter Assay.

**Figure 4.**
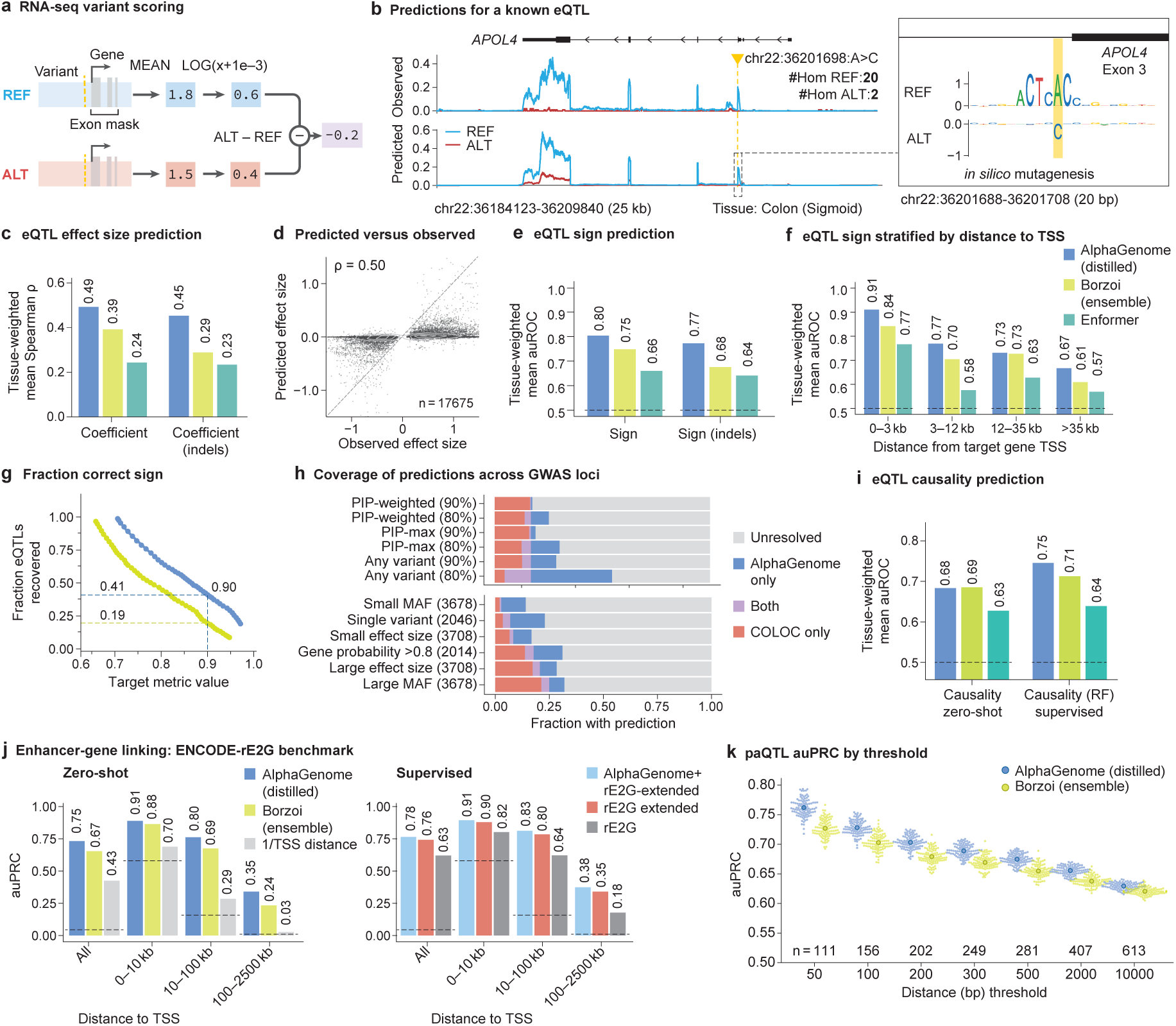
**AlphaGenome predicts the effect of variants on gene expression**. **(a)** Schematic of the variant scoring strategy summa- rizing the predicted effect of a genetic variant on the expression of a target gene (Methods). **(b)** Example predictions for a known eQTL (chr22:36201698:A>C) with SuSiE PIP > 0.9 in GTEx Colon (Sigmoid) tissue. The alternative ‘C’ allele is associated with lower expression (SuSiE ’beta posterior’ = -0.709), recapitulated in AlphaGenome’s variant score of -1.52 (quantile score -1.00; Methods) for the *APOL4* gene. Predicted and observed tracks are from Colon (Sigmoid) tissue. Observed RNA-seq coverage is obtained from GTEx by averaging samples with either the REF or ALT allele. The number of samples per genotype are annotated. #Hom REF: number of individuals homozygous for the REF sequence in GTEx. #Hom ALT: number of individuals for the ALT sequence in GTEx. *Inset:* Comparative *in silico* mutagenesis (ISM) on reference and alternative sequences over a 20 bp window centered on the variant (Methods). Lower values indicate that the allele shown is predicted to result in lower expression of *APOL4* relative to other alleles. **(c)** eQTL effect size (‘Coefficient’) prediction performance. Bar plot comparing tissue-weighted mean Spearman 𝜌 for AlphaGenome (distilled) against Borzoi (ensemble) and Enformer for predicting eQTL effect magnitudes for single nucleotide variants (SNVs; ‘Coefficient’) and insertions/deletions (‘Coefficient (indels)’) across 49 GTEx tissues. **(d)** Observed versus AlphaGenome-predicted eQTL effect sizes. Scatterplot comparing observed SuSiE ‘beta posterior’ values to AlphaGenome (distilled) predicted effect sizes for 17,675 fine-mapped GTEx eQTLs (SNVs). Each point is a unique variant/gene/tissue combination). Spearman’s 𝜌 (signed) = 0.50; Spearman’s 𝜌 (unsigned, absolute values) = 0.10. **(e)** eQTL direction of effect (‘Sign’) prediction performance. Bar plots comparing tissue-weighted mean Area Under the ROC Curve (auROC) for AlphaGenome (distilled) against Borzoi (ensemble) and Enformer for SNVs and indels. **(f)** eQTL Sign prediction performance stratified by variant-TSS distance for SNVs. Bar plots show tissue-weighted mean auROC per model across distance bins. **(g)** Sign prediction accuracy versus eQTL recovery. Plot relating target Sign accuracy (achieved by applying different AlphaGenome or Borzoi variant score thresholds) to the fraction of GTEx eQTLs recovered for AlphaGenome (distilled) versus Borzoi (ensemble). Dashed lines indicate eQTL recovery rates at score thresholds yielding 90% Sign accuracy for each model. **(h)** GWAS credible set direction of effect (Sign) prediction coverage. Fraction of GWAS loci (from Open Targets^36^) with a predicted direction of effect for a plausible target gene, comparing AlphaGenome (distilled) predictions at different accuracy thresholds (80% or 90%) against an eQTL colocalization approach (COLOC H4>0.95^35^, stratified by variant/locus properties (Methods). Bar segments indicate loci resolved by: COLOC only (blue), AlphaGenome only (red), both methods (purple), or neither (grey). **(i)** eQTL causality prediction (‘Causality’, tissue-weighted mean auROC). The bar plot compares zero-shot performance of AlphaGenome (distilled) against baselines and supervised performance using a Random Forest on model variant scores (‘Causality (RF)’). **(j)** Enhancer-gene linking performance (ENCODE-rE2G CRISPRi dataset^17^). Zero-shot evaluation: Performance (auPRC) comparison stratified by enhancer-TSS distance for AlphaGenome (distilled) vs Borzoi vs TSS distance baseline. Supervised evaluation: AlphaGenome input gradient score integrated into ENCODE-rE2G-extended vs ENCODE-rE2G models. **(k)** Performance of paQTL variant effect prediction for Borzoi and AlphaGenome, thresholded by distance from the polyadenylation site. AlphaGenome outperforms Borzoi at all the considered distances, although auPRC declines for both models as more distal paQTLs are included. Each swarmplot represents the values obtained from 100 permutations of randomly matching each positive SNP with one of its distance and expression-matched negatives (Methods) and computing the auPRC, with the overlaid dot representing the mean value.

**Figure 5.**
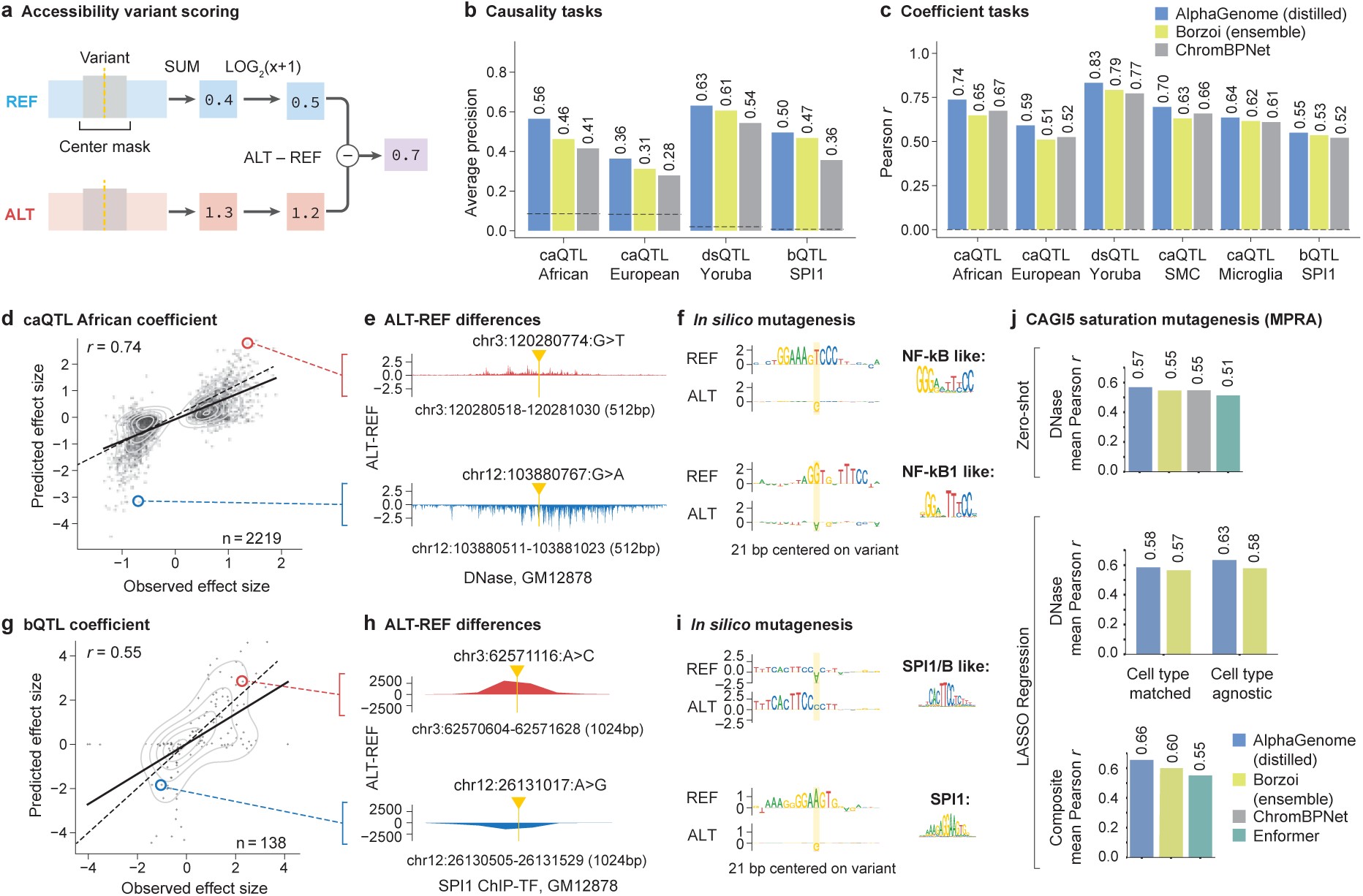
AlphaGenome accurately predicts variant effects on chromatin accessibility and SPI1 transcription factor binding. **(a)** Schematic of the center-mask variant scoring strategy. This approach, detailed in Methods, is used for accessibility (DNase-seq, ATAC-seq) and ChIP-seq predictions. **(b)** Performance comparison on QTL causality prediction. Average Precision (AP) for AlphaGenome, Borzoi, and ChromBPNet across QTL types (caQTL, dsQTL, bQTL) and ancestries. **(c)** Performance comparison on QTL effect size prediction. Pearson *r* is shown for AlphaGenome, Borzoi, and ChromBPNet across QTL types (caQTL, dsQTL, bQTL) and ancestries. **(d)** AlphaGenome’s predicted versus observed effect sizes for causal caQTLs (African ancestry). Scatterplot displays predictions using the DNase track for the GM12878 cell line. Signed Pearson *r* = 0.74; unsigned Pearson *r* = 0.45. Signed Pearson *r* correlation uses raw values; unsigned Pearson *r* uses absolute values. Red and blue circles highlight variants detailed in **(e, f)**. **(e)** Example AlphaGenome predictions for selected caQTLs. Shown are ALT-REF differences in predicted DNase track (GM12878) around the variants highlighted in **(d)**. **(f)** ISM-derived sequence logos for REF and ALT alleles of example caQTLs from **(e).** The examples suggest variant disruption or modulation of TF binding motifs. Putative binding factors and JASPAR^38^ matrix IDs (MA0105.1, MA0105.3) are indicated on the right. **(g)** AlphaGenome’s predicted versus observed effect sizes for causal SPI1 bQTLs. Scatterplot displays predictions using the SPI1 ChIP-seq track for the GM12878 cell line. Signed Pearson *r* = 0.55; unsigned Pearson *r* = 0.12. Red and blue circles highlight variants detailed in **(h, i)**. **(h)** Example AlphaGenome predictions for selected SPI1 bQTLs. Shown are ALT-REF differences in predicted SPI1 ChIP-TF track (GM12878) around the variants highlighted in **(g)**. **(i)** ISM-derived sequence logos for REF and ALT alleles of example SPI1 bQTLs from **(h)**. Examples indicate potential motif impacts such as creation or disruption of SPI1 or related motifs. Putative binding factors and JASPAR matrix IDs (MA0081.2, MA0080.5) are indicated on the right. **(j)** CAGI5 MPRA challenge performance (average across loci). **Top:** Average zero-shot Pearson *r* performance, using cell type-matched raw DNase model outputs. **Middle:** Average Pearson *r* from LASSO regression using cell type-matched or cell type-agnostic DNase outputs. **Bottom:** LASSO regression Pearson *r* performance using features from multiple modalities and the full set of cell types (DNase + RNA + ChIP-Histone output types for AlphaGenome and Borzoi; DNase + CAGE output types for Enformer).

**Figure 6.**
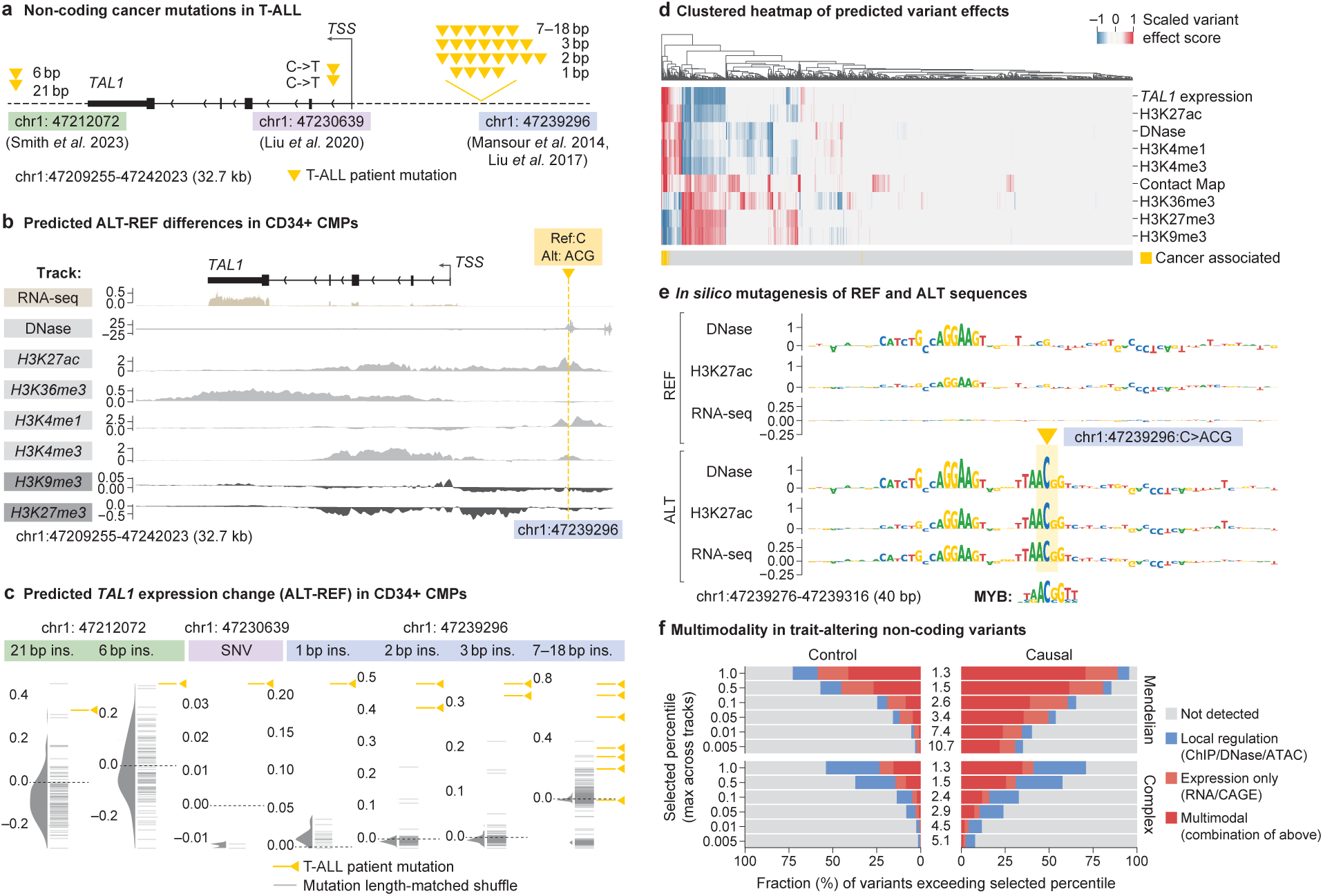
Interpreting variant effects across modalities with AlphaGenome. **(a)** Overview of groups of mutations affecting *TAL1* in T-ALL patients. **(b)** Detailed ALT-REF predictions for an oncogenic insertion (chr1:47239296:C>ACG) characterized in Mansour et al. 2014^42^. Shown are differences between AlphaGenome predictions between the ALT sequence and the REF sequence of the variant in CD34+ CMP tracks. The ALT sequence increases expression of the *TAL1* gene 7.5kb away. **(c)** RNA-seq variant scores for TAL1 expression in CD34+ CMPs. Oncogenic mutations (orange) are compared against randomly sampled, length-matched indels (gray). **(d)** Multimodal heatmap of variant effects. Each column is a distinct variant from **(c)**. Each row is a variant effect score associated with a genome track in CD34+ CMPs, except for contact map variant effect scores, which were averaged across tissues (as there is no CD34+ CMP contact map in our data). Background mutations are included alongside oncogenic mutations. Variants were grouped by their insertion length and position (as displayed in Fig. 6c) and their scores were min-max scaled. **(e)** ISM results for DNase, H3K27ac and TAL1 RNA-seq expression prediction by AlphaGenome in CD34+ CMPs. Top: ISM on the reference sequence. Bottom: ISM on the oncogenic insertion sequence (chr1:47239296:C>ACG). Myb motif from the Mansour et al 2014 study^42^, originally from UniProbe^46^. **(f)** Fraction of trait-affecting variants^47^ (338 for mendelian and 1140 for complex traits), as well as matched control variants^47^ (3042 and 10260, respectively), which exceed varying quantile-score thresholds in at least one predicted track. Variants are categorized depending on the tracks where the threshold was passed: ‘Local Regulation’ (ChIP/DNase/ATAC), ‘Expression only’ (RNA/CAGE) and ‘Multimodal’ (combination of the above). Numbers in the center indicate the relative enrichment of detected variants among causal variant candidates.

**Figure 7.**
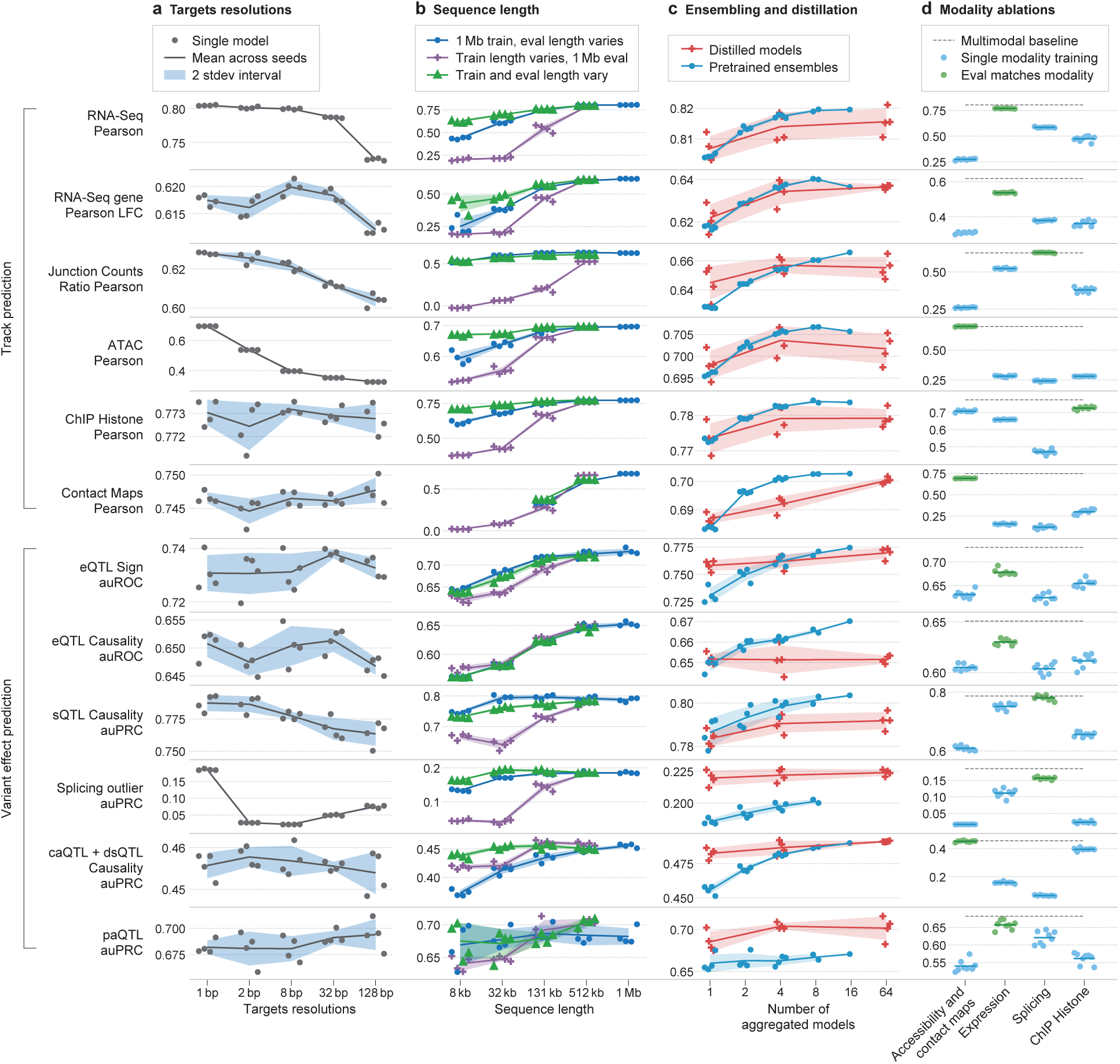
Impact of resolution, sequence length, ensembling, distillation, and multimodal training on AlphaGenome performance. Ablation studies evaluating key model design choices across various performance metrics (y-axis). For all panels, lines represent the mean over replicate training runs with different random seeds (𝑛 = 4 unless otherwise stated), and shaded contours denote the uncertainty interval (two standard deviations). **(a)** Impact of target resolution. Performance comparison across models trained to predict targets (e.g., DNA accessibility, gene expression, splicing) at varying resolutions (x-axis, 1 bp to 128 bp). stdev: Standard deviation. **(b)** Impact of sequence length during training and inference. Blue dots represent a single set of models trained with 1 Mb input, evaluated using varying input sequence lengths (x-axis). Purple crosses represent models trained at the sequence length indicated on the x-axis but evaluated at a fixed 1 Mb input length. Green triangles represent models trained and evaluated using the same matched sequence length (x-axis). **(c**) Impact of the number of sub-models in ensembling and distillation. Performance comparison for mean ensembles of pre-trained models (blue dots/contours, x-axis indicates ensemble size) versus single models produced by distillation using 1, 4, or 64 teacher models (orange crosses/contours, x-axis indicates number of teachers). (**d)** Impact of multimodal learning. Performance comparison evaluating models trained only on specific modality groups (blue dots, 𝑛 = 8 seeds per group, highlighted in green if the modality matches the evaluation metric) against the full multimodal model (black dashed line, 𝑛 = 4 seeds average). During training for these models, we ensured that only the target modality group’s prediction heads contributed updates to the shared representations, allowing assessment of that modality group’s contribution to overall model performance. Groups shown (x-axis) include models trained using gradients only from Accessibility (ATAC, DNase, Contact Maps), Expression (RNA-Seq, CAGE, PRO-cap), Splicing (sites, usage, junctions), or Histone ChIP-seq.

## Results

### A unifying DNA sequence-to-function model

AlphaGenome is a deep learning model designed to learn the sequence basis of diverse molecular phenotypes from human and mouse DNA (Fig. 1a). It simultaneously predicts 5,930 human or 1,128 mouse genome tracks across 11 modalities covering gene expression (RNA-seq, CAGE, PRO-cap), detailed splicing patterns (splice sites, splice site usage, splice junctions), chromatin state (DNase, ATAC-seq, histone modifications, TF binding), and chromatin contact maps. These span a variety of biological contexts, such as different tissue types, cell types and cell lines (see Supplementary Table 1 for summary and Supplementary Table 2 for complete metadata). These predictions are made on the basis of 1 Mb of DNA sequence, a context length designed to encompass a substantial portion of the relevant distal regulatory landscape; for instance, 99% (465/471) of validated enhancer-gene pairs fall within 1 Mb^17^.

AlphaGenome employs a U-Net-inspired^3,18^ backbone architecture (Fig. 1a; Extended Data Fig. 1) to efficiently process input sequences into two types of sequence representations: 1-dimensional embeddings (at 1 bp and 128 bp resolutions) that correspond to representations of the linear genome, and 2-dimensional embeddings (2048 bp resolution) that correspond to representations of spatial interactions between genomic segments. The 1-dimensional embeddings serve as the basis for genomic track predictions, while the 2-dimensional embeddings are the basis for predicting pairwise interactions (contact maps). Within the architecture, convolutional layers model local sequence patterns necessary for fine-grained predictions, while transformer blocks model coarser but longer-range dependencies in the sequence such as enhancer-promoter interactions. Basepair resolution training on the full 1 Mb sequence is enabled by sequence parallelism across 8 interconnected Tensor Processing Unit (TPUv3) devices. Genomic track predictions are linear transformations of these sequence embeddings, aside from splice junction count prediction, which uses a separate mechanism that captures interactions between 1-dimensional embeddings of donor/acceptor pairs (Extended Data Fig. 1).

We trained the model using a two-stage process: pre-training and distillation. The pre-training phase (Fig. 1b) utilizes the observed experimental data to produce two types of models. *Fold-specific* models are trained using a 4-fold cross-validation scheme (Methods), training on 3/4 of the reference genome while holding out the remaining 1/4 for validation and testing, and are used to evaluate AlphaGenome’s generalization by predicting genomic tracks on unseen (test) reference genome intervals (Fig. 1b). Additionally, *all-folds* models are trained on all available intervals of the reference genome, and serve as teachers for the second stage: distillation (Fig. 1c). In the distillation phase, a single *student* model, sharing the pre-trained architecture, is trained to predict the output of an ensemble of *all-folds* teachers using randomly augmented input sequences (Methods). This distilled student model, as shown previously^19^, achieves improved robustness and variant effect prediction (VEP) accuracy in a single model instance, making predictions across all modeled modalities and cell types with a single device call per variant. Taking less than one second on a NVIDIA H100 GPU, the student model is highly efficient for large-scale variant effect prediction relative to the alternative approach of ensembling several independently trained models.

### Evaluating AlphaGenome on genome track and variant effect prediction tasks

To characterize AlphaGenome’s model performance, we first assessed its generalization to unseen genome intervals, a prerequisite for high-quality variant effect prediction. We conducted 24 genome track evaluations, encompassing all 11 predicted modalities (Methods, Supplementary Table 3). For out-of-fold evaluations, pre-trained, fold-specific AlphaGenome models were used and compared against the strongest available external model for each respective task. AlphaGenome outperformed these external models on 22 out of 24 evaluations (Fig. 1d; Extended Data Fig. 3, Supplementary Table 3). Notably, AlphaGenome exhibited a +17.4% relative improvement in cell type-specific gene-level expression log-fold change (LFC) prediction compared to Borzoi^3^, another multimodal sequence model (Fig. 1e, stratified metrics in Extended Data Fig. 3e). AlphaGenome also outperformed specialized single-modality models on their respective tasks, such as Orca^4^ on contact maps (contact map Pearson *r* +6.3%, cell type-specific differences +42.3%, Fig. 1d; Extended Data Fig. 4), ProCapNet^8^ on transcription initiation tracks (+15% total counts Pearson *r*, Extended Data Fig. 3f), and ChromBPNet^10^ on accessibility (+8% for ATAC, +19% for DNase on total counts Pearson *r*, Extended Data Fig. 3g).

We next evaluated the model’s performance on predicting variant effects. We assembled a second set of 26 variant effect prediction benchmarks across gene expression, splicing, polyadenylation, enhancer-gene linking, DNA accessibility and TF binding. Again, we compared against the strongest externally available model on each task (Methods, Supplementary Table 4). For variant effect prediction, we used the distilled student model. AlphaGenome matched or outperformed the external models on 24 out of 26 evaluations (Fig. 1e, Supplementary Table 4). This included strong performance in quantitative trait loci (QTL) evaluations, such as sign prediction for expression QTLs (+25.5% versus Borzoi^3^) and accessibility QTLs (+8.0% versus ChromBPNet^10^, average across five datasets; Methods), demonstrating its strength against both multimodal and specialized single-modality baselines. Collectively, these results demonstrate that AlphaGenome more accurately models both genome tracks and variant effects.

### AlphaGenome achieves state-of-the-art genome track prediction across modalities

Given AlphaGenome’s strong performance on genome track evaluations, we investigated its track predictions in more detail. Fold-specific, pre-trained AlphaGenome models demonstrated high concordance between predicted and observed read coverage on unseen genome intervals (Fig. 2). As an example, predicted HepG2 genome tracks over the *LDLR* gene showcased strand-specific, basepair-resolution RNA-seq coverage over exons, along with predicted splice sites, splice site usage and splice junction read coverage (Fig. 2b). Additional examples illustrating splicing, gene expression, and chromatin track predictions are provided in Supplementary Figs. 1 to 3, and finer delineation of genomic features such as exon boundaries is highlighted in Supplementary Fig. 4.

Quantitatively, we observed strong Pearson correlations (*r* ) between predicted and observed signals for functional genomics tracks in both human and mouse genomes (Fig. 2c), both across all tracks and when subsetting by biosample types or data sources (Supplementary Fig. 5). While overall expression levels are predicted well, accurately capturing cell type-specific expression deviations remains a challenging task (Fig. 2d, Supplementary Fig. 2j).

On splicing modalities (Extended Data Fig. 2a), AlphaGenome accurately predicts splice sites (Extended Data Fig. 2b) and splice site usage (Extended Data Fig. 2b,c). It also accurately predicts quantitative splice junction read coverage, PSI5, and PSI3 within various tissues, achieving high correlation with experimental measurements (Fig. 2e; Extended Data Fig. 2b,d,e; Methods). While AlphaGenome demonstrates the ability to predict tissue-specific alternative splicing in some instances (Supplementary Fig. 1), further improvements are needed to precisely predict intermediate splicing efficiencies and to capture tissue-specific nuances (Extended Data Fig. 2c,e).

### AlphaGenome is a state-of-the-art splicing variant effect prediction model

One of the major ways genetic variants cause disease is by disrupting splicing^21^, a process that produces mature RNA sequences by excising introns and ligating exons at splice junctions. Splicing outcomes can be modeled at three levels: the probability that any given nucleotide acts as a splice donor or acceptor (splice site prediction)^5,11,22^, the competitive selection among potential splice sites (splice site usage prediction)^11,22^, and the prediction of specific introns (splice junction prediction). AlphaGenome is the first system to predict all three of these quantities alongside direct RNA-seq coverage prediction, thereby providing a more comprehensive view of the splicing-related molecular consequences of variants (Fig. 3a).

To illustrate the distilled AlphaGenome’s capacity to simultaneously predict multiple relevant splicing variant effects, we first probed its ability to recapitulate known biological outcomes. We interrogated a 4 bp deletion (chr3:197081044:TACTC>T), a variant empirically observed to cause exon skipping in tibial artery tissue in a GTEx^23^ sample (Fig. 3b). AlphaGenome accurately predicted this established consequence across all levels: a substantial reduction in the predicted usage of the affected exon’s splice site, the loss of predicted junctions linking the skipped exon edges, the emergence of a putative junction bypassing the exon, and a strong decrease in predicted RNA-seq coverage of the exon. Similarly, AlphaGenome’s predictions accurately captured the novel splice junction and extended exon induced by the chr21:46126238:G>C variant, an effect observed in a heterozygous GTEx RNA-seq sample (Fig. 3c). Finally, *in silico* mutagenesis (ISM), which systematically predicts effects of all possible single nucleotide variations in a sequence region (Methods), revealed the sequence determinants of the splicing predictions. For example, ISM analysis of exon 9 of the gene *U2SURP* and its flanking introns highlighted recognizable splicing-related sequence motifs^24,25^ (Fig. 3d). Further examples of experimentally validated splice disrupting variants identified in individuals with autism spectrum disorders^5^ are shown in Supplementary Fig. 6.

Building on AlphaGenome’s multifaceted splicing predictions, we developed a unified splicing variant scorer to systematically detect splicing-disrupting variants. Specifically, we designed a custom variant scoring strategy for each prediction modality (Fig. 3e, Methods) and summed the individual scores to provide a composite measure of a variant’s predicted effect. We benchmarked this composite scorer against existing methods on a wide range of splicing-related VEP tasks: AlphaGenome performed best on fine-mapped splicing QTL (sQTL) classification^3,26^, including both single nucleotide polymorphisms (SNPs) within 10kb of the closest splice site, and proximal variants within 200 bp of splice sites (Fig. 3f); furthermore, it achieved the highest performance at predicting rare SNPs and indels associated with GTEx splicing outliers in both supervised and unsupervised settings (Fig. 3g, Methods). We also evaluated AlphaGenome’s performance at distinguishing pathogenic variants from benign variants in ClinVar^27^, where its composite scores outperformed the existing best method in each category across all three variant categories: deep intronic and synonymous (area under the precision-recall curve, auPRC, 0.66 vs. 0.64 by Pangolin), splice region (auPRC 0.57 vs. 0.55 by Pangolin), and missense (auPRC 0.18 vs. 0.16 by DeltaSplice and Pangolin; Fig. 3h, Supplementary Fig. 7). Conversely, when assessing AlphaGenome’s ability to predict if rare variants disrupt splicing using data from a massively parallel splicing minigene reporter assay (MFASS)^28^, it was outperformed by Pangolin (auPRC 0.54 vs. 0.51) but surpassed SpliceAI and DeltaSplice (both auPRC 0.49; Fig. 3i). Notably, the splice junction scorer alone outperformed previous approaches on all benchmarks except “deep intronic and synonymous” ClinVar variants and MFASS variants, underscoring the importance of modeling splicing at the junction level. In summary, AlphaGenome achieves SOTA splicing variant effect prediction on 6 out of 7 benchmarks, providing a more comprehensive view of altered splicing events and transcript structure.

### AlphaGenome improves performance across gene expression regulation tasks

Beyond impacting isoform composition via splicing modulation, non-coding variants can significantly influence traits and cause diseases by altering gene expression^31,32^. We evaluated AlphaGenome’s ability to predict the impact of variants on gene expression across a range of regulatory mechanisms, including expression quantitative trait loci (eQTLs), enhancer-gene interactions, and alternative polyadenylation (APA).

### Improved prediction of eQTL effects

We first evaluated AlphaGenome’s ability to predict the impact of eQTLs, which are variants associated with gene expression variation. We developed a variant scoring strategy to quantify a variant’s predicted effect on a gene’s expression level (Fig. 4a; Methods). An example at a known eQTL/sQTL locus (rs9610445, chr22:36201698:A>C, posterior inclusion probability (PIP)=0.90) illustrates the model’s predictions aligning with observed allele-specific expression differences from GTEx data (Fig. 4b). Furthermore, ISM around the variant suggests a possible mechanism: disruption of a predicted splicing motif by the alternative ‘C’ allele, potentially leading to an aberrant transcript (Fig. 4b, inset). Additional examples of eQTL variant prediction and mechanistic interpretation by ISM are shown in Supplementary Fig. 8.

Using fine-mapped GTEx eQTLs as ground truth, we benchmarked AlphaGenome against the SOTA models Borzoi^3^ and Enformer^2^. AlphaGenome demonstrated significantly improved prediction of both the magnitude (‘Coefficient’, Spearman 𝜌 with SuSiE^33^ ‘beta posterior’) and direction (‘Sign’, area under the receiver operating curve, auROC) of eQTL effects compared to the previous SOTA, Borzoi (Fig. 4c-f). AlphaGenome improves tissue-weighted mean Spearman 𝜌 from 0.39 to 0.49 and mean Sign auROC from 0.75 to 0.80. These improvements in Coefficient and Sign prediction were observed broadly across the majority of GTEx tissues, variant-to-TSS distance bins, and variant functional annotation classes (Extended Data Fig. 5a-c).

As previously reported^34^, performance decays with distance to the target gene across all eQTL tasks (Extended Data Fig. 5f); however, AlphaGenome exhibits mild improvement on Sign/Coefficient prediction for distal variants (>35kb, Fig. 4f, Extended Data Fig. 5b), and top-scoring predictions exhibit high sign accuracy across distance categories (Extended Data Fig. 5g). Additionally, AlphaGenome outperforms Borzoi on Coefficient and Sign prediction for indel (insertion or deletion) eQTLs (Fig. 4c,e, Extended Data Fig. 5d). Notably, AlphaGenome’s effect size predictions for high-confidence eQTLs (scores >99th percentile of common variant effects) highly correlate with observed effects (Spearman 𝜌 = 0.73; Extended Data Fig. 5h) and consistently surpass Borzoi’s performance across various quantile score thresholds (e.g., Spearman 𝜌 0.73 vs. 0.61 at the ∼99th percentile threshold; Extended Data Fig. 5i).

The overall improvement in prediction accuracy, particularly for the sign of a variant’s effect, translates to substantial gains in practical applications. At a score threshold yielding 90% Sign prediction accuracy, AlphaGenome recovers over twice as many GTEx eQTLs (41%) as Borzoi (19%; Fig. 4g). Applying this improved Sign prediction capability to GWAS interpretation, we evaluated AlphaGenome’s ability to assign a direction of effect to candidate target genes for 18,537 GWAS credible sets (Methods). Using a threshold calibrated to 80% accuracy on eQTLs (Fig. 4g), AlphaGenome assigned a confident sign prediction for at least one variant in 49% of GWAS credible sets (11% using a conservative PIP-weighted scoring approach; Fig. 4h). AlphaGenome and a widely used colocalization method for sign prediction (COLOC H4>0.95)^35^ resolve the direction of effect for largely non-overlapping sets of loci (Fig. 4h). This indicates their complementary utility, collectively increasing the total yield of loci with determined effect directions. Furthermore, AlphaGenome resolved ∼4-fold more credible sets in the lowest minor allele frequency (MAF) quintile compared to COLOC, likely reflecting its reduced dependence on population genetics parameters that affect power to detect associations (Fig. 4h, stratified bars). Thus, AlphaGenome expands our ability to generate functional hypotheses regarding the direction of GWAS signals, particularly for low-frequency variants.

AlphaGenome’s performance on distinguishing fine-mapped eQTLs from distance-matched variants (‘Causality’, auROC) was comparable to Borzoi (Fig. 4i). However, leveraging AlphaGenome’s predictions in a supervised framework significantly boosted performance on the causality task: training a random forest model using AlphaGenome’s scores from multiple modalities increased the mean auROC from 0.68 to 0.75, also surpassing previous SOTA performance (mean auROC 0.71 for Borzoi, Fig. 4i). Notably, using features derived from variant scores across *all* predicted modalities provided a significant performance uplift in this supervised setting compared to using RNA-seq derived scores alone (Extended Data Fig. 5e), highlighting the practical benefit of AlphaGenome’s multimodal predictions for identifying causal expression-modulating variants.

### AlphaGenome enables state-of-the-art enhancer-gene linking

We then assessed whether AlphaGenome can link enhancer elements to their target genes, given that tissue-specific gene expression is modulated by enhancer-promoter (E-P) interactions, often involving enhancers in distal genomic regions. For this task, we leveraged an independent CRISPRi perturbation dataset from the ENCODE-rE2G study^17^. Evaluated zero-shot, AlphaGenome outperformed Borzoi in identifying validated E-P links, particularly for enhancers located beyond 10kb from the target gene’s TSS (Fig. 4j, Extended Data Fig. 7a; Methods), although both models still somewhat underestimate the impact of very distal enhancers (Extended Data Fig. 7d). Furthermore, AlphaGenome’s zero-shot performance was comparable (within 1% auPRC) to the ENCODE-rE2G-extended model which was explicitly trained on this task and cell line data (Fig. 4j). It also strongly outperformed the simpler DNase-based ENCODE- rE2G model and a distance-to-TSS baseline. Beyond their standalone predictive power, features derived from AlphaGenome enhanced supervised E-P linking models. Incorporating AlphaGenome predictions into the ENCODE-rE2G-extended model yielded a new SOTA performance across all distance-to-TSS categories (Fig. 4j, Extended Data Fig. 7b-c; Methods).

Altogether, these results together demonstrate AlphaGenome’s improved capacity to capture long-range functional regulatory connections directly from sequence, which is vital for interpreting distal genetic variants.

### Accurate prediction of alternative polyadenylation and paQTLs

We further evaluated AlphaGenome on variants affecting alternative polyadenylation (APA). APA is a process that generates transcript diversity by varying the 3’ untranslated region (3’ UTR) of mRNA molecules, often with concomitant impacts on RNA half-life and tissue specificity^37^. By predicting RNA-seq coverage, AlphaGenome also intrinsically models the competition between proximal and distal polyadenylation signals (PASs) and thus can be used to detect alternative polyadenylation. We found that AlphaGenome achieves SOTA performance in predicting APA (Spearman 𝜌 = 0.894 vs. Borzoi’s 0.790; Extended Data Fig. 6a; Methods)^3^. Given this, we next examined its ability to distinguish 3’ polyadenylation QTLs (paQTLs) from expression-matched negatives (Fig. 4k; Methods). AlphaGenome outperformed Borzoi on this task (all paQTLs within 10,000 bp of PAS: auPRC 0.629 vs. Borzoi’s 0.621; proximal paQTLs within 50bp of PAS: auPRC 0.762 vs Borzoi’s 0.727), although the predictive accuracy of both models decayed with distance to the polyadenylation site (Fig. 4k, Supplementary Table 5). *In silico* mutagenesis revealed that AlphaGenome learned the relevance of the canonical polyadenylation motif and could detect variants which disrupt or create this motif (Extended Data Fig. 6c- h), although it occasionally underpredicted the expression impact (Extended Data Fig. 6i-j). Overall, these results demonstrate AlphaGenome’s improved accuracy in predicting genetic variant effects on 3’ UTR processing – a distinct regulatory mechanism – using only RNA-seq predictions and without any explicit training on polyadenylation data or associated variants.

### AlphaGenome improves on predicting variant effects on chromatin accessibility and transcription factor binding

We next assessed AlphaGenome’s ability to predict the effects of variants on chromatin states, focusing on QTLs associated with chromatin accessibility (caQTLs), DNase sensitivity (dsQTLs), and transcription factor binding (bQTLs). Variant effects were scored by comparing model predictions for reference and alternative alleles in a local window around the variant (Methods; Fig. 5a). Benchmarking performance using fine-mapped QTLs from diverse ancestries (using benchmarks from ChromBPNet^10^), AlphaGenome consistently achieved SOTA performance compared to both Borzoi and the specialized ChromBPNet model. This was observed across QTL types and ancestries for both predicting QTL causality (Average Precision, Fig. 5b, Supplementary Fig. 9) and correlating with experimentally determined effect sizes (‘Coefficient’, Pearson r, Fig. 5c, Supplementary Fig. 9). This strong performance generalized across additional datasets, including European ancestry caQTLs, Yoruba ancestry dsQTLs, and predictions within specific cell types like microglia and cardiac smooth muscle cells (SMC). To illustrate this performance and gain mechanistic insight, we examined specific examples including fine-mapped caQTLs (African ancestry) and SPI1 bQTLs^39^. In both cases, AlphaGenome outperformed alternative models in causality prediction (precision-recall curves, Supplementary Fig. 9e, h) and its predicted effect sizes correlated with observed values (Fig. 5d,g; Pearson r=0.74 for caQTLs, r=0.55 for SPI1 bQTLs). Furthermore, ISM applied to high-impact variants revealed that predicted changes in accessibility or TF binding scores generally corresponded to altered motifs for transcription factors known to modulate chromatin accessibility, such as NF-𝜅B (Fig. 5e,f). For SPI1-specific variants, ISM further highlighted altered SPI1 motifs within the local sequence context (Fig. 5h,i).

Beyond population-level QTLs, understanding the impact of local sequence context on gene regulation is crucial. We therefore assessed AlphaGenome’s performance on the CAGI5 saturation mutagenesis MPRA challenge^40,41^ (Supplementary Note: AlphaGenome shows competitive performance on the CAGI5 MPRA benchmark). MPRAs measure the regulatory activity of short DNA sequences (typically via reporter gene expression), a process that is closely linked to local chromatin accessibility and transcription factor binding. We therefore evaluated this benchmark using DNase, RNA-seq, and ChIP output types, comparing against Enformer, Borzoi, and ChromBPNet (Methods; Extended Data Fig. 8). Notably, AlphaGenome’s cell type-matched DNase predictions achieved performance comparable to both ChromBPNet and the Borzoi Ensemble (Fig. 5j, top; Pearson *r* =0.57). Furthermore, aggregating features from DNase predictions across all modeled cell types using LASSO regression further improved performance over cell type-matched DNase features and over the analogous approach with Borzoi Ensemble (Fig. 5j, middle; Pearson *r* =0.63). Finally, by integrating features across multiple modalities and all cell types with LASSO, AlphaGenome achieved SOTA performance on the CAGI5 benchmark (Fig. 5j, bottom; Pearson *r* =0.65). Overall, these analyses highlight AlphaGenome’s strong performance in predicting variant effects on chromatin states in QTL benchmarks, as well as its ability to leverage accessibility predictions to model the regulatory activity of sequences as measured by MPRAs.

### AlphaGenome enables a multimodal analysis of variants

We next interrogated how AlphaGenome’s unified modeling approach can be used to virtually screen a locus. We evaluated three groups of gain-of-function mutations affecting the *TAL1* gene in T-cell acute lymphoblastic leukemia (T-ALL): a cluster of 5’ neo-enhancer mutations upstream of the *TAL1* TSS^42,43^, an intronic single nucleotide variant^44^, and a 3’ neo-enhancer^45^. All three variant groups were shown to converge on a common mechanism – upregulation of the *TAL1* oncogene (Fig. 6a,b).

To assess AlphaGenome, we analyzed predictions in CD34+ common myeloid progenitor (CD34+ CMP) data, the closest available match to T-ALL cell of origin. For the oncogenic mutation chr1:47239296: C>ACG^42^, AlphaGenome predicted increases in activating histone marks H3K27ac, H3K4me1 and H3K4me3 at the variant, consistent with experimentally observed neo-enhancer formation at that position^42,43^ (Fig. 6b). It also predicted decreased levels of the repressive histone marks H3K9me3 and H3K27me3 near the *TAL1* TSS, and elevated active transcription mark H3K36me3 across the *TAL1* gene body, both of which are concordant with its predicted increase in *TAL1* mRNA expression levels (Fig. 6b).

We expanded our analysis to include all oncogenic variants across studies^42–45^, comparing each variant to a background set of length-matched and sequenced shuffled control variants (for the sole intronic SNV, we compared it to other possible SNVs at that site). Oncogenic mutations were predicted to increase *TAL1* expression in CD34+ CMPs more than shuffled controls (Fig. 6c). Across modalities, oncogenic variants exhibited a distinct predicted mechanism compared to shuffled controls, as shown via unsupervised clustering (Fig. 6d).

To understand the sequence determinants of AlphaGenome’s predictions in CD34+ CMP, we performed ISM on the reference and alternative sequence of the oncogenic chr1:47239296:C>ACG variant. No mutations within 40 bp of the variant had a predicted effect on *TAL1* expression in the reference sequence (Fig. 6e). In contrast, the alternative sequence introduces a MYB motif at the variant position predicted to increase *TAL1* expression, chromatin accessibility, and H3K27ac chromatin marks (Fig. 6e), as discovered previously^42^. The model additionally highlights a second ETS-like motif nearby, which only affects *TAL1* expression in the alternative but not in the reference sequence. The role of this motif is currently unknown.

We then quantitatively evaluated AlphaGenome’s utility for analysing trait-altering non-coding variants. While models of molecular effect do not directly predict phenotypic consequences (Supplementary Note:

Trait-altering variants, Supplementary Fig. 10), AlphaGenome’s predictions could nevertheless prove useful for interpreting the gene-regulatory implications of such variants. Setting a high threshold on AlphaGenome’s quantitative scores (e.g., predicted expression change) strongly enriched for causal variant candidates among matched controls (Fig. 6f), albeit at the cost of low recall, particularly for GWAS variants. Inspecting some of these predictions for Mendelian disease variants in detail (Extended Data Fig. 9a-h) illustrates how the multimodal outputs of the model can be used to simultaneously elucidate a variant’s effect on transcription factor binding, accessibility and gene expression, particularly also predicting the direction of variant-induced expression changes. This makes AlphaGenome a useful complement to conservation-based measures of deleteriousness which are agnostic to a variant’s mechanism of action.

### Model and data ablations

To understand the contributions of key design and training decisions to AlphaGenome’s performance, we conducted ablation studies evaluating the impact of target resolution, sequence length, ensembling, distillation, and multimodal learning (Fig. 7). First, we found that base-pair resolution is important for achieving the highest performance. Training with target tracks at 1 bp resolution consistently yielded the best results, particularly for tasks requiring fine-scale detail like splicing (PSI5 and PSI3) and accessibility (ATAC), where performance generally declined as target resolution became coarser (Fig. 7a). In contrast, performance on tracks with coarser assay-specific resolution such as contact map correlation or histone ChIP-seq correlation was relatively insensitive to the target resolution used during training. Similarly, variant effect prediction metrics which involve aggregating effects over larger regions (like gene bodies or exons) were also robust to resolution changes.

We next evaluated AlphaGenome’s performance across various combinations of training sequence lengths and inference context lengths, finding that training with 1 Mb input sequences and subsequently performing inference using the full 1 Mb context yielded the best overall results (Fig. 7b). First, models trained on longer DNA sequences (up to the full 1 Mb) significantly outperformed those trained on shorter sequences (32kb or less), even when the latter were evaluated using 1 Mb contexts (Methods; Fig. 7b, purple crosses), underscoring the benefits of providing more sequence context during training. Second, inference also benefited from longer context: the 1 Mb-trained AlphaGenome model performed optimally when evaluated on the full 1 Mb sequence, with its performance progressively declining as shorter segments were used for inference input (Fig. 7b, blue dots). Finally, the 1 Mb-trained model, even when assessed on shorter inference contexts, often achieved comparable performance to models that were specifically trained and evaluated at those same matched shorter lengths (Fig. 7b, green triangles). This implies that the same model trained on the full 1 Mb sequence can also be used with a more limited context at inference time, providing an option for further increasing prediction speed if needed.

We compared distillation (Fig. 1c) against standard ensembling as a strategy for achieving high performance efficiently (Fig. 7c). Distillation using many teacher models (e.g., 64; orange crosses) produced single models whose performance was competitive with, or sometimes surpassed, mean ensembles of several independently pre-trained models (e.g., 4-model ensemble; blue dots). Interestingly, distillation of even just a single teacher model was beneficial for some variant effect prediction tasks such as caQTLs, splicing outliers or eQTL sign prediction. Overall, distillation offers a way to achieve strong performance with significantly reduced inference cost compared to evaluating large ensembles, facilitated by scalable training strategies like assigning one unique teacher per training device (Methods).

Finally, we assessed the importance of multimodal learning (Fig. 7d, Supplementary Fig. 11). The fully multimodal model generally outperformed models trained on single modality groups, confirming the overall benefit of integrating diverse data types for learning shared representations (Fig. 7). This benefit was task-dependent; for example, predicting accessibility variants was effective using only Accessibility data, whereas predicting eQTLs benefited from the full multimodal input ( a more complete set of experiments shown in Supplementary Fig. 11). Complementary experiments showed that excluding single modality groups during training typically resulted in only modest performance decreases, suggesting model robustness likely aided by informative redundancy between modalities (Supplementary Fig. 11, left columns). Together, these results demonstrate the value of the unified multimodal approach for achieving broad, high performance, particularly for tasks integrating diverse regulatory signals.

## Discussion

AlphaGenome advances efforts to decipher the genome’s regulatory code, offering a unified deep learning model that simultaneously predicts diverse functional genomic signals from megabase-scale DNA sequences. It matches or surpasses specialized SOTA models in their respective domains, which highlights the model’s robust grasp of fundamental regulatory principles encoded in DNA and its utility for mechanistic interpretation of non-coding variation. A core strength is AlphaGenome’s efficient multimodal variant effect prediction, which simultaneously scores variant impacts across all predicted modalities in a single inference pass. This integrated capability is crucial for understanding variants with complex mechanisms, as illustrated by the recapitulation of oncogenic *TAL1* variant effects, and could power large-scale analyses that dissect regulatory sequence elements genome-wide. Furthermore, AlphaGenome’s novel capability to directly model splice junctions enables a more holistic view of splice-disrupting variants.

Going forward, AlphaGenome holds promise across diverse biological disciplines. For molecular biology, AlphaGenome can serve as an engine for *in silico* experimentation, enabling rapid hypothesis generation and prioritization of resource-intensive wet-lab experiments. For rare disease diagnostics, AlphaGenome’s improved variant effect predictions could provide additional functional evidence to current annotation pipelines, offering new diagnostic avenues for non-coding variants of uncertain significance. AlphaGenome’s improved splicing, expression, and accessibility predictions could be used “in-the-loop” to accelerate sequence design applications such as therapeutic antisense oligonucleotides^48^ and tissue-specific enhancers^49,50^. Finally, AlphaGenome can complement the capabilities of generative models trained on DNA sequences by predicting functional properties of newly generated sequences^51^. To widely enable these broad applications, we provide accessible tooling to access AlphaGenome through a hosted model and API.

Despite its advances, AlphaGenome shares challenges common to current sequence-based models and has specific scope limitations. Accurately capturing the influence of distal regulatory elements (>100 kb) remains an ongoing objective. Moreover, although the model predicts tissue and cell type-specific genome tracks with some success, accurately recapitulating tissue-specific patterns across cellular contexts and predicting condition-specific variant effects remains challenging. Our species coverage remains limited to human and mouse, and the evaluations in this work are primarily human-focused. We have not yet benchmarked the model on personal genome prediction, which is a known weakness of models in this space^52,53^. Finally, application to complex trait analysis is limited given that AlphaGenome predicts molecular consequences of variants, while these phenotypes involve broader biological processes (including gene function, development, and environmental factors) beyond the model’s direct sequence-to-function scope.

Addressing these limitations motivates several future research directions. Key avenues include refining variant prediction accuracy and utility (e.g., through task-specific calibration, fine-tuning on perturbational datasets, or integration of single-cell data^15,54^), incorporating a broader range of data modalities (such as DNA methylation and DNA/RNA structural features), and pursuing foundational model improvements (e.g., leveraging DNA language models^55–57^, expanding multi-species capabilities, and developing robust methods for assay bias correction^10,58^). Moreover, integrating AlphaGenome predictions with other measures of variant effect, such as conservation-based scores, as well as existing data on gene functions and pathways could prove useful in advancing common and rare-variant analysis.

In summary, AlphaGenome provides a powerful and unified model for analyzing the regulatory genome. It significantly advances our ability to predict molecular functions and variant effects from DNA, offering valuable tools for biological discovery and applications in biotechnology. Ultimately, AlphaGenome serves as a foundational step towards the broader scientific goal of deciphering the complex cellular processes encoded in DNA sequence.

## Methods

### Data acquisition and processing

#### Reference Genomes and Genomic Annotations

AlphaGenome was trained using functional genomics data derived from human and mouse samples. Input sequences were extracted from the hg38 (human) and mm10 (mouse) reference genomes. For sequence intervals that extended beyond chromosomal boundaries, padding with ‘N’ characters was used to ensure consistent input length. Gene and transcript annotations used for model evaluations were based on the GTF V46 annotations (https://www.gencodegenes.org/human/release_46.html), unless otherwise specified.

#### Genomic coordinate system

For all genome intervals reported in the manuscript, we use a 0-based coordinate system (in which the starting element receives an index of 0, the second receives an index of 1, etc.). Each 0-based interval is non-inclusive or “half-open”, meaning that each reported interval includes the basepairs at the *start* position (again, reported in a 0-based manner) up to the base pair at the *end* -1 position, and excludes the base pair at the end position. The width of a 0-based interval is therefore simply *end - start*.

For variants, we follow the 1-based coordinate system that is conventional in the human genetics literature.

#### Metadata Standardization and Data Grouping

To ensure consistent data interpretation and enable robust aggregation across experiments, metadata were standardized using established ontologies. These included UBERON^59^ for tissue-level contexts, the Experimental Factor Ontology (EFO^60^) and Cell Line Ontology (CLO^61^) for cell lines, and Cell Ontology (CL^62^) for bulk primary cell type contexts.

Datasets were subsequently grouped to delineate distinct biological contexts for downstream analysis and signal averaging. The primary parameters for this grouping included:

- The assigned ontology term for the biosample, referenced using Compact Uniform Resource Identifiers (CURIEs). CURIEs are standardized, abbreviated codes (e.g., ‘UBERON:0001114’ for liver) that uniquely identify specific ontology terms.
- The specific assay type (e.g., ‘polyA plus RNA-seq’ versus ‘total RNA-seq’).
- The data source (e.g., ENCODE versus GTEx via RECOUNT3).
- Strand information, where applicable.
- For ChIP-seq experiments, the specific transcription factor (TF) or histone modification target (e.g., ‘CTCF’ or ‘H3K27ac’) was also a key grouping factor.

Within these groups, genomic signals were averaged across experiments to create representative tracks as detailed in the processing sections for individual assay types below. The selection of which experiments contributed to these averaged tracks was itself guided by hierarchical priority filters (as detailed in relevant sections, such as ENCODE Audit Tags Processing).

#### Training Data

To generate training targets for AlphaGenome, RNA-seq and epigenomic datasets were sourced from the ENCODE^63^, FANTOM5^64^, and GTEx^23^ consortia. Specifically, this included RNA-seq, PRO-cap, DNase-seq, ATAC-seq, transcription factor (TF) and histone ChIP-seq data from ENCODE; RNA-seq data from GTEx; CAGE data from Fantom5 portals; and genomic contact maps from the 4D Nucleome portal^65^.

**ENCODE Audit Tags Processing** A consistent quality control (QC) step based on ENCODE’s extensive audit system was applied to all ENCODE-derived datasets used in this study. For each experiment, metadata including detailed audit logs were downloaded via the ENCODE REST API (download link: https://www.encodeproject.org/search/?type=Experiment&format=json). Each ENCODE experiment can be associated with multiple audit flags, which are categorized by type and severity level (e.g., ERROR, NOT_COMPLIANT, WARNING; comprehensive details available at https://www.encodeproject.org/data-standards/audits/).

To facilitate an initial quality assessment, we processed these audit results to generate a single binary audit_filter_pass status for each experiment. An experiment was assigned audit_filter_pass = True (i.e., passed this initial QC) unless it contained any ‘ERROR’ level audits. Furthermore, experiments were also failed (assigned audit_filter_pass = False) if they possessed specific ‘NOT_COMPLIANT’ flags deemed critical for data quality. These exclusionary ‘NOT_COMPLIANT’ flags relating to issues with ligation motifs, long-range interactions, insufficient coverage or read counts, poor peak reproducibility or replicate concordance, low TSS enrichment, high PET redundancy, low library complexity, significant bottlenecking, or age consistency concerns (see Supplementary Table 6 for the comprehensive list and definitions of all exclusionary audit flags). This audit_filter_pass status served as a primary QC filter for all processed ENCODE datasets. The original, more granular audit categories for experiments passing this initial filter were also retained for consideration in subsequent prioritization steps.

**RNA-seq Data** To capture gene expression information, we integrated RNA-seq data from both ENCODE^63^ and GTEx^23^.

**ENCODE RNA-seq Data** Processing of ENCODE RNA-seq data commenced with the metadata file report (downloaded between January 9-17, 2025) and associated experiment, analysis, and biosample reports from the ENCODE portal. We filtered for ‘total RNA-seq’ and ‘polyA plus RNA-seq’ assays available as bigWig files for the GRCh38 (hg38) or mm10 assemblies. The selected output types were ‘minus strand signal of unique reads’, ‘plus strand signal of unique reads’, and ‘signal of unique reads’. Only ‘released’ files processed in 2020 or later were considered, and files known to have missing chromosome information were excluded.

To focus on baseline biological conditions, biosamples were selected if annotated as non-genetically modified, untreated, and not arrested in a specific cell cycle phase, according to the ‘Simple biosample summary’ field. Quality control measures included:

1. Applying an audit_filter_pass filter as described in the ENCODE Audit Tags Processing section above.
2. Ensuring a minimum mapped read length of 50 nucleotides (nt).
3. Selecting the single most representative bigWig file per experiment and output type, based on ENCODE’s replicate information and processing date.

Finally, to ensure consistency when multiple experiments represented the same biological context, we applied a hierarchical priority filtering step. This favored experiments that, in order:

1. Used non-genetically modified samples (this preference primarily impacts TF ChIP-seq selection).
2. Had less severe audit flags (PASS > WARNING > NOT_COMPLIANT).
3. Used paired-end sequencing.
4. Derived from primary cells (vs. *in vitro* differentiated cells) or tissues (vs. organoids).
5. Originated from more commonly profiled life stages (e.g., adult > embryonic > child).
6. Were not from subcellular fractions.

By default, ENCODE RNA-seq BigWigs represent read coverage per position and come normalized as Reads per Million (RPM). As each read is counted for each alignment position, this means that each bigWig (or bigWig pair for stranded assays) sums to 1 million reads times the read length. To ensure robust aggregation of tracks within a metadata group as defined above, we renormalized bigWigs to a common factor of 1 million reads times a common read length (100 in this work). This re-normalized signal representation was then used for all subsequent analyses.

**GTEx RNA-seq Data** We utilized processed bigWig files derived from the RECOUNT3 project^66^, which are unnormalized. From this, we selected only high-quality samples that corresponded to the set used for eQTL analysis within the GTEx consortium. In contrast to Borzoi, in which specific representative GTEx individuals were selected^3^, our analysis incorporated data from all individuals for each tissue. Each individual bigWig file was normalized as RPM similarly to ENCODE.

**RNA-seq Processing and Aggregation** Normalized RNA-seq tracks from ENCODE and GTEx were grouped by their ontology CURIE and assay type, distinguishing between ENCODE total RNA-seq, ENCODE polyA plus RNA-seq, and GTEx RNA-seq. Within each group, the normalized signals were averaged across all included experiments or individuals. This procedure yielded a final set of consolidated RNA-seq tracks, each representing the average normalized expression profile for a distinct biological context.

**CAGE Data** Cap Analysis Gene Expression (CAGE) data from the FANTOM5 consortium was used to profile transcription start site (TSS) activity^64,67^. We obtained CAGE Tag Starting Site (CTSS) information as BED files, filtering to include only samples derived from untreated biological conditions. The stranded counts from these CTSS BED files were converted into corresponding stranded signal bigWig files. Following the pre-aggregation normalization strategy applied to RNA-seq data, each individual CAGE signal bigWig track was normalized to represent a total of 100 million reads. These normalized tracks were then grouped based on their mapped ontology CURIE and assay type (‘LQhCAGE’ or ‘hCAGE’). Finally, within each group, the signals from the normalized tracks were averaged to produce a consolidated set of CAGE tracks representing average TSS activity across replicates or related samples for specific biological contexts.

**PRO-cap Data** For Precision Run-On sequencing with cap analysis (PRO-cap), we utilized a set of 12 stranded, processed bigWig files previously curated for the ProCapNet training dataset^8^, obtained directly from the ENCODE portal (https://www.encodeproject.org). Consistent with the normalization strategy used for RNA-seq and CAGE data, each of these 12 individual bigWig tracks was normalized so that its signal summed to 100 million. Further aggregation was not needed as the tracks were already unique by biosample and strand.

**DNase and ATAC Data** DNase-seq and ATAC-seq datasets were obtained from the ENCODE portal. Strict Fraction of Reads in Peaks (FRiP) score thresholds were applied (>10% for both assays, determined via manual inspection to balance coverage versus quality), along with different minimum read length requirements (*≥*36 nt for DNase-seq, *≥*45 nt for ATAC-seq, following ENCODE guidelines).

Instead of using ENCODE pre-processed p-value or fold-change signal bigWigs, which can obscure base-resolution information, we ingested the raw alignment (BAM) files for individual replicates. These BAM files were converted to base-resolution count bigWig files using the reads_to_bigwig.py script (from the ChromBPNet repository^10^, https://github.com/kundajelab/chrombpnet/blob/master/chrombpnet/helpers/preprocessing/reads_to_bigwig.py), with default parameters. These files record counts of Tn5 transposase insertions (for ATAC-seq) or DNase-I cleavage sites (for DNase-seq) at each base. This approach preserves the base-resolution nature of the data and facilitates decoupling enzyme cut bias from true signal by applying appropriate read shifts. Specifically, shifts of +4/-4 for ATAC-seq and 0/+1 for DNase-seq were applied to correct for the misalignment of strand-specific Tn5 and DNase-I position weight matrices (PWMs), respectively.

The assays were grouped by biosample ontology CURIE. The resulting base-resolution count BigWigs within each ontology group were then averaged. Following this averaging step, these consolidated tracks underwent post-aggregation normalization, rescaling the signal such that the total counts summed to 100 million insertions per track.

**Transcription Factor and Histone ChIP-seq Data** Processing of ENCODE transcription factor (TF) and histone ChIP-seq data began with the same comprehensive metadata acquisition from the ENCODE portal (https://www.encodeproject.org/, downloaded January 9-17, 2025) and initial bigWig file filtering criteria (GRCh38/mm10 assemblies, ‘released’ status, processed 2020 or later, exclusion of files with missing chromosome data) as detailed for ENCODE RNA-seq data. Specific to ChIP-seq, only files representing ‘fold change over control’ signal were selected.

Key biosample filters, focusing on baseline cellular states (untreated, non-arrested), were applied as described previously, with specific criteria for genetic modifications: Histone ChIP-seq samples were required to be non-genetically modified, while for TF ChIP-seq, genetically modified samples were initially included if necessary (to enable profiling of TFs not typically expressed in the most studied cancer cell lines used by ENCODE), but non-modified samples were strongly preferred during priority filtering.

For quality control:

1. Experiments were filtered based on the audit_filter_pass status, determined as described in the ENCODE Audit Tags Processing section.
2. A frip_filter_pass flag was assigned based on the Fraction of Reads in Peaks (FRiP) score (>1% for TF ChIP-seq, >6% for Histone ChIP-seq). Experiments were retained if they passed either the initial audit or the FRiP filter (audit_filter_pass = True or frip_filter_pass = True), allowing inclusion of high-signal experiments potentially affected by single-replicate audit issues.

A minimum mapped read length of 50 nt was required, consistent with RNA-seq data. Where multiple signal files existed per experiment accession, the most recent file representing the most complete set of biological replicates (guided by the ‘Biological replicates’ metadata field) was chosen.

Finally, to ensure consistency for experiments representing the same biological context (cell type and target), we applied the hierarchical priority filtering step as outlined in the *ENCODE RNA-seq Data* section. The preference for non-genetically modified samples within this scheme primarily impacts TF ChIP-seq selection.

Following these steps, selected ChIP-seq experiments were grouped by their ontology CURIE, strand, and specific ChIP-seq target (TF or histone modification, Supplementary Table 2). Within each group, the ‘fold change over control’ signal was averaged. These resulting ChIP-seq tracks were used without further normalization, as fold-change values inherently account for control signals.

**Summary of data processing parameters** Supplementary Table 7 provides a consolidated summary of the data processing parameters applied to generate each type of 1D genomic track used in this study. It outlines critical aspects for each assay, including data sources, quality control filters, normalization approaches, and final aggregation methods.

**Preparation of RNA-seq Data for Splicing Analyses** Training data for AlphaGenome’s splicing-related predictions (splice junctions, splice site usage (SSU), and splice site classification) were derived from the same ENCODE and GTEx RNA-seq datasets used for gene expression analyses, and samples were grouped by ontology CURIEs consistently with the RNA-seq processing pipeline.

To quantify splice junction count, reads from each RNA-seq sample were realigned using STAR (version 2.7.11b)^68^ from the BAM files downloaded from GTEx and ENCODE. For human samples, alignment was performed against the GRCh38.p13 reference genome, with GENCODE v32 gene annotations guiding splice junction discovery. For mouse samples, the GRCm38.p6 reference genome and GENCODE vM23 annotations were used.

Key STAR alignment parameters were configured to optimize for junction detection, including setting a minimum splice junction overhang of 8 base pairs (--alignSJoverhangMin 8), allowing a maximum of 20 alignments for multi-mapping reads (--outFilterMultimapNmax 20), defining standard intron size limits (e.g., --alignIntronMin 20, --alignIntronMax 1000000), and outputting strand information derived splice dinucleotide motifs (--outSAMstrandField intronMotif).

The precise STAR command used was:

The primary output files containing splice junction information (sj.out.tab) from STAR served as the raw data for all subsequent splicing data curation, specifically to define training targets for the three types of splicing-related predictions. Samtools (version 1.21)^69^ was used for standard intermediate processing of BAM files generated during alignment.

**1. Splice Junction Quantification, Filtering, and Normalization** Individual splice junction output files for each sample were first combined into a single comprehensive table, indexed by chromosome, junction start coordinate, junction end coordinate, and strand, separately for human and mouse species.

A stringent quality filtering pipeline was then applied to these compiled junction lists using the splicemap package (available at https://github.com/gagneurlab/splicemap) to ensure high data fidelity:

- *Human GTEx Samples*: A junction was retained if its 90th percentile read count across all GTEx samples was greater than 1, AND the median of total read counts supporting any alternative splicing event sharing either its donor or acceptor site was at least 1.
- *Human ENCODE Samples*: Junctions from human ENCODE RNA-seq samples were filtered against the set of high-confidence junctions derived from GTEx; any ENCODE human RNA-seq splice junction not present in the filtered GTEx junction set was discarded.
- *Mouse ENCODE Samples*: For mouse ENCODE RNA-seq samples, a junction was retained if its median read count across all mouse RNA-seq samples within the same ontology CURIE group was greater than 3.

After these filtering steps, the retained junction counts for each sample were normalized to 1 million total filtered junction reads per sample.

For use in model training (loss calculation) and evaluation, these normalized junction counts underwent further tissue-specific preprocessing. Within each tissue:

1. Raw counts were first clipped at the 99.99th percentile for that tissue to mitigate the influence of extreme outliers.
2. Subsequently, these clipped counts were scaled by dividing by the mean count value, where this mean was calculated only across actively expressed junctions (defined as those with a clipped count > 0) within that specific tissue.

During training on splice junctions, the donor-acceptor pairing of a maximum of 512 splice sites on each strand are considered per input interval. If the sampled interval has more than 512 splice sites, we narrow the interval size to consider splice junctions until a maximum of 512 splice sites are in the interval on either strands. (Note: This only affects the interval used for inference; the input sequence length of the model, for example 1 Mb, remains unchanged). For each strand and each CURIE condition, splice junction counts are represented as a square matrix of shape [512, 512] corresponding to donor/acceptor pairs. Each element of the matrix represents the observed normalized read counts for the donor/acceptor pair. If less than 512 splice sites are observed, the matrix is padded to 512 with 0.

1. **2. Splice Site Usage** Splice Site Usage (SSU) was calculated for each potential splice site using the formula:

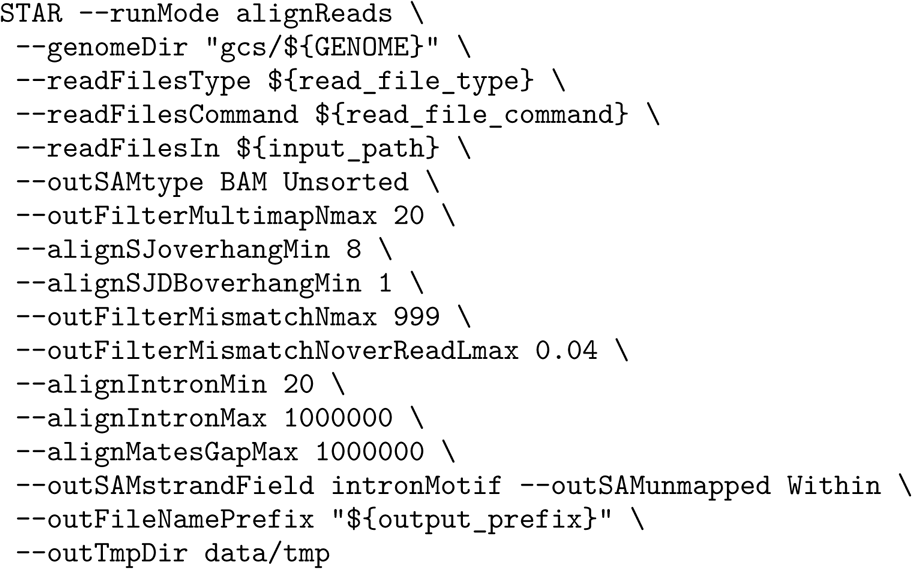

We adapted the basic splice site strength definition from Dent et al^70^. SSU quantification was performed using a custom script. For this calculation, we considered all reads spanning the splice sites regardless of the strand. SSU counting was done with a custom script. Reads flagged as PCR/optical duplicates, those with a mapping quality (MQ) below 30, or reads containing base calls with a base quality (BQ) below 20 were excluded from the counts. The counting of each RNA-seq sample was performed independently, and the SSU for each RNA-seq sample was calculated independently. Only splice sites that were detected in the corresponding STAR splice junction output (sj.out.tab files) and passed the above splice junction filtering step were considered in this quantification process.

The splice site usage has two tracks per CURIE condition, corresponding to the two strands. Unlike splice sites, SSU does not distinguish between donor usage and acceptor usage.

1. **3. Splice Site Definition for Classification** The set of splice sites used for training the splice site classification task was defined as the union of all unique donor and acceptor sites present in the filtered splice junction data (from step 2 above) for each ontology CURIE. Notably, unlike splice junction counts or SSU values which can be tissue/sample-specific, the defined splice site training examples were not treated as tissue-specific.

The splice site classification task was formulated as a 5-class classification problem, where each relevant position could be classified as:

- Donor site on the positive strand (Donor+)
- Acceptor site on the positive strand (Acceptor+)
- Donor site on the negative strand (Donor-)
- Acceptor site on the negative strand (Acceptor-)
- Not a splice site on either strand

**Contact maps** Chromatin contact maps, which represent average inter-nucleotide contact probabilities typically derived from assays like Hi-C or Micro-C, are crucial for understanding gene regulation via 3D genome organization. We sourced contact map datasets from the 4D Nucleome portal ( https://data.4dnucleome.org, accessed 2021/03/04), aiming for quality comparable to or exceeding that of the 5 datasets used by Akita^71^ and the 2 datasets used by Orca. To achieve this, we selected datasets provided as multi-resolution cooler files (.mcool) that were at least 7.74 GB (the size of the smallest Akita contact map). We retained only those files that were at 1000 bp resolution. This resulted in a curated collection of twenty-eight human and eight mouse datasets (see Supplementary Table 2 for file accessions), which importantly includes the 5 Akita and 2 Orca datasets. We did not directly compare our model’s performance to Akita’s, as their dataset preprocessing includes an additional Gaussian blurring step not present in Orca’s methods or ours, which prevents direct output comparisons.

We preprocessed these contact maps following the protocol established for the Orca 1 Mb model^19^. Briefly, 1000 bp resolution contact maps underwent two standard preprocessing steps: matrix balancing to scale the values and adaptive coarse-graining to apply smoothing. Subsequently, a distance-based normalization was applied:

- First, an average coarse-grained contact value was computed for each pairwise genomic distance (𝑚𝑒𝑎𝑛[𝑎𝑏𝑠(𝑖 *−* 𝑗)]) across each dataset (excluding all zero-valued bins).
- Distance-based normalization was then applied: the normalized contact map values y(i, j) were computed as the log-fold change over the distance-dependent means: 𝑦[𝑖, 𝑗] = log((𝑥[𝑖, 𝑗] + 𝑒𝑝𝑠)/(𝑚𝑒𝑎𝑛[𝑎𝑏𝑠(𝑖 *−* 𝑗)] + 𝑒𝑝𝑠)). Here 𝑥[𝑖, 𝑗] is the coarse-grained count and the numerical relaxation constant, 𝑒𝑝𝑠, was set to the minimum value of the mean profile.

During model training, these preprocessed 1000 bp resolution contact matrices were interpolated to the 2048 bp resolution of AlphaGenome’s pairwise representation blocks and aligned with the sampled input intervals. This interpolation used an area-weighted averaging scheme, where the value for each target 2048 bp bin was the average of overlapping source 1000 bp bins, weighted by their fractional area of overlap.

**Dataset splitting and cross-validation** For a robust comparison with Borzoi and other baselines, we utilized the identical cross-validation fold definitions (*fold-0*, *fold-1*, *fold-2*, and *fold-3*) previously established by Borzoi for both human and mouse genomes (sequences_*.bed.gz files in https://github.com/ calico/borzoi/tree/5c9358222b5026abb733ed5fb84f3f6c77239b37/data). Each genome is split into eight distinct sections; each fold uses six sections for training, one for validation, and one for testing. In addition to these cross-validation setups, we also trained comprehensive *all-folds* models using all eight genomic sections for the training set; these models were exclusively evaluated on variant interpretation benchmarks.

Within the genomic regions assigned to each set by these folds, we specifically used the pre-defined target intervals (approximately 196kb) defined in the Borzoi study. Since AlphaGenome operates on 1 Mb input and output sequences, each 1 Mb input window was centered around the midpoint of the corresponding 196kb Borzoi target interval. Crucially, to prevent data leakage between sets arising from AlphaGenome’s larger input window, a strict exclusion criterion was applied: any validation or test interval was removed from its respective set if its corresponding 1 Mb AlphaGenome input window overlapped with any 1 Mb input window derived from a training interval within that same fold.

**Final data representations** Processed genomic tracks, serving as model prediction targets, were converted to brain floating point (bfloat16) format for numerical efficiency and stored in z-standard compressed sharded matrices. Most data tracks were maintained at base-pair resolution, except for ChIP-seq (TF and Histone) tracks, which represent fold-change values, and were stored as base-resolution cumulative sums. This strategy allows for efficient querying of their average signal at 128 bp resolution.

The model directly uses the track values resulting from the upstream processing steps, which typically represent normalized read or insertion counts (often scaled to a total of 100 million signals per track for sequencing assays) or fold-change enrichments (for ChIP-seq). No additional scaling transformations (e.g., log-scaling or z-score normalization across tracks) are applied before model input. Predicted splice junction counts are an exception; while their values are additionally scaled (as described in the training methods), this does not affect their primary use in calculating ratios for relative splice site assessment.

### Model

AlphaGenome is a deep learning based model that processes a 1 Mb DNA sequence along with a species identifier (human or mouse) to predict a diverse array of genomic features. These outputs include 1D tracks representing signals such as chromatin accessibility, TF binding, and gene expression at various resolutions; 2D chromosomal contact maps; and specific splicing-related predictions like splice site probabilities, splice site usage (SSU), and splice junction counts.

AlphaGenome employs an encoder-decoder framework that integrates convolutional and transformer components, drawing inspiration from models such as Enformer^2^ and Borzoi^3^. It introduces several key architectural features, including a U-Net-style decoder incorporating residual (skip) connections from the encoder. The model processes information through five core components: (1) a sequence encoder, (2) a transformer tower, (3) pairwise interaction blocks, (4) a sequence decoder, and (5) task-specific output heads. AlphaGenome has approximately 450 million trainable parameters (20% in the encoder, 28% in the sequence transformer, 15% in the pairwise blocks, 25% in the decoder, and 12% in the output embedding and prediction heads). The model was implemented using JAX^72^ and Haiku^73^.

#### Sequence Encoder

The sequence encoder progressively downsamples the input DNA sequence from 1 bp resolution (length 1 Mb or 2^20^ bp) to 128 bp resolution embeddings (length 8192 or 2^13^) over 7 stages. The number of feature channels increases from 768 up to 1536 to enhance representational capacity. This downsampling is achieved through repeated convolutional blocks, each followed by max-pooling with a stride of 2. The corresponding pseudocode is as follows:

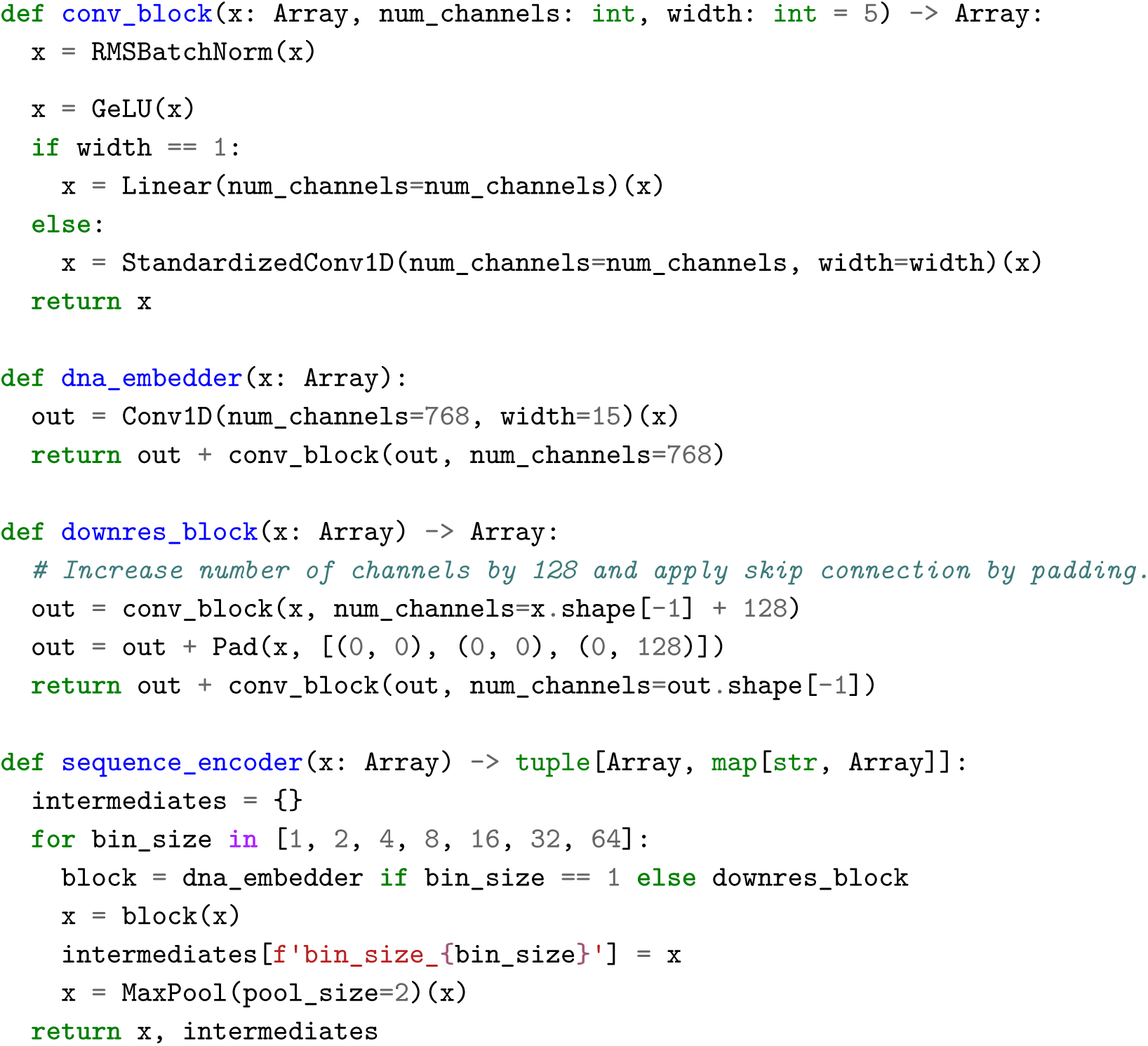

The core component of the above code is a conv_block, which applies root mean square batch normalization (RMSBatchNorm—same as BatchNorm^74^ with learned scale and offset, but without shifting by the sample mean), a GeLU^75^ activation, and then a standardized 1D convolution (StandardizedConv1D) with a kernel width of 5. Following Brock *et al.*^76^, these convolutions employ scaled weight standardization – re-parameterizing by standardizing and scaling the weights – which we found effective for stabilizing activation magnitudes. Each RMSBatchNorm layer maintains an exponential moving average (EMA, decay 0.9) of per-channel variance during training; this EMA variance is used for normalization during inference. To facilitate training and gradient flow^77^, residual skip connections are added around each convolutional block. When channel dimensions differ across a residual connection (as in downres_block), the input is padded with zeros before addition. All convolution layers used “same” padding.

The initial stage uses the dna_embedder block, which applies a wider 1D convolution (kernel width 15, embedding to 768 channels), similar to Enformer^2^, followed by a residual conv_block. The subsequent six downres_block stages each apply two conv_block layers (the first increasing the channel count by 128) with skip connections. Intermediate representations from each of the 7 stages, corresponding to resolutions of 1 bp, 2 bp, 4 bp, . . . , 64 bp before the max-pooling step, are stored for use as U-net skip connections in the sequence decoder. The final output of the encoder has a resolution of 128 bp, sequence axis length 8192 (1 Mb / 128 bp) and 1536 channels (768 + 6 * 128).

#### Transformer Tower

Following the encoder, a transformer tower processes the 128 bp resolution sequence embeddings to model long-range interactions across the full 1 Mb input context. The tower consists of 9 stacked transformer blocks. Each block contains a multi-head attention (MHA) layer followed by a two-layer perceptron (MLP) layer, both wrapped in residual connections. We use standard array dimension conventions where ‘B’ denotes batch size, ‘S’ sequence length (𝑆 = 1𝑀𝑏/128 𝑏𝑝 = 8192), ‘C’ channels, ‘P’ pairwise sequence length, and ‘F’ pairwise channels. In pseudocode, these components can be written as:

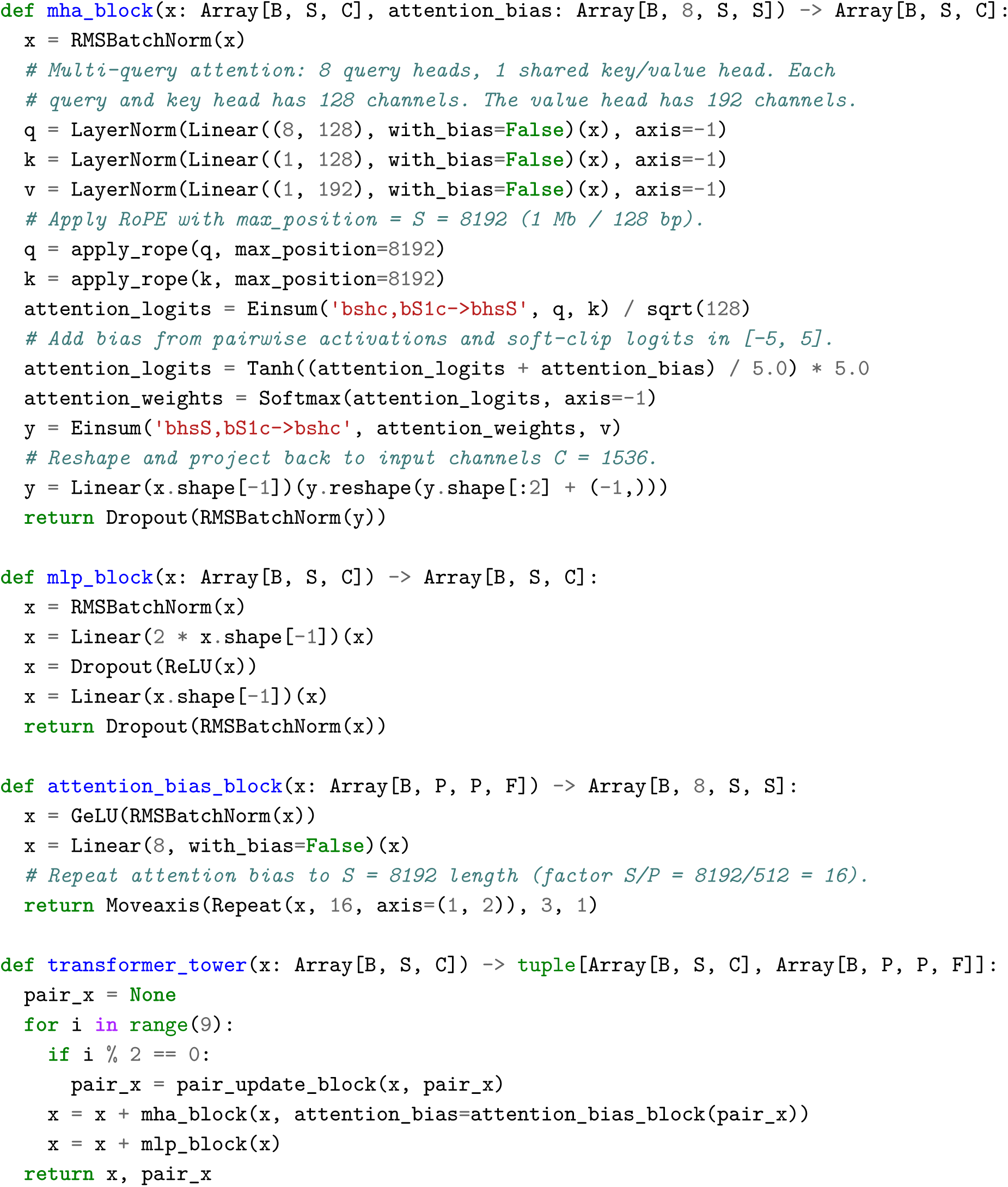

Before every second MHA block, pairwise representations at 2048 bp resolution (𝑃 = 1𝑀𝑏/2048 𝑏𝑝 = 512) are initialized or updated based on the sequence embeddings (detailed in Pairwise blocks section). They are primarily used for contact map prediction but also provide a bias term to the MHA layers. The attention_bias_block function projects the pairwise channels (F) to match the number of attention heads (8) and then repeats the result 16 times (2048 bp / 128 bp) along the two sequence axes to match the sequence length dimension. This bias is added directly to the attention logits before the softmax, analogous to techniques used in AlphaFold2^78^.

Our MHA implementation (mha_block) uses multi-query attention^79^, where key (k) and value (v) projections are shared across all 8 attention heads, reducing computational cost. We apply smooth clipping to the attention logits^80^, constraining them to the range [-5.0, 5.0] using a scaled hyperbolic tangent function before the Softmax. Relative positional information is incorporated using Rotary Position Embeddings (RoPE^81^) applied to query and key vectors. We use a modified calculation for the inverse frequencies compared to the standard RoPE formulation (apply_rope). This modification reduces the density of rotational frequencies corresponding to short relative distances and is applied consistently throughout the model. The core of this implementation is detailed in pseudocode:

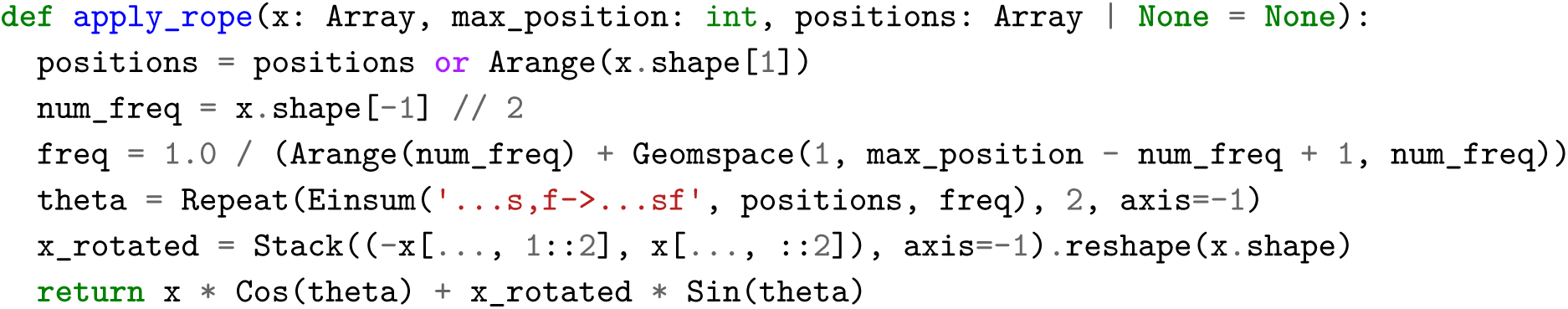

The MLP block (mlp_block) follows standard transformer design, using ReLU activation and expanding the number of channels by a factor of 2 in the hidden layer. Dropout is applied within the MHA and MLP blocks during training only, with rates of 0.3 for pretraining and 0.1 for distillation.

#### Pairwise Blocks

Interleaved with the transformer tower blocks (specifically, executed before the MHA layer in blocks 0, 2, 4, 6, and 8), pair_update_block modules operate on and update a 2D tensor representing pairwise interactions between sequence positions. These pairwise representations have 2048 bp resolution (sequence length 𝑃 = 512) and have 𝐾 = 128 channels; they are primarily used for contact map prediction but also contribute a bias to the transformer’s attention mechanism (described in the Transformer Tower section). Each pair_update_block first computes an update from the sequence embeddings (sequence_to_pair_block), which initializes or is added to the previous pairwise state, and then performs two residual refinement steps (row_attention_block and pair_mlp_block).

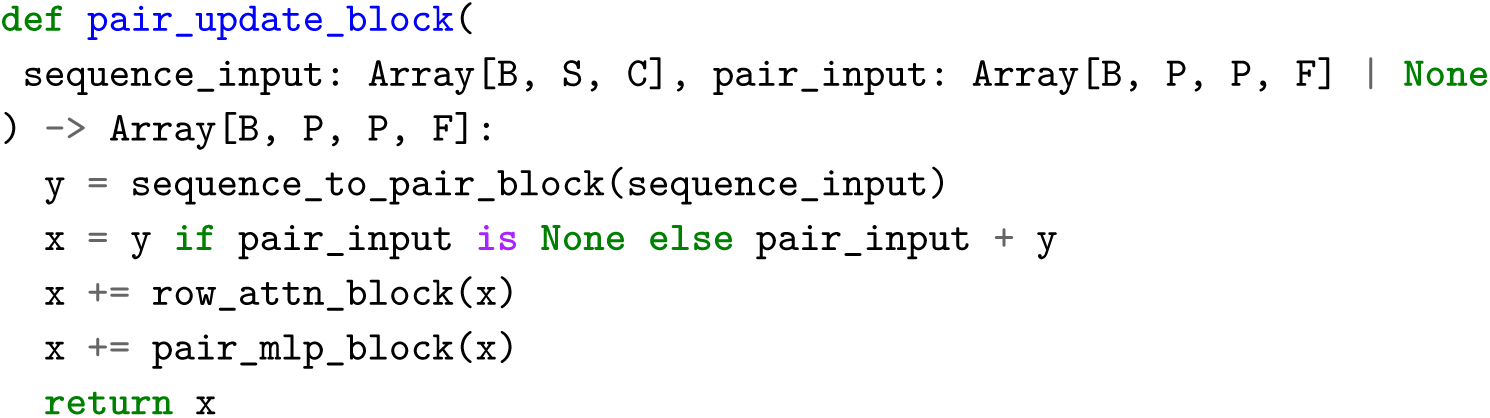

The sequence_to_pair_block function first downsamples the 128 bp resolution sequence embeddings (𝑆 = 8192) to 2048 bp resolution (𝑃 = 512) using average pooling. It then computes pairwise features using a form of attention enhanced with relative positional encodings. The positional encodings are generated by central_mask_features using 32 exponentially spaced distance thresholds. Directionality is then incorporated by concatenating these 32 features with a copy multiplied by the sign of the relative distance, yielding a 64-channel positional representation. This relative attention term is calculated efficiently using the relative_shift operation, as detailed in Transformer-XL^82^. The final sequence-to-pair representation is obtained by projecting the combined query-key and relative position terms, adding an outer sum of projections derived from the sequence embeddings, and applying dropout.

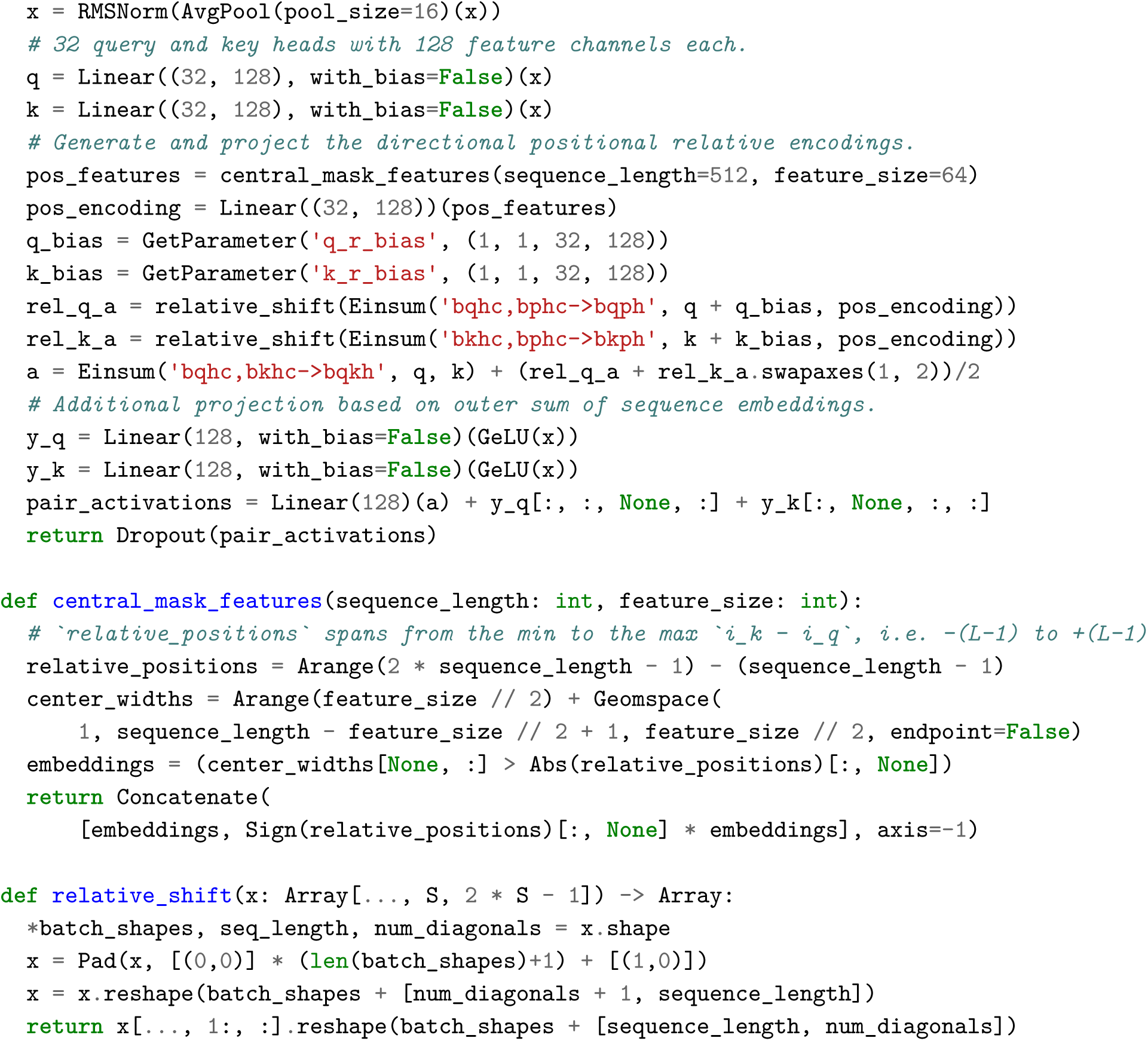

The pairwise embeddings undergo further refinement by two transformer-style operations, each within a residual connection. The row_attention_block implements a self-attention mechanism that operates only along the second axis of the PxP pairwise matrix (each position (i,j) attends to all positions (i,k) in the same row i). We found that applying a second attention block that operates over columns yielded no empirical accuracy gains for downstream tasks (likely due to the explicit symmetrization applied before contact map prediction) and was less computationally efficient, particularly in distributed training settings. The pair_mlp_block then applies a standard two-layer MLP with a ReLU activation and 2x channel expansion for further feature refinement.

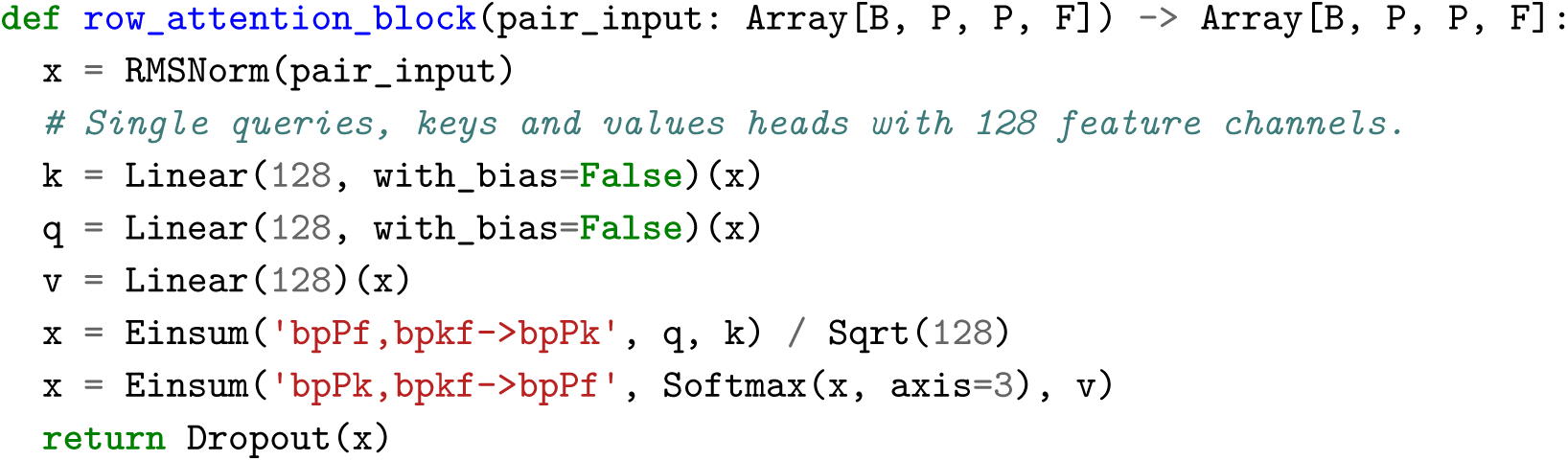

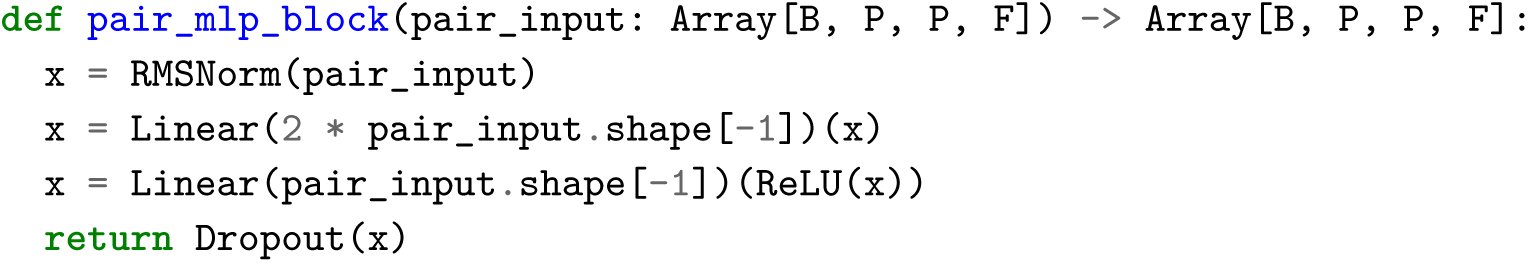

#### Sequence Decoder

The sequence decoder mirrors the encoder’s structure but operates in reverse: it progressively upsamples the 128 bp resolution embeddings (output from the transformer tower, with 1536 channels) back to the original 1 bp resolution, while simultaneously decreasing the number of feature channels down to 768. This is achieved through 7 iterative stages, each applying an upres_block module. As is standard in U-Net architectures^18^, the decoder incorporates skip connections, adding embeddings (intermediates) stored at corresponding resolutions during the encoder pass. The corresponding pseudocode is as follows:

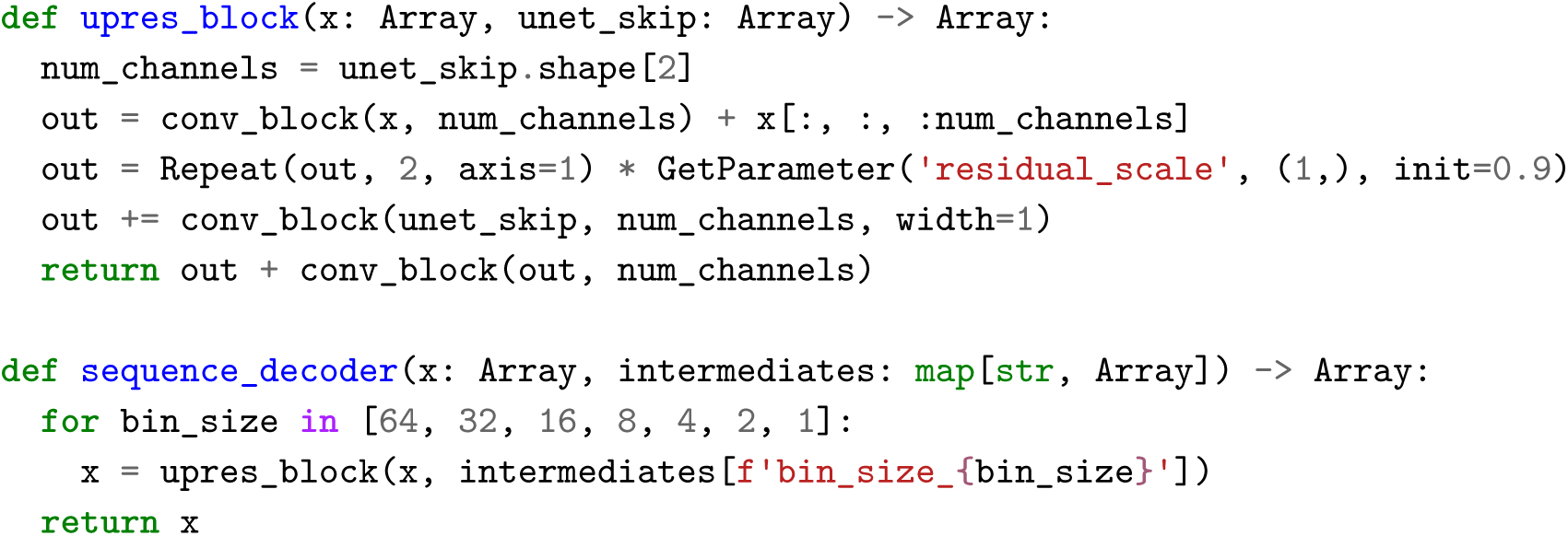

Each upres_block takes the output from the previous stage and the corresponding resolution skip connection from the encoder. First, a conv_block reduces the channel dimension of the embedding to match that of the skip connection. As usual, the convolution is paired with a residual connection which handles the reduction in channel dimension by cropping. The result is then upsampled by a factor of two along the sequence length axis via repetition and scaled by a learnable coefficient specific to that resolution level. Concurrently, the incoming skip connection is processed through its own conv_block (using a kernel width of 1). This processed skip connection is then added to the scaled, upsampled representation. Finally, this combined tensor passes through one more conv_block, again including an internal residual connection, before being passed to the next stage or returned as the final 1 bp resolution sequence embedding.

#### Output Heads

The AlphaGenome architecture, after processing the input DNA through its encoder, transformer, and decoder stages, produces key internal representations at multiple resolutions. These learned embeddings (specifically the 1D embeddings at 1 bp and 128 bp resolution along the sequence, and the 2D pairwise embedding at 2048 bp resolution) form the basis for predicting a diverse array of genomic features. Predictions are adapted to the specific organism (human or mouse) by incorporating learned, organism-specific embeddings within these functions. The following pseudocode illustrates the core data flow that generates these embeddings via the main model components (sequence_encoder, transformer_tower, sequence_decoder) and output embedding functions (output_embedder, output_pair).

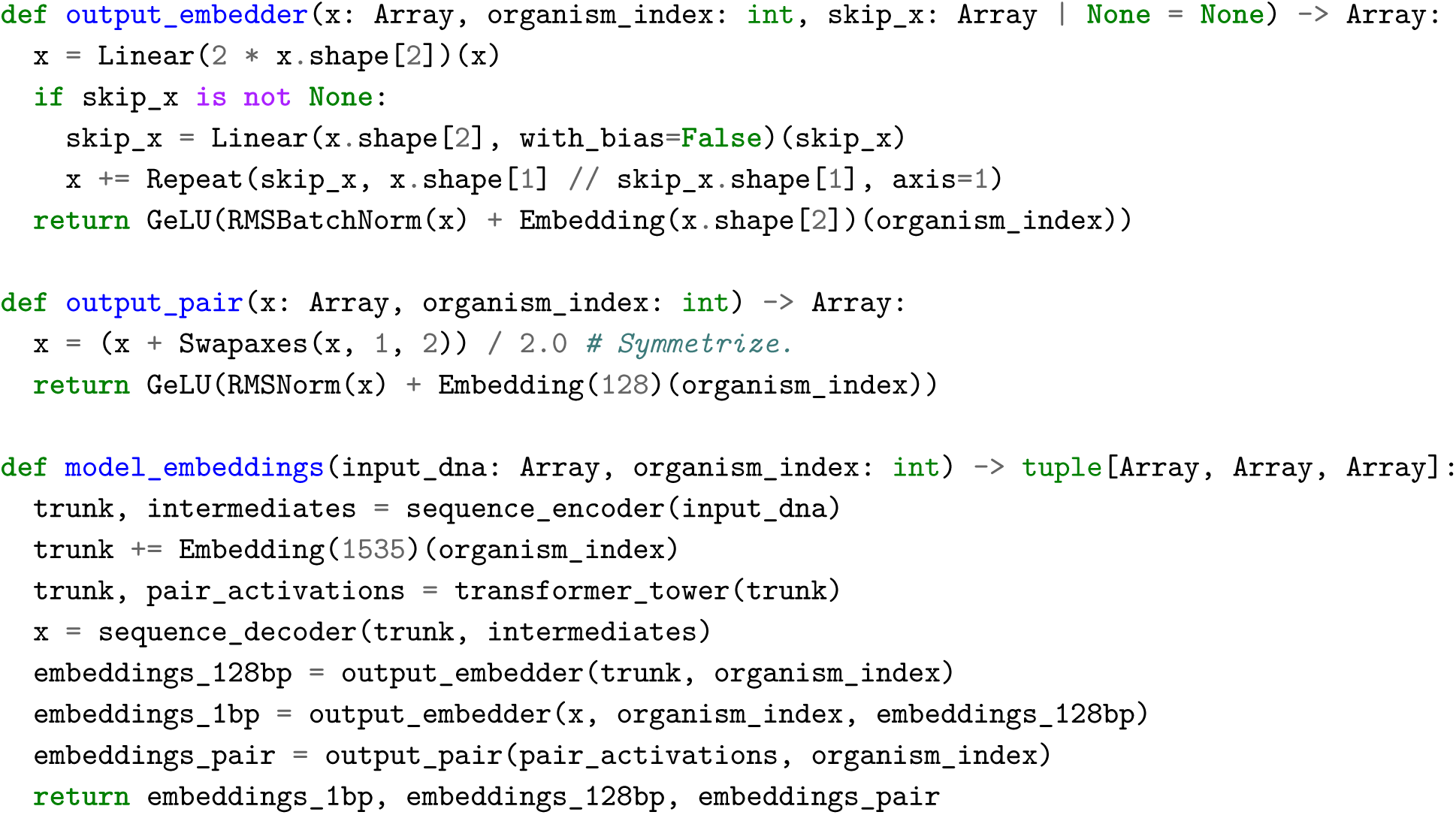

Beyond the organism-specific embeddings applied during the generation of these shared representations (embeddings_1bp, embeddings_128bp, embeddings_pair), each output head itself learns a distinct set of parameters for human and mouse predictions. The subsequent sections detail the specific parameterization and loss function defined for each prediction modality. The total training loss for AlphaGenome is the sum of these losses defined by each head with no additional weighting coefficients.

**RNA-seq, CAGE, ATAC, DNase and PRO-Cap Output Heads** Predictions for these assays are generated from both the 1 bp and 128 bp resolution embeddings. Each resolution uses a dedicated head consisting of a linear layer mapping the input channel dimension to the number of target tracks for that assay type. A softplus activation is applied to the linear layer’s output, which is then multiplied by a learnable, per-track positive scaling factor, ensuring non-negative outputs. Strand-specific readouts are treated as distinct output tracks. Each head predicts tracks as in the pseudocode:

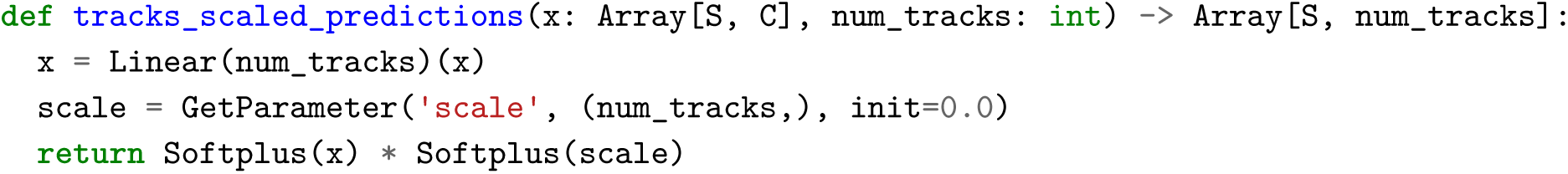

To improve numerical stability, experimental target tracks undergo a scaling procedure before loss calculation (targets_scaling). Tracks are first normalized by dividing by the mean of their non-zero values (pre-calculated per track across the entire genome during dataset generation). High values are then dampened using square-root-based smooth clipping. For RNA-Seq tracks only, an additional power transformation is applied for further compression. These scaling and clipping operations are reversed (predictions_scaling) when evaluating model predictions against the original experimental data. The scaling and inverse scaling functions are:

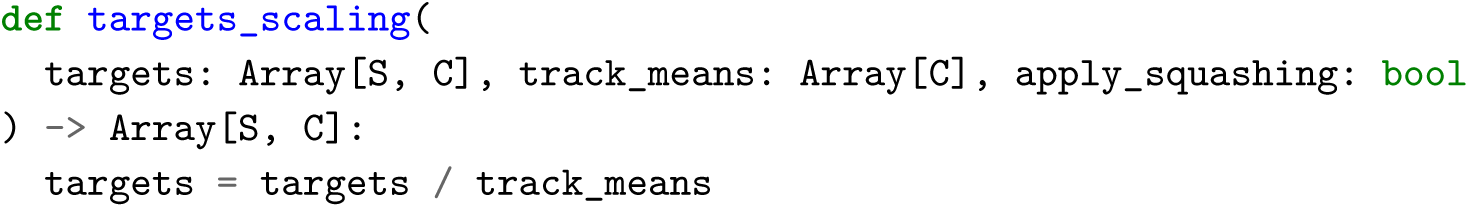

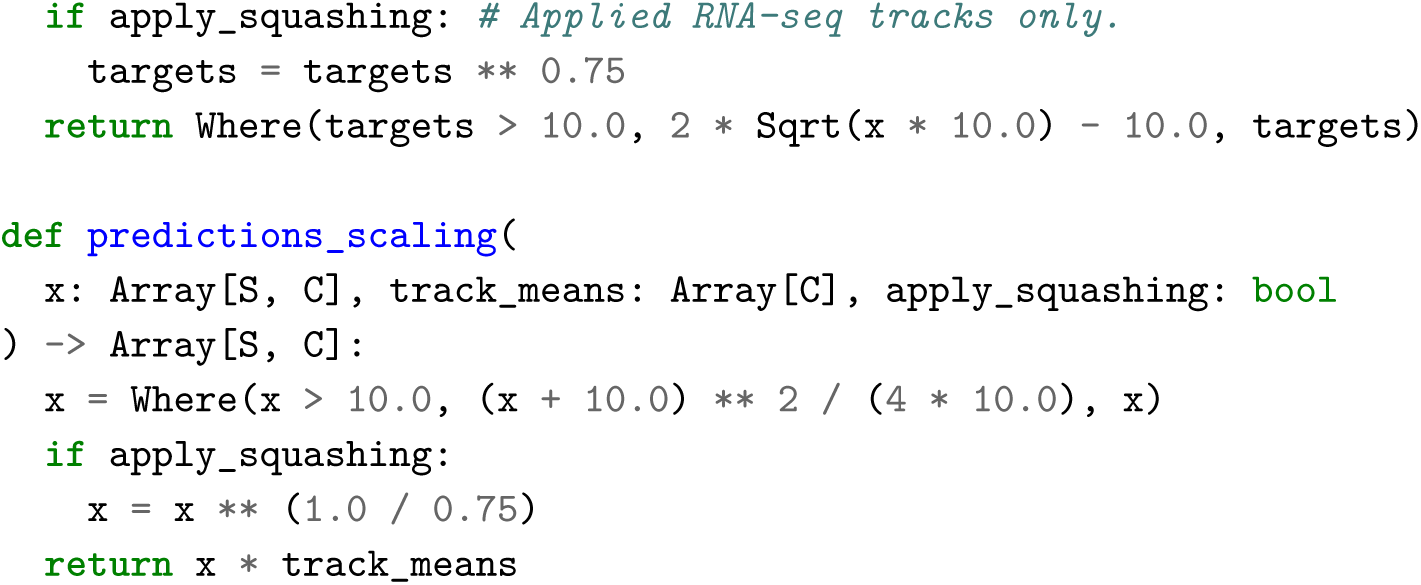

The training loss follows the approach of Borzoi, calculated as a weighted sum of Poisson and Multinomial negative log-likelihoods (NLL). For each track, the relevant sequence axis (e.g., 2^20^ bins for 1 bp, 8192 bins for 128 bp resolution outputs) is divided into 8 equal segments. Within each segment, the loss combines two components:

1. A Multinomial NLL term comparing the predicted distribution of counts across bins to the target distribution, which ensures the model learns the relative values and overall shape of the predictions within each segment. This multinomial component is weighted by a factor of 5.0, as in Borzoi.
2. A Poisson NLL term comparing the sum of predicted and target counts for each segment, which encourages the model to learn the accurate overall count (regardless of shape). This term is scaled

inversely by the segment length (multinomial_resolution = 2^17^).

Calculating the loss over 8 segments, rather than the full sequence, mitigates numerical instability (using smaller segments was found to empirically degrade performance). The loss function is implemented as shown in the pseudocode:

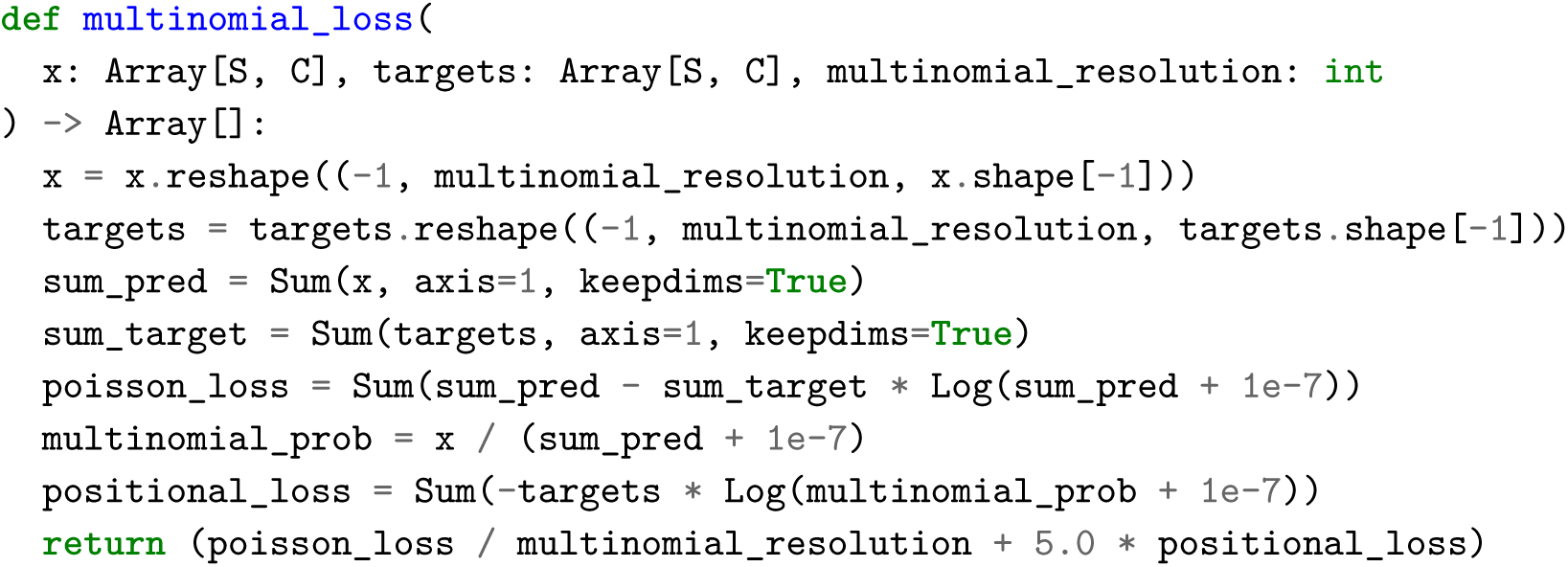

For RNA-seq tracks, an additional loss component, inspired by Decima^54^, promotes tissue-specific expression patterns. First, predicted and scaled target counts are aggregated within annotated gene boundaries found in the input interval and subsequently normalized by gene length to yield per-gene counts. These per-gene counts are then used to compute a loss across the tissue/cell type dimension for each gene. This loss calculation mirrors the structure of the main track loss described above: it combines a Poisson NLL term on the total normalized expression per gene across all tissues and a Multinomial NLL term on the distribution across tissues. The Multinomial term within this calculation is weighted by a factor of 5.0. This auxiliary gene-level loss contributes to the model’s total training loss with an overall weight of 0.1.

**TF ChIP-seq and Histone ChIP-seq Output Heads** These tracks are predicted from the 128 bp resolution embeddings. The prediction head structure, target scaling (without squashing), loss calculation (Poisson and Multinomial over 8 segments), and inverse scaling are identical to those used for the other genomic tracks described above, but applied only to the 128 bp resolution pathway (e.g., multinomial_resolution = 2^17^/128 = 1024).

**Contact Maps Output Head** Contact frequency predictions are derived from the 2048 bp resolution pairwise embeddings. The raw pairwise activations are first explicitly symmetrized. An organism-specific embedding is added, followed by RMSNorm and GeLU activation. A final linear transformation (not shown in snippets) maps these processed pairwise embeddings (with 𝐾 = 128) to the predicted target contact map tracks. The contact map outputs are trained using a mean squared error (MSE) loss between the predictions and the targets.

**Splice Sites Classification Output Head** This head predicts the probability of each base belonging to one of five classes: Donor+, Acceptor+, Donor-, Acceptor-, or not a splice site (see Splice Site Definition for Classification section in Training Data). A linear layer is applied to the 1 bp resolution embeddings that maps to 5 logits per base, followed by a Softmax activation to produce class probabilities. Training uses a standard per-base cross-entropy loss function against the true splice site labels.

**Splice Site Usage Output Head** This head predicts the proportion of splicing events utilizing each potential splice site (SSU), separately for each strand and tissue/cell type. It takes the 1 bp resolution embeddings as input. A linear layer maps the embedding dimension to the number of tissues/cell types per strand at each potential splice site location, followed by a Sigmoid activation to constrain outputs between 0 and 1. The model is trained using a binary cross-entropy loss against the observed SSU values.

**Splice Junctions Output Head** This head predicts counts for potential splice junctions between donor and acceptor sites. It operates on the 1 bp resolution embeddings. Because each sequence in the batch might have a different number of donor and acceptor splice sites, in practice we perform the calculation using batch padding. In the pseudocode below, we show the calculation for a single sequence:

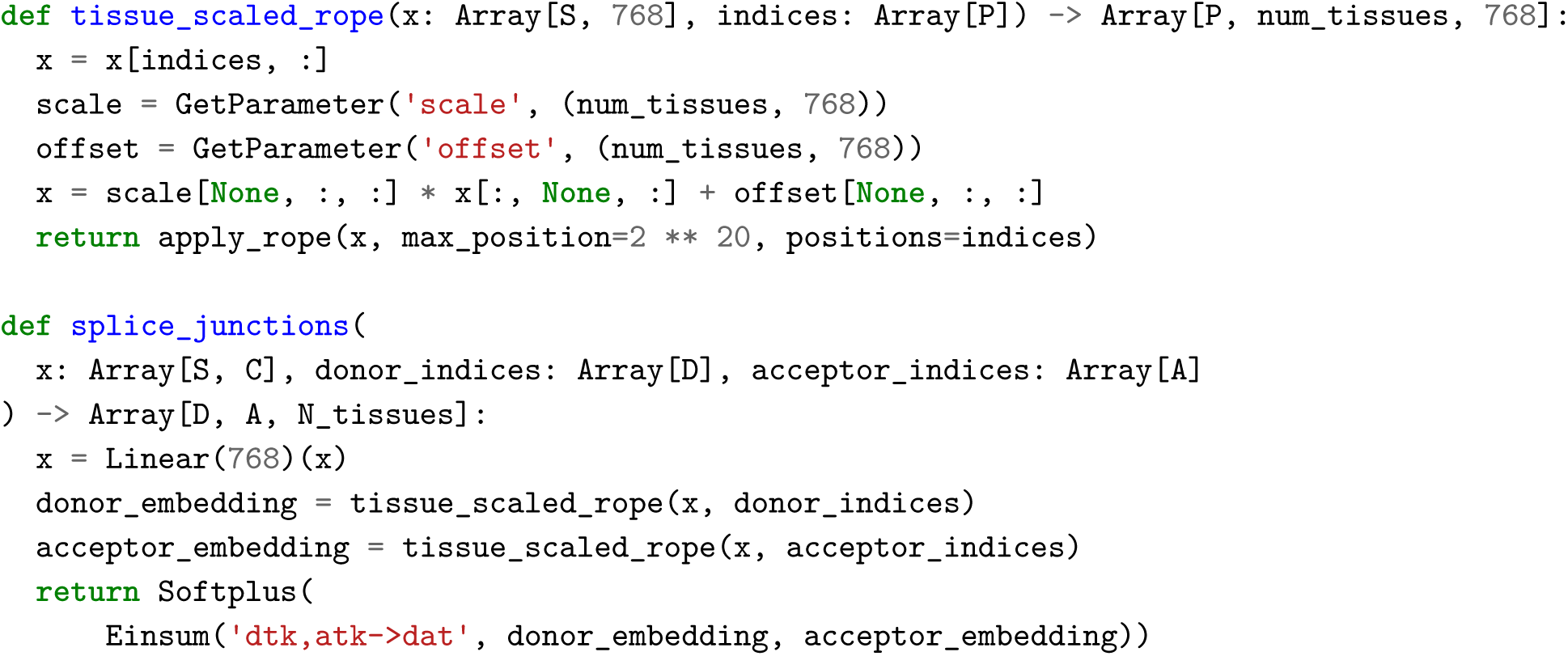

First, a linear layer projects the embeddings to an intermediate dimension. For sequence-level pre-training and evaluation (as described in this section), the tissue-specific genomic coordinates of expressed donor (donor_indices) and acceptor (acceptor_indices) sites, derived from the data processing pipeline, are provided as input. During distillation, these sites are obtained by selecting the top-k (with 𝑘 = 512) highest probability sites from the teacher’s splice site classification head. The procedure for identifying splice sites differs during variant effect prediction, as described in the Variant

Scoring section. The projected embeddings corresponding to these ground truth site coordinates are then processed by the tissue_scaled_rope function. This function first extracts the embeddings at the provided site locations and then applies the modified RoPE implementation described previously. Because splice sites are not contiguous, the function uses relative genomic coordinates as positions to incorporate distance information accurately. The modification to RoPE’s frequency calculation is particularly relevant here, reducing density at short ranges to better focus on the longer distances typical for splice junctions. Finally, the function applies a learnable, tissue-specific affine transformation (with distinct parameters per site type, strand, and organism). The main splice_junctions function computes the predicted count for each potential junction via an inner product (Einsum) between the corresponding processed donor and acceptor embeddings, followed by a Softplus activation to ensure positive values.

The training loss for splice junction predictions (junctions_loss) combines multiple terms. Two cross-entropy terms compare predicted and target count ratio distributions. These ratios represent conditional splicing probabilities. The first term evaluates the distribution of acceptor site usage for each donor, akin to a Percent Spliced In from the 5’ site (PSI5) perspective. This is achieved by normalizing counts over all acceptors for each donor (i.e., 𝑎𝑥𝑖𝑠 = 1, effectively 𝑃 (Acceptor *|* Donor, Tissue)).

The second term evaluates the distribution of donor site usage for each acceptor, akin to a PSI3 perspective. This normalizes counts over all donors for each acceptor (e.g., axis=0, effectively 𝑃 (Donor *|* Acceptor, Tissue)). Additionally, two Poisson loss terms compare the marginal sums: one compares the total predicted and target counts summed across all acceptors for each donor (e.g., 𝑎𝑥𝑖𝑠 = 1), and the other sums across all donors for each acceptor (e.g., 𝑎𝑥𝑖𝑠 = 0). For numerical stability of the Poisson loss term calculation, the sums of the target counts undergo an additional soft_clip operation. The final loss is a weighted combination of the cross-entropy and Poisson components. This entire loss calculation is performed independently for positive and negative strand predictions using their respective splice sites and targets. It is specified by the pseudocode:

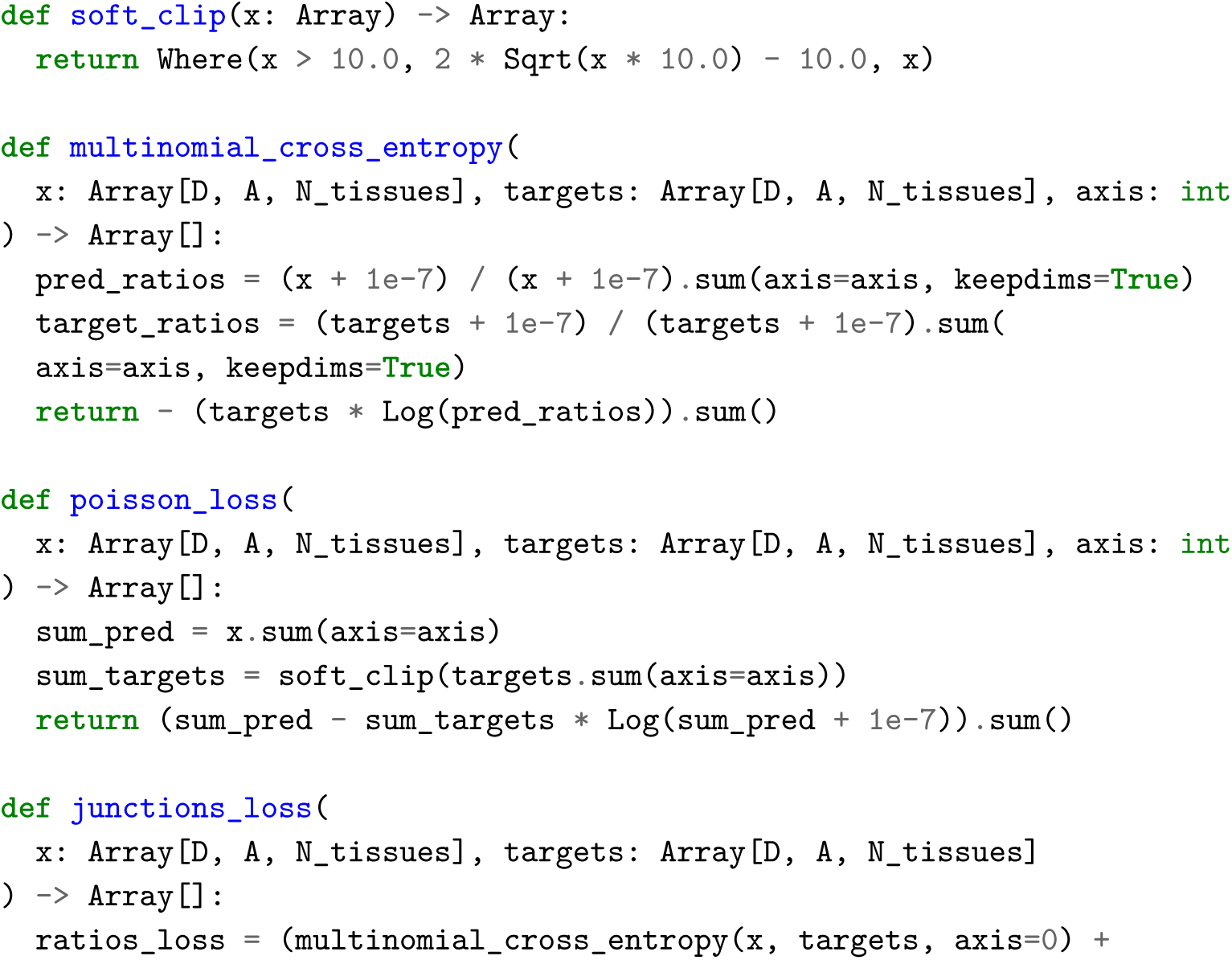

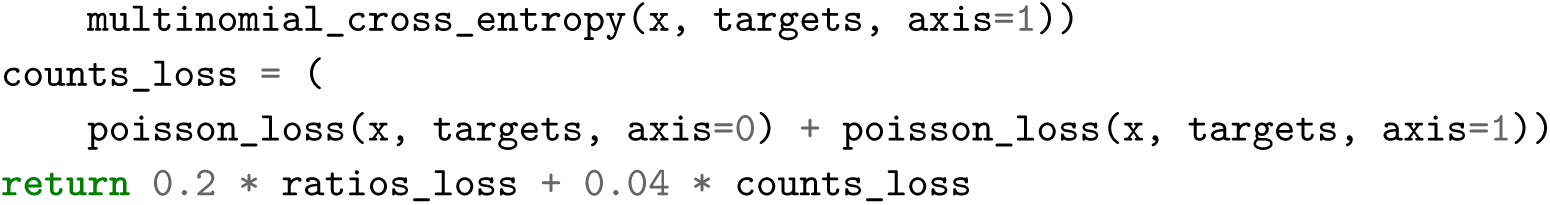

### Training

AlphaGenome was trained in a multi-task setting to jointly predict genomic features across both human and mouse reference genomes. We use two training strategies: the first is pretraining, where a model learns from experimental data, and the second is distillation, where a model is trained to reproduce the predictions of an ensemble of pretrained models. Model architecture is unchanged between training regimes.

#### Pretraining

When training from data, at every step we sampled contiguous 1 Mb intervals from the training set along with the corresponding experimental data for all model outputs. The training, validation, and test sets for these pretraining runs strictly followed the Borzoi-defined cross-validation folds. Borzoi’s original target intervals of 196Kb were extended to 1 Mb input windows for AlphaGenome, and any validation or test intervals that overlapped with training windows after extending were excluded (see Training Data).

We employed two forms of data augmentation during training. First, we applied a shift augmentation, where each sampled interval was shifted by a distance sampled uniformly at random from -1024 to +1024 bp. Second, reverse complementation was applied to the input sequence and corresponding outputs with a 50% probability.

**Sequence Parallelism** Pretraining utilized TPU v3 hardware, employing sequence parallelism to distribute the computation for each 1 Mb input interval across 8 TPU cores. AlphaGenome’s architecture, which alternates between local convolutional processing at high resolutions and global transformer-based processing at lower resolutions, is particularly amenable to this strategy.

For the sequence encoder, the 2^20^ bp input (1 Mb) is divided into 8 sub-sequences, each of length 2^13^ bp. To handle convolutional operations efficiently with minimal cross-device communication, each sub-sequence is extended with a 1024 bp copy of sequence from its neighbors on both sides before being processed by its assigned TPU core. After the encoder converts the input to 128 bp resolution embeddings, 8 embedding vectors (corresponding to the 1024 bp overlap) are trimmed from each end of the processed sub-sequences. This overlap/trimming strategy ensures that the encoder’s receptive field is accounted for (each 128 bp output bin embedding vector is influenced by an additional context of 513 bp to either side) and yields results nearly identical to non-parallel execution. A minor deviation is caused by batch normalization, which requires communication across all devices to aggregate statistics and slightly alters results due to double-counting the overlapping regions (2048 bp from each subsequence are double counted in the variance calculation). However, we observed no practical impact on model performance from this effect.

Processing the transformer tower and pairwise blocks under sequence parallelism requires more inter-core communication compared to the encoder’s convolutional layers. For instance, computing multi-query attention does not require communicating the queries (𝑞) but requires gathering the key (𝑘) and value (𝑣) tensors from all 8 cores onto each core. The pairwise activations are distributed along only the first sequence axis; this allows optimizations, such as only gathering the 𝑘 tensor (and not the 𝑞 tensor) during the sequence_to_pair operation.

Sequence parallelism for the sequence decoder mirrors the encoder’s strategy but operates in reverse. The 128 bp resolution embeddings entering the decoder on each core are first extended by concatenating 8 embedding vectors (corresponding to 1024 bp) from neighbors on each side. After the decoder upsamples these to 1 bp resolution, the overlapping regions (1024 bp on each side) are removed from the final output sub-sequences. Subsequent track prediction heads operate independently on the 1 bp output of each core, and the final loss computation and gradient updates are aggregated across all devices, akin to standard data parallelism.

**Model Training Parameters** In pretraining and distillation for the fold-specific and *all-folds* models, we trained the model using the AdamW^83^ optimizer with default hyperparameters (𝛽_1_ = 0.9, 𝛽_2_ = 0.999, 𝜖 = 10*^−^*^8^) and a weight decay of 0.4. Each gradient step processed a batch size of 64 samples using 8-way sequence parallelism, requiring 512 TPUv3 cores, with pre-training runs typically completing in approximately 4 hours. Training proceeded for a fixed duration of 15,000 steps without early stopping. The learning rate followed a schedule with a linear warmup from 0 to 0.004 over the first 5,000 steps, followed by a cosine decay to 0 over the remaining 10,000 steps. The number of steps was selected to balance expected model performance across both reference genome prediction tasks and zero-shot variant effect prediction tasks, as evaluated on validation data subsets.

Using this protocol, we trained a model for each of the four cross-validation folds (*fold0*, *fold-1*, *fold-2*, and *fold-3*), evaluated on their respective held-out reference genome validation sets. Following best practice, final test set performance was assessed only once, after all dataset choices and model training runs were finalized. Additionally, 64 models were trained using all available reference genome intervals (*all-folds*). As these models lack a dedicated reference genome holdout set, they were not used for track prediction evaluations, and they serve as the teacher ensemble for distillation.

#### Distillation

Model distillation begins with the same 1 Mb interval sampling and 50% reverse complement augmentation strategy used during pretraining. At this stage, we introduce additional input perturbations to increase the diversity of input sequences. Firstly, 4% of the nucleotides in each input sequence are randomly mutated by substituting with a nucleotide chosen uniformly at random. Secondly, we apply structural variations: insertions (using random sequences), deletions, and inversions. The number of such structural variations per 1 Mb sequence is sampled from a Poisson distribution (𝜆 = 1.0), and the length of each variation is chosen uniformly from the range [1, 20] base pairs.

Distillation training was performed without sequence parallelism across 64 NVIDIA H100 GPUs, with a batch size of 64 (effectively one sample per GPU). Each GPU loaded a different frozen teacher model from the pool of 64 pretrained *all-folds* models. For each GPU, an interval is randomly sampled and predictions were generated using its assigned frozen teacher model. The student model’s parameters were replicated across all devices and it was then trained using the teacher predictions as targets by minimizing the same set of loss functions used during the pretraining phase. By averaging gradients computed against different teachers, this setup provides a computationally efficient stochastic approximation of training the student against the expected predictions of the full teacher ensemble, without needing to run all teachers for every sample.

**Model Training Parameters** The distillation process ran for 250,000 steps, taking approximately 3 days. We used the AdamW optimizer with default parameters and a reduced weight decay coefficient of 0.04 compared to pretraining. The learning rate schedule consisted of three phases: a linear ramp-up to 0.002 over the initial 5,000 steps, a constant phase at 0.002 for the next 120,000 steps, and finally a cosine decay down to 0 over the concluding 125,000 steps.

### Benchmarking Against Existing Methods

AlphaGenome’s performance was benchmarked against several existing state-of-the-art sequence-to-function models. These included general-purpose models like Enformer^2^ and Borzoi^3^, as well as models specialized for specific tasks: ProCapNet^8^ for PRO-cap signal prediction; SpliceAI^5^, DeltaSplice^84^, Pangolin^11^, and Splam^85^ for splicing predictions; Orca^19^ for contact map prediction; and ChromBPNet^10^ for local chromatin feature prediction. For these comparisons, evaluations were performed on held-out human test intervals, using metrics appropriate for each prediction task. Pearson correlation coefficient (Pearson *r* ) was used for quantitative coverage-based tracks, while the Area Under the Precision-Recall Curve (auPRC) was used for binary classification tasks (details and full results in Supplementary Table 3 and Supplementary Table 4).

Publicly available pre-trained models, weights, or source code for these comparison models were utilized as explained in Supplementary Table 8. Specific considerations were applied for certain comparisons to ensure fair evaluation. When scoring variants with SpliceAI, Pangolin, and DeltaSplice, predictions were made by considering a 2,000 base pair window on either side of the variant, overlapped with the gene mask for this region. For performance comparisons between AlphaGenome and ChromBPNet, evaluations were conducted exclusively on experimental tracks with directly matching ENCODE accessions. For ATAC, these were ENCSR200OML (IMR-90) and for DNase: ENCSR000EOT (K562), ENCSR149XIL (HepG2), ENCSR000EMT (GM12878) and ENCSR477RTP (IMR-90).

### Genome Track Evaluation

We evaluated AlphaGenome’s ability to predict various genomic features using several metrics tailored to the nature of each output type.

#### Correlation for Continuous Tracks

Concordance between predicted and observed continuous track signals (such as those for ChIP-seq, DNase-seq, ATAC-seq, CAGE, and PRO-cap) was primarily measured using the Pearson correlation coefficient (r). For a given track and a held-out test interval, Pearson r was calculated between the vector of predicted values and the vector of observed values across all corresponding genomic bins within that interval. The distributions shown in Fig. 2c represents the Pearson r values calculated for all tracks within a specific assay group (e.g., all TF ChIP-seq tracks) and organism (e.g., human and mouse) across all held-out test intervals. The average Pearson r for each group (shown as text and circle) is the mean of these individual track correlations.

#### Gene Expression Correlation

Pearson correlation (r) was also computed between observed and predicted gene expression values on held-out test intervals. Gene expression values for both observed and predicted tracks were calculated as the log-transformed mean read coverage across all annotated exons for a given gene, using GENCODE version 46 annotations and considering strand matching. To avoid duplicate genes across test intervals, we only considered genes with at least 50% of their exons falling within a test interval. Three specific correlation approaches were used:

- *Raw; Across Genes (Fig. 2d, left):* For each cell type/track, correlation was computed across all genes between their predicted and observed log-transformed expression values.
- *Normalized; Across Genes (Fig. 2d, middle):* For each track, log-transformed expression values were quantile normalized across genes. Then, for each gene, its mean expression across all tracks was subtracted. Correlation was then computed across genes between these normalized predicted and observed values within each track.
- *Normalized; Across Tracks (Fig. 2d, right):* Using the same quantile-normalized, gene-mean-centered data as above, correlation was computed *across* tracks/cell types for each gene separately, assessing per-gene prediction consistency over different cellular contexts.

#### Alternative Polyadenylation

To assess the model’s understanding of alternative polyadenylation site usage, we compute polyadenylation- centric coverage ratios (COVR), as defined in the Borzoi paper. These COVR values quantify the relative usage of the distal versus proximal polyadenylation sites for a gene as annotated by PolyADB^86^. We compare AlphaGenome’s and Borzoi’s predicted COVR values on held-out test sites against the true COVR values derived from tissue-pooled GTEx data.

#### Splicing Prediction Performance

AlphaGenome’s performance on various splicing-related track prediction tasks was assessed using specific test sets, metrics, and comparisons depending on the specific output type.

**Splice Site Classification** The model’s ability to correctly identify and classify splice sites was primarily evaluated using the auPRC metric. For this task, we evaluated separately with true labels derived from RNA-seq observed splice sites and those annotated in GENCODE GTF files. At each relevant genomic position, the model predicted probabilities for four positive splice site classes: Donor site on the plus strand (Donor+), Acceptor site on the plus strand (Acceptor+), Donor site on the minus strand (Donor-), and Acceptor site on the minus strand (Acceptor-). The test sequences were split into batches with input sequence length 1M base pair. A separate auPRC was computed for each of these four classes, comparing predicted probabilities against true labels, and the overall reported performance is the average auPRC across them.

For specific comparisons against models like SpliceAI and DeltaSplice, AlphaGenome’s performance (using the ensemble of four *fold-1* models, chosen for maximal test set overlap with these peer models) was assessed on test intervals derived from human chromosomes 1, 3, 5, 7, and 9 for consistency. Pangolin was not evaluated for this task as it does not predict separately donors or accepters, only splice sites in general.

**Splice Site Usage Prediction** The evaluation of splice site usage (SSU) was conducted using Pearson correlation coefficients (r). These were computed between the vector of AlphaGenome’s predicted SSU values and the vector of observed SSU values (derived from RNA-seq) for each tissue across held-out test intervals, treating each tissue’s SSU profile as a distinct track.

For comparative benchmarking (e.g., against DeltaSplice), the same held-out test intervals and SSU predictions FROM AlphaGenome *fold-1* were used. Annotated SSU values were derived from RNA-seq. Comparisons with Pangolin for SSU were not performed due to differing SSU definitions and due to Pangolin not using its SSU prediction head for variant effect prediction.

**Splice Junction Prediction** The model’s performance in predicting splice junctions was evaluated through three distinct approaches on the held out test intervals of *fold-1* with an ensemble model trained with *fold-1* training data.

1. *Classification of true vs. false junctions:* This task assessed the model’s ability to distinguish true junctions (defined as donor-acceptor pairs with supporting RNA-seq read counts after the filtering steps described in the data section) from the vast majority of false junctions (defined as donor-acceptor pairs with no RNA-seq supporting). This task is challenging since only a small set

of donor/acceptor pairs are true within 1M base range. The model’s predicted junction counts were the classification scores. For Splam, since it does not directly predict a junction score, we used the min of the donor and acceptor splice site probabilities as a prediction for the junction score for each splice junction. An auPRC was computed for each tissue independently by comparing flattened matrices of these predicted scores against the corresponding flattened binary ground truth labels. The final reported performance is the average auPRC across all tissues.

**1.** 2. *Quantitative prediction of junction counts:* The accuracy of predicting the strength of junction usage was assessed using the Pearson correlation between log(1 + 𝑥) transformed predicted junction counts and log(1 + 𝑥) transformed measured junction counts (interpreted as junction strength).

This correlation was only computed over junctions that had non-zero read counts in the ground truth test set data.

*Prediction of PSI5 and PSI3 levels:* The model’s ability to predict local splicing choices was further evaluated by comparing measured and predicted PSI5 and PSI3 levels using Pearson correlation. These metrics are defined as: PSI5(D,A)=n(D,A) /A’n(D,A’), PSI3(D,A)=n(D,A) /D’n(D’,A), where n(D,A) is the number of split reads supporting the specific junction from donor D to acceptor A, A’n(D,A’) is the total number of split reads supporting all splice junctions originating from donor D, and D’n(D’,A) is the total for all splice junctions ending at acceptor A. The PSI correlation reported in Extended Data Fig. 2 are from the test intervals in chr2 only.

Together, these three approaches provide a multifaceted assessment of AlphaGenome’s ability to accurately predict both the presence and quantitative strength of splice junctions, as well as local splice isoform choices.

#### Contact Maps Performance

Our evaluation of contact map predictions was designed to closely mirror the evaluations performed on the Orca 1 Mb model^19^. The primary metric for assessing performance was the mean Pearson correlation coefficient (r), where individual Pearson r values were calculated between AlphaGenome’s predicted contact scores and the observed experimental contact scores within each held-out genomic interval in a specific human tissue type. The held-out test intervals were defined by intersecting the Orca test split (human chromosomes 9 and 10) with AlphaGenome’s cross-validation splits (specifically, those aligned with Borzoi FOLD_0). Human tissue types chosen for this evaluation matched key training samples from the Orca study, specifically the H1-hESC and HFFc6 Micro-C datasets.

To align with Orca 1 Mb model’s headline evaluations, genomic intervals for comparison were 1,000,000 bases (1 Mb) in length, divided into 250 bins of 4 kb each (the native output resolution of Orca). Since AlphaGenome’s native output resolution for contact maps is 2048 bp bins over a 1,048,576 bp interval, AlphaGenome’s predictions were resized and interpolated to match Orca’s 4 kb resolution using bilinear interpolation (jax.image.resize).

Furthermore, the H1-hESC - HFF Micro-C *difference* prediction evaluation was re-implemented following the specific protocol detailed in the Orca code repository (Cell 20 of https://github.com/jzhoulab/orca_manuscript/blob/main/Model_performance.ipynb). This difference analysis was performed over the same held-out chromosomal splits mentioned above.

#### Benchmarking Against Enformer Track Predictions

To facilitate a direct and fair comparison with Enformer^2^, a specialized AlphaGenome model was trained using the identical training sequence intervals as the original Enformer model. This AlphaGenome variant incorporated a dedicated “Enformer head” to predict Enformer’s target tracks at their native 128 bp resolution (while the model’s other heads predicted standard AlphaGenome outputs). For this Enformer-specific head, loss was calculated exclusively over the central 114,688 bp region corresponding to Enformer’s target width, using AlphaGenome’s full 1 Mb input. To prevent data leakage, test set intervals whose 1 Mb AlphaGenome input windows overlapped these Enformer training intervals were excluded from evaluation.

#### Benchmarking Against Borzoi Track Predictions

For a direct comparison with the Borzoi *fold-1* model^3^, an AlphaGenome model, which was initially trained on the identical Borzoi *fold-1* data split, underwent fine-tuning. This fine-tuning process involved augmenting the AlphaGenome architecture with two additional heads to mirror Borzoi’s outputs.

The first head aggregates AlphaGenome’s 1 bp embeddings into 32 bp embeddings and makes predictions matching the Borzoi tracks (7,611 human and 2,608 mouse). This head is trained on Borzoi’s TFRecords dataset (original resolution and scaling) to allow for a direct comparison with the published Borzoi model. This head is used for metrics reported in Fig. 1d where we compare against Borzoi at 32 bp resolution.

The second RNA-seq head is trained on the same RNA-seq tracks as Borzoi but reprocessed at 1 bp resolution and without any Borzoi specific scaling. When comparing this head against Borzoi, we unscale and repeat Borzoi’s predictions 32 times (to equal 1 bp unscaled predictions). This head is used for metrics reported in Fig. 1d where we compare against Borzoi at 1 bp resolution.

We validated the approach of upsampling and scaling the additional RNA-seq head to match Borzoi by applying the same procedure to the training data. AlphaGenome’s base-resolution data was aggregated to a 32 bp resolution and Borzoi’s original scaling methodology was applied. Compared to Borzoi’s provided TFRecord files, we achieved high concordance (0.988 average Pearson r correlation) . This comparison excluded unmappable regions, as flagged by the ‘umap’ entry in the Borzoi data examples. Finally, in the Borzoi’s and AlphaGenome’s RNA-seq comparison at 1 bp resolution, we only include the tracks for which the Pearson r correlation is larger or equal than 0.99 and exclude the unmappable regions.

#### Relative Improvement Metric

To contextualize performance gains, particularly when comparing AlphaGenome against various models and random baselines across different tasks, we calculated a Relative Improvement score (as shown in Fig. 1d for tracks and Fig. 1e for variant effect prediction). This metric is defined by the formula:

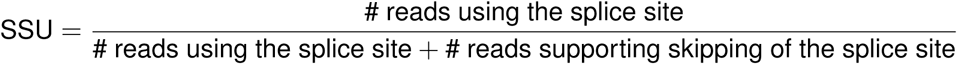

In this formula:

- alphagenome metric refers to the performance score (e.g., auPRC, Pearson *r* ) achieved by AlphaGenome.
- comparison_model_metric is the performance score of the specific model AlphaGenome is being compared against (e.g., Borzoi, Enformer) on the identical task and dataset.
- random_baseline_metric represents the expected performance of a random predictor, chosen appropriately for the primary evaluation metric being used.

Random baseline metrics were computed as follows:

- For tasks evaluated with either Pearson or Spearman correlation, the random_baseline_metric was set to 0.
- For tasks evaluated with auPRC, the random_baseline_metric was set to the fraction of positive instances in the dataset (i.e., the class prevalence).
- For tasks with class-balanced datasets evaluated with auROC, the random_baseline_metric was set to 0.5.

This relative improvement metric normalizes the gain achieved by AlphaGenome over a comparison model by the amount that comparison model itself improved over a random baseline. Detailed individual comparisons that contribute to these relative improvement calculations are presented in Supplementary Table 3 and Supplementary Table 4.

For accessibility variants, we evaluated AlphaGenome and the SOTA models on a number of highly similar tasks, specifically five directionality benchmarks and three causality benchmarks, as described in the section Chromatin accessibility variants & bQTLs. We outperform SOTA in each individual case. However, to present a consolidated number in Fig. 1, we computed the average relative improvement across these benchmarks.

#### Variant evaluations Variant scoring

Variant scoring aims to distill AlphaGenome’s multifaceted genomic predictions into a single, informative scalar value for each genetic variant, representing its predicted biological impact. This score quantifies the difference in predicted features (e.g., chromatin accessibility, TF binding, gene expression, splicing outcomes) between sequences carrying the reference (REF) and alternative (ALT) alleles. The methodology for deriving this score is critical, involving choices about the genomic region analyzed, the function used to aggregate allelic differences or activity levels across this region, and any applied transformations (e.g., log ratios).

Based on optimization against downstream evaluation tasks, we established a set of 19 recommended variant scoring configurations covering diverse genomic modalities (details in Supplementary Table 9, with illustrated workflows in Supplementary Fig. 12). These scorers primarily fall into two categories – those using masks centered on the variant (*center mask scorers*) and those using masks derived from gene annotations (*gene mask scorers*) – supplemented by specialized scorers for features like contact maps and specific splicing or polyadenylation events.

#### Scoring pipeline overview

Before detailing scorer-specific configurations, we first outline the overall pipeline for center mask and gene mask variant scorers.

**1. Allele-Specific Predictions** AlphaGenome generates two sets of genomic predictions centered on the variant: one for the DNA sequence containing the REF allele and another for the sequence with the ALT allele. The REF allele sequence used for comparison is either the standard reference genome allele or, when specified, the specific REF allele defined by the evaluation dataset.
**2. Indel Alignment (Optional)** Following previous work^5^, for insertion or deletion (indel) variants , the ALT allele’s prediction profile is aligned to the REF allele’s coordinate space. Inserted bases are summarized by taking the maximum value over the inserted segment, while deleted bases are treated as having zero signal in the ALT context, thereby enabling consistent positional comparisons.
**3. Mask Application and Spatial Aggregation** A spatial mask is applied to focus on the relevant genomic regions within the 1 Mb input window.
a. For **center-mask approaches**, a fixed-width window (e.g., 501 bp for local marks like ATAC-seq or DNase-seq; 2001 bp for broader histone modifications) is defined around the variant.
b. For **gene-mask approaches**, windows for each gene are defined by gene annotations (e.g., gene bodies, exons, or TSS locations from a GTF file).

Predictions across the spatial windows are then aggregated using sum, mean, L2 norm, or taking the maximum across all REF and ALT signals. This results in a single output per track (center-mask) or per track and gene (gene-mask) for both the REF and ALT allele.

1. **4. Transformation and Allelic Comparison** Mathematical transformations (e.g., log, log1p, absolute value) may be applied to stabilize variances or emphasize particular types of effects, either before or after spatial aggregation, depending on the specific scorer configuration. Finally, the REF and ALT predictions are compared to yield a final score per track (center-mask) or per track and gene (gene-mask).
2. **5. Track Aggregation (Optional)** After a score is derived for each output track (e.g., per cell type or assay target), a final step may aggregate these scores (e.g., by taking the maximum absolute score across relevant tracks or selecting a specific track of interest) to produce a single scalar value representing the variant’s overall predicted impact for that specific scoring configuration.

**Core Differential Variant Scorers** We provide the following core scorers for quantifying the impact of variants by comparing REF and ALT allele predictions:

- *Gene expression (using RNA_SEQ outputs):* To score effects on RNA abundance (from RNA_SEQ model outputs), predictions are aggregated over annotated gene regions (mean signal across exons). The variant score is the log-fold change of this aggregated gene expression level between the ALT and REF alleles (Supplementary Fig. 12b). Only genes fully contained within the input interval are scored.
- *Splicing – Site-based Impact (SPLICE_SITES AND SPLICE_SITE_USAGE outputs):* Variant effects on predicted splice site probabilities (from SPLICE_SITES outputs) or usage percentages (from SPLICE_SITE_USAGE outputs) are evaluated within gene boundaries. The score is the maximum absolute difference in these predicted values between the REF and ALT alleles across all positions within a gene body (Supplementary Fig. 13a). Only genes overlapping the variant are scored.
- *Splicing – Junction-based Impact (SPLICE_JUNCTIONS output):* To assess effects on specific splice junctions, scores are derived from the change in predicted junction counts or strengths between REF and ALT alleles, after a log transformation. Splice junction prediction for the REF and ALT are based on the same set of splice sites. In contrast to the model training stage, the splice sites used are a union of those observed from RNA-seq data (described in splice site training data), those predicted from the reference sequence, and those predicted from the alternative sequence. The splice site probability is an element-wise maximum of the three sources (RNA-seq splice sites are labeled as probability 1 to ensure they have the highest priority). For indels, an alignment step is performed before taking the element-wise maximum. Only splice sites within the gene body of the genes overlapping the variant are considered. For each variant, we consider a maximum of 256 donor or acceptor sites. To score the variant for each tissue, the maximum absolute log fold change of the junction score per junction across all junctions within a gene is reported (Supplementary Fig. 13b).
- *Polyadenylation site usage (using RNA_SEQ outputs):* Variant effects on polyadenylation (from RNA_SEQ model outputs) are scored using a method analogous to Borzoi’s paQTL approach. This gene-based score compares predicted RNA coverage at annotated 3’ cleavage junctions for REF and ALT alleles. It reflects the maximum change in relative usage between any two potential polyadenylation sites (one considered proximal, one distal) within a gene’s 3’ UTR, expressed as an absolute log2 fold change of these isoform ratios (Supplementary Fig. 12c). Only genes with 80%+ of their polyadenylation sites within the input interval are scored.
- *Local chromatin accessibility and activity (using ATAC, DNASE, CAGE, PRO_CAP outputs):* For assays measuring local chromatin state or transcriptional start site (TSS) activity, variant effects are assessed using a mask centered on the variant (501 bp). The score represents the log2 fold change of total signal summed within the window for the ALT allele versus the REF allele (e.g., log2((sum(ALT) + p) / (sum(REF) + p)), Supplementary Fig. 12a).
- *Transcription factor binding and histone modifications (CHIP_TF and CHIP_HISTONE outputs):* For these assays, variant effects are assessed using a 501 bp centered window for CHIP_TF and 2001 bp for CHIP_HISTONE to account for the broader nature of some histone marks. The score represents the log2 fold change of summed ALT vs. REF signals.
- *Contact Maps (CONTACT_MAPS output):* For variants affecting 3D chromatin contacts, a method similar to that used by Orca is employed for SNVs. This calculates the mean absolute difference between REF and ALT contact map predictions for all interactions involving the single genomic bin containing the variant and other bins within a defined local window (e.g., 1 Mb).

This suite of scoring strategies allows for a multifaceted interpretation of variant effects across various genomic functions predicted by AlphaGenome.

**Active Allele Scoring** In addition to the differential scores described above, we also provide scoring configurations that capture the *absolute activity level* associated with an allele, rather than quantifying the change between REF and ALT. This is calculated by taking the maximum of the aggregated signals from the REF and ALT alleles over the central window (center-mask scorers) or gene region (gene-mask scorers). Active allele scorers provide insight into the overall strength of a regulatory element harboring the variant by reporting the maximum of the summed signals for the REF and ALT alleles within the defined centered window.

**Composite splicing variant scorer** To provide a comprehensive measure of a variant’s overall impact on splicing, predictions from the splice junction, splice site, and splice site usage heads are integrated. Splice site and usage scores inherently range from 0 to 1. Splice junction scores, representing absolute log-fold changes, predominantly exhibit values up to approximately 5 in typical scenarios. Therefore, to ensure comparable contributions from each component, the splice junction scores are first normalized by dividing by 5. The final aggregated splicing score is then the sum of this normalized junction score and the original splice site and usage scores.

#### Calibration Methodology

To make raw variant scores more interpretable and comparable across different assays and genomic contexts, we implemented an empirical quantile calibration procedure against a background set of common genetic variants. This variant set comprised 348,126 common human SNPs from chromosome 22 with MAF>0.01 in any of gnomAD’s ancestral groups (gnomAD v3). We used this set of variants to estimate a background score distribution for each variant scorer and track. We can then derive a ‘quantile score’ for any arbitrary variant (using the procedure below), representing its percentile rank (or signed percentile rank) within the distribution of scores from common variants for a given scorer and track. This provides a measure of predicted impact that is standardized to the same scale across variant scorers and tracks. We used common variants as they are more likely to be depleted for high-impact variants compared to random genomic variants, and are less likely to be biased to a particular use-case or modality.

The calibration process was as follows:

1. *Raw score generation:* The full set of 19 recommended AlphaGenome variant scorers was run on these 348,126 background variants to generate their raw effect scores.
2. *Quantile computation:* For each unique combination of scorer and output track (e.g., a specific epigenetic mark in a particular cell type), empirical quantile probabilities were computed from the distribution of raw scores. This was achieved by stratifying the raw scores into 1,000 bins. To enhance precision at the extremities of the score distributions (i.e., for very strong or very weak predicted effects), the spacing of these bins was determined using a sigmoid function.
3. *Handling of invalid scores:* Any Not-a-Number (NaN) scores were ignored during this process. NaN scores arise in scenarios such as when a gene-based scorer is applied to a variant outside of any relevant gene mask (e.g., a splicing scorer applied outside of a gene mask, such as for an intergenic variant) or when predictions are made for genes on the incorrect strand relative to the scorer’s expectation.
4. *Minimum thresholding:* Quantile probabilities were thresholded at 1 *×* 10*^−^*^5^ (e.g., values below this

were floored to this minimum) to mitigate the impact of extreme outliers, and due to the number of variants in the background set, 𝑂(1𝑒5).

1. 5. *Adjustment for signed scores:* For signed variant scores (which indicate effect direction like up-

regulation or down-regulation), their [0,1] quantile probabilities – derived directly from the rank order of the original signed raw scores – are linearly transformed to a [-1,1] range. This rescaling ensures the calibrated score reflects the original directionality: for instance, the 0th percentile (representing the most negative raw scores) maps to -1, the 50th percentile (raw scores around zero) to 0, and the 100th percentile (most positive raw scores) to +1.

1. 6. *Tie-breaking:* Occasionally, multiple background variants yielded identical raw scores, such that the same score would map to multiple quantiles. To enforce unique ranks, a tie-breaking procedure was applied: variants in a tied group were randomly (with uniform distribution) assigned a distinct quantile score, spanning the percentile range they collectively occupy.

#### *In Silico* Mutagenesis (ISM) for Contribution Scores

To interpret which nucleotides in a sequence of interest contribute most to AlphaGenome’s predictions for a specific genomic feature, we employed an *in silico mutagenesis* (ISM) approach. This method estimates a contribution score for each position in the sequence.

The procedure is as follows:

1. *Systematic single nucleotide variant (SNV) generation:* For each position within the 𝐿-length input sequence, all three possible alternative nucleotide substitutions are generated. This creates a comprehensive set of SNVs covering every possible single base change from the original sequence.
2. *Variant scoring:* Each of these generated SNVs is then scored using a chosen AlphaGenome variant scorer (as described in the Variant Scoring section) to obtain a single scalar value representing the predicted impact of that specific mutation on a particular model output (e.g., an epigenomic track or gene expression value in a specific cell type).
3. *Construction of an effect matrix:* The resulting scalar scores for the three alternative mutations at each position are organized into an 𝐿 *×* 4 matrix, where 𝐿 is the sequence length and the 4 columns correspond to the four possible nucleotides (A, C, G, T). For each position, the entries for

the three alternative (mutant) nucleotides are populated with their respective variant scores, while the entry for the original reference nucleotide at that position has a score of 0 (as it represents the baseline against which mutations are compared).

**1.** 4. *Mean-centering:* To normalize these effects, for each position 𝑖 in the sequence, the mean of the three variant scores (i.e., the scores associated with mutating the original base to its three alternatives) is calculated. This mean is then subtracted from all four entries (A, C, G, T) in the matrix at that position 𝑖. Consequently, after this step, the matrix entry corresponding to the original

reference base at position 𝑖 now holds the value 0 *−* mean_of_alternative_scores.

**1.** 5. *Computing the final contribution scores*. The final contribution score for each nucleotide in the original input sequence is then taken directly from this mean-centered 𝐿 *×* 4 matrix by selecting the value that corresponds to the actual reference base at that position. These 𝐿 resulting scores

(one per position) represent the influence of each reference base relative to the average of its alternatives and are typically visualized as a contribution score track or “saliency map”.

These per-position contribution scores are then visualized as sequence logos, where the height of each letter in the original reference sequence is scaled by its calculated contribution score. The value thus obtained for each position quantifies the specific contribution of the reference base to the predicted outcome, relative to the average effect of substituting it with alternative nucleotides.

Signed values are meaningful for two-sided scorers. For variant scorers producing strictly non-negative scores, we always report non-negative values corresponding to the mean of alternative scores.

**Comparative ISM for Variant Effect Interpretation** Furthermore, to specifically investigate how a genetic variant might alter local sequence motifs, this ISM procedure is applied independently to both the reference (REF) sequence and the sequence containing the alternative (ALT) allele. The resulting REF and ALT contribution score profiles (as sequence logos) are then compared. This comparison is powerful because the single nucleotide change in the ALT sequence can alter the model’s interpretation of the entire local region, not just the mutated base itself. This comparative approach is particularly useful for identifying if a variant potentially disrupts an existing regulatory motif present in the REF sequence or, conversely, creates a novel motif in the ALT sequence.

#### Test-time augmentation

A test-time augmentation (TTA) strategy involving strand averaging was employed when evaluating performance on variant and unaggregated track evaluations. Predictions were generated for both the forward DNA sequence and its reverse complement. For track evaluations, performance metrics were computed on the average of the two predictions. For variant evaluations, variant effect scores were computed independently based on each stranded prediction, and the final reported score for a variant was the average of these two. This approach generally improved the performance metrics observed for non-distilled models, but did not consistently enhance the performance of AlphaGenome’s distilled model evaluations.

#### Chromosome Splits for Variant Benchmarks

For zero-shot evaluations, each variant was assigned to either the validation or test set, according to the following chromosome split: variants in chromosomes 1, 2, 4, 5, 7, 8, 10, 11, 13, 14, 15, 17, 20, 22, X were used as the validation set to inform model design and for variant scoring strategy development, whereas variants in chromosomes 3, 6, 9, 12, 16, 18, 19, 21 were used as the test set, on which all final performance metrics are reported.

For supervised evaluations, we used the following split: chromosomes 1, 4, 7, 8, 10, 13, 15 were used as the training set, chromosomes 2, 5, 11, 14, 17, 20, 22, X as the validation set, and chromosomes

3, 6, 9, 12, 16, 18, 19, 21 as the test set. Supervised models were developed on the basis of the training and validation set, with final performance metrics reported on the test set.

#### Splicing Variant Benchmarks

To assess AlphaGenome’s ability to predict the functional consequences of variants on splicing, we utilized several curated datasets and benchmarks, each focusing on different aspects of splicing regulation. Different models were evaluated on these benchmarks using suitable variant scorers corresponding to their supported output types. AlphaGenome was evaluated using composite splicing scores and splice junction scores. Borzoi was evaluated using RNA-seq scores computed as the maximum absolute difference in normalized coverage across the gene span, as in the original study^3^. SpliceAI was evaluated using splice site prediction scores. DeltaSplice and Pangolin were evaluated using splice site usage scores.

**1. Splicing Quantitative Trait Loci (sQTLs)** Splicing QTLs (sQTLs) are genetic variants that influence gene splicing, often by altering isoform abundance. We utilized a curated dataset previously developed for the Borzoi model based on the eQTL catalogue^3,87^. This dataset comprises 21,514 fine-mapped causal sQTLs and tissue pairs (positive examples) and 21,514 carefully selected distance-matched non-sQTL variants and tissue pairs (negative controls) across 49 GTEx tissues.

The task was to distinguish between these causal sQTLs and the matched negative controls, evaluated using auPRC per tissue.

1. **2. ClinVar Variants** ClinVar variants were downloaded from ClinVar ftp portal (https://ftp.ncbi.nlm.nih.gov/pub/clinvar/vcf_GRCh38/, release date 20250323). Only variants that meet the following criteria were considered:
2. Annotated as one of ‘Pathogenic’, ‘Likely_Pathogenic’, ‘Benign’, ‘Likely_Benign’.
3. Has at least one review star.
4. In autosome
5. Annotated either intronic, synonymous, or missense

All ‘Pathogenic’, ‘Likely_Pathogenic’ variants are labelled as 1 and all ‘Benign’ and ‘Likely_Benign’ variants are labelled as 0. ClinVar variants are divided into three categories based on molecular consequences from the ClinVar VCF file and distance to the nearest splice site:

1. *Deep intronic and deep synonymous*: variants annotated as ‘intronic’ and >6 bp from the closest splice site, or variants annotated as ‘synonymous’ and >3 bp from the closest splice site. Since the majority of intronic and and synonymous variants are benign, a further sampling step was performance for ‘clinvar_splicing’. For each pathogenic/likely_pathogenic variant, up to 100 be- nign/likely_benign were sampled randomly. All benign/likely_benign were sampled for the gene if the negative variant is less than 100 fold of the positive variant. 1,628 positive and 95,269 negative variants are in this category.
2. *Missense variant* : all variants annotated as ‘missense’ and are predicted by AlphaMissense as ‘Likely Benign’. 5,108 positive and 100,031 negative variants are in this category.
3. *Splice site region*: intronic, synonymous, and missense variants that are less than 3 bp from splice sites for exonic variants or less than 6 bp from splice sites for intronic variants. 7,155 positive and 47,354 negative variants are in this category.
4. **Splicing Outliers from GTEx** Splicing outliers junctions are characterized by highly aberrant splicing patterns (e.g., exon skipping, intron retention, or cryptic splice site usage). Aberrant spliced splicing outliers and the associated rare variants were derived from GTEx RNA-seq data following the description in the AbSplice study^29^. Specifically, splicing outliers were detected with the FRASER 2.0^88^ and DROP

pipeline (version 1.3.3)^89^. Variant allele frequency was derived from GnomAD release 4.0 (accessed February 2024)^90^. Only GTEx variants with minor allele frequency less than 0.1% and that are present in at most two individuals are considered. Splicing outlier junctions are paired with rare GTEx variants matching the individual and require that the rare variant is at most 250 bp from the aberrant splice junction. We refer to the AbSplice study^29^ for more detailed data processing steps. This comprehensive dataset includes over 1 million rare variants of which 5,819 are splicing outliers in one or more GTEx human tissues, annotated for their potential to cause tissue-specific aberrant splicing.

On this benchmark, AlphaGenome was evaluated in two ways on the same test subset of variants:

- *Zero-shot evaluation:* Directly using AlphaGenome aggregated splicing scores and splice junction scores.
- *Supervised evaluation:* Training an Explainable Boosting Classifier model (from the InterpretML python package^91^) and evaluating on the held-out test subset of variants. Features for this classifier included splicing-related variant scores derived from AlphaGenome (scores from splice site, splice site usage, and splice junction predictions, as well as RNA-seq scores computed as the maximum absolute difference in normalized coverage across the gene span^3^). An additional binary feature, indicating whether a splice site is expressed in a given tissue (using a cutoff of 10 reads for the median number of split reads sharing the splice site from GTEx RNA-seq samples), was also included. As baseline, we used the AbSplice ensemble model^29^ re-trained and evaluated on the same dataset split as AlphaGenome.

The metric used was auPRC across all GTEx tissues, computed by assigning a tissue to each rare variant: positive variants were mapped to their corresponding GTEx tissue, while negative variants were randomly assigned a tissue according to the positive variants’ tissue distribution.

**1. 4. MFASS (Massively Parallel Assay of Splicing Sequences)** To evaluate predictions against high-throughput experimental measurements of splicing, we used data from the MFASS study^28^, a massively parallel reporter assay that tested the exon skipping effects of 27,733 ExAC single nucleotide variants (SNVs) spanning or adjacent to 2,339 exons. Data processing for the MFASS dataset followed the pipeline detailed in the MMSplice paper’s code repository (https://github.com/gagneurlab/MMSplice_ paper/blob/master/code/Figure2/MFASS.ipynb).

Model performance was assessed using auPRC for predicting whether a variant is a “splice disrupting variant” (SDV) or not. SDVs were defined in the MFASS study as variants that change the exon inclusion index by at least 0.5. Variants with mislabelled strand information were removed. The final dataset has 1,040 positive splice disrupting variants and 26,464 negative variants.

To score MFASS variants, we found that using predicted splice site logits instead of probabilities improved the performance of all models, therefore logits are used to score variants in this benchmark instead of probabilities as in other benchmarks. MFASS measures exon inclusion index. To score MFASS variants, the mean predicted splice site probability and splice site usage differences at the donor and acceptor of the target exon are used to score variants. To score variants with the splice junction head, we compute the predicted junction score differences for all junctions using the donor or acceptor of the target exon, take the max absolute difference for donor and acceptor separately and then take the sum. The aggregate splicing score sums the scores from splice sites, SSU, and splice junctions. Scores across tissues are averaged.

#### Expression Quantitative Trait Loci (eQTL) Variants

To evaluate AlphaGenome’s ability to predict the impact of variants on gene expression regulation, we built upon eQTL benchmarks established by Borzoi^3^.

**1. eQTL Dataset Preparation** For SNP eQTL evaluations we use a dataset of eQTLs from GTEx v8^23^, fine-mapped using the SuSiE method^33^. Variants with a posterior inclusion probability (PIP) *≥* 0.9 are labeled as causal, while those with PIP < 0.01 are deemed noncausal. For all reported metrics and counts of eQTLs we used variants only in the ‘test’ set chromosomes (see ‘Chromosome splits for variant benchmarks’ above), which correspond to approximately one third of the fine-mapped variants.

For Indel eQTL evaluations we downloaded the reprocessed and SuSiE fine-mapped GTEx data provided via the EMBL-EBI eQTL catalogue^92^. We filtered this data down to non-SNV variants with PIP

*≥* 0.9. We excluded variant-gene pairs where the eGene was not annotated in the GTF used throughout our analyses (Gencode V46). The resulting set contains 2645 indel eQTL (variant/gene/tissue triplets), 1535 of which are deletions.

**1. 2. Variant Effect Scoring for eQTLs** For all zero-shot eQTL evaluations we used the gene-specific RNA-seq scorer, using the ALT and REF alleles as specified for each eQTL in a given evaluation dataset, regardless of whether this REF allele corresponds to the reference genome used. This scorer summarizes the impact of the ALT allele compared to the REF allele by aggregating log fold change in predicted RNA-seq counts over a gene’s exons (Fig. 4a; Methods). We use the score for the GTEx tissue and gene corresponding to each eQTL in the evaluation dataset. For simplicity, we applied the same scoring strategy to Borzoi’s RNA-seq predictions (we note that this yielded very similar results to the author’s published best scoring method for eQTLs). For GTEx tissues unavailable in Borzoi outputs, we averaged the scores corresponding to other tissues belonging to the same broad category, as defined by Borzoi^3^ (e.g. for ‘Brain (Hippocampus)’ we used the average of scores from other brain tissues available in Borzoi outputs). For Enformer, a linear layer was fitted to predict the GTEx expression counts at TSSs from the output predicting on Enformer’s training intervals. Scores were calculated by aggregating the log fold change in predicted RNA-seq counts over a gene’s annotated transcription start site, following the variant scoring strategy as previously published (with the primary adaptation being that input sequences consistently centered the variant, in line with AlphaGenome’s standard variant input preparation).
**2. Prediction of eQTL Effect Size (Coefficient)** To assess the model’s ability to predict the magnitude of the effect of known causal variants, we evaluate its performance at predicting an eQTL’s effect size (SuSiE ‘beta posterior’). This value need not be in the same scale as the predicted effect size, but we expect the ranks and sign to be consistent. Thus, for each GTEx tissue, we compute the Spearman correlation between the predicted variant scores and observed effect size across all variant/gene pairs. We report this Spearman 𝜌 value averaged across all tissues, weighted by the number of eQTLs in each tissue. For this evaluation we used causal eQTLs as described above, consisting of n=17,675 unique variant/gene/tissue combinations, which comprises 6,626 unique variant/gene pairs, and 6,208 unique variants.
**3. Prediction of eQTL Effect Direction (Sign)** We evaluate the ability of the model to predict whether a causal variant will have a positive or negative effect on gene expression in their causal gene and tissue. An eQTL is designated as “positive” if its beta posterior is *≥* 0 (n=9,759) and “negative” if beta posterior is < 0 (n=7,916). For each tissue, we compute the auROC for the predicted values against the binary sign labels, and report the average auROC across tissues, weighted by the number of eQTLs in each tissue.
**4. Zero-Shot eQTL Causality Prediction** We evaluated the model’s zero-shot ability to differentiate causal from non-causal eQTLs using the same eQTL causality dataset as Borzoi^3^ (described above), with one adjustment to account for a distance-to-TSS bias in the original dataset (Extended Data Fig. 5b). Specifically, we computed the log of the absolute distance (bp) between the eQTL variant and the TSS of its target gene (using the median TSS across all transcripts of that gene in the GENCODE V46 GTF). We then divided the range of distances into 10 equal-width bins, and within each bin, randomly downsampled

the variant/gene/tissue tuples belonging to the non-causal class to match the number in the causal class. This resulted in 21,342 variant/gene/tissue tuples in the test set, with a class balance of exactly 50%. We measured model performance at distinguishing causal from non-causal eQTLs using the auROC metric, averaged across GTex tissues weighted by the fraction of eQTLs in each tissue. We also evaluated distance-to-TSS as a predictor using the same metric, and confirmed that it was no longer predictive in the distance-balanced causality dataset.

**1. 6. Supervised eQTL Causality Prediction** In order to leverage AlphaGenome’s multimodal predictions (not just RNA-seq) to differentiate causal from non-causal eQTLs, we trained a random forest model using the absolute values of the variant scores from multiple modalities as features (‘Causality (RF)’). We used the same dataset and metric as for zero-shot eQTL causality evaluation (see above), but used a 5-fold cross-validation approach to train and evaluate the random forest. Specifically, we trained a random forest model for each fold only using eQTLs within chromosomes *not* in the test set. We then evaluated the supervised model by making predictions using these trained models for the eQTLs in the held-out test set, and calculated the mean auROC across the 5 folds.

We used the following sets of features, each trained and evaluated separately. For a full list, see the results reported in Extended Data Fig. 5e:

- For AlphaGenome, the features consisted of a comprehensive set of variant scores derived from all its output modalities, using the scorer configurations described in the “Variant Scoring” section. For gene-specific variant scorers we used the score corresponding to an eQTL’s target gene.
- For Borzoi, features were generated using its published gene-agnostic variant scoring strategy^3^,

i.e. the L2 norm of log1p-transformed differences computed over a 524,288 bp window across all track outputs. For RNA-seq we used the same scorer as for AlphaGenome.

The Random Forest classifier was implemented using Scikit-learn’s RandomForestClassifier (version 1.0.2)^93^, with a max_depth of 5 and default values for other hyperparameters.

**1. 7. Stratification of eQTLs by Functional Annotation** To investigate the influence of functional context on model performance (Extended Data Fig. 5c), we categorized variants based on their overlap with known functional annotations, including:
Locations of 15 ChromHMM-derived chromatin states (like enhancers and repressed polycomb regions) from ROADMAP Epigenomics^23,94^.
Locations of candidate *cis* regulatory elements (enhancers, promoters, and CTCF-bound regions) from ENCODE^63,95^.
Locations of enhancers in 5 cell types from the activity-by-contact (ABC) method^96^ and enhancers and promoters (tissue-aggregated) from the updated ABC paper^97^.
Variant effect prediction annotation from Ensembl’s VEP^98^ for variants with a minor allele frequency (MAF) *≥* 0.01.
Gene-level features on expression levels and tissue-specificity of gene expression derived from GTEx.
We also derive GTF-based features such as whether the variant is in an intron, whether the target gene is protein-coding or a lncRNA etc. from a GTF.

#### Measuring Sign Prediction Coverage of GWAS

To evaluate AlphaGenome’s ability to predict the direction of effect for variants underlying GWAS signals, we developed a pipeline involving curation of GWAS credible sets, derivation of predicted effect signs, calibration against eQTLs, and stratified analysis.

**1. Curation of GWAS Credible Sets** We accessed GWAS data from Open Targets (data release 22_09, sourced from bigquery-public-data.open_targets_genetics). Starting with the Open Targets *variant_disease_credset* table, we first retained only GWAS studies. Subsequently, we selected credible sets exhibiting robust association signals, defined by having at least one variant with genome-

wide significant marginal and conditional p-values (marginal 𝑃 < 5 *×* 10*^−^*^8^ and conditional 𝑃 < 5 *×* 10*^−^*^8^), and where the sum of Posterior Inclusion Probabilities (PIPs) for all variants within the set was 1.

To ensure the independence of these signals, we resolved instances of colocalizing signals (identified through Open Targets’ colocalization analysis across studies) by selecting the credible set from the GWAS study that reported the largest number of associated loci genome-wide, using the num_assoc_loci field in the Open Targets *studies* table. If credible sets still shared the same lead variant after this step, we prioritized the study where the shared lead variant had the highest PIP, a step that excluded approximately 1,400 credible sets. Finally, we excluded credible sets from any GWAS study (n=73) that were not available in Open Targets’ *locus2gene* table. This curation process yielded 18,537 distinct credible sets from 2,476 unique GWAS studies, encompassing 581,248 unique variants.

1. **2. Deriving a Predicted Effect Sign for GWAS Loci** To hypothesize the direction of effect for the causal variant underlying a GWAS signal (i.e., a credible set) on its putative causal gene, we utilized AlphaGenome’s RNA-seq variant scores. These scores provide predictions for all proximal genes across numerous GTEx tissues, which is advantageous for subsequent calibration using eQTLs. The putative causal gene for each credible set was assigned using Open Targets’ *locus2gene* table, selecting the gene with the highest y_proba_full_model score.

We first ensure that all variants in the credible set are aligned with respect to sign. That is, the GWAS- signed AlphaGenome score for a variant and gene, 𝑆^^^_𝑣𝑔_, is the AlphaGenome score (𝑆_𝑣𝑔_) flipped such that it is aligned with respect to the risk allele (or trait-increasing allele) for a given binary (quantitative) phenotype, p. For example, if the alternative allele is associated with *decreased* risk of diabetes, we flip the sign of the score. This ensures that variants within the same credible set are always oriented such that their sign is with respect to the risk (or trait-increasing) allele.

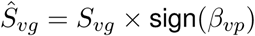

Since typically the causal tissue(s) is unknown for a GWAS, we define the AlphaGenome score 𝑆^^^_𝑣𝑔_ as the one with the largest absolute value across tracks corresponding to GTEx tissues.

Given that the true causal variant within a GWAS credible set is also usually unknown, we applied several strategies to derive a single, representative AlphaGenome score (SCG) per credible set (C) and its putative causal gene (G):

- *PIP-weighted average:* A credible set provides a discrete probability distribution over multiple candidate variants, assuming it contains a single causal variant. We can estimate the expected value of the score for the causal variant (and a given gene), by integrating over the uncertainty about which variant is causal using the law of total expectation:

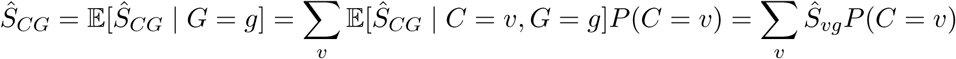

We can estimate 𝑃 (𝐶 = 𝑣) by using the PIP for each variant in a 99% credible set, normalized to sum to one.

- *PIP-max:* We use the AlphaGenome score corresponding to the variant with the largest PIP in a credible set, thus assuming that the largest PIP is the causal variant.
- *Any-variant max effect:* We use the variant with the largest absolute AlphaGenome score in the credible set. This approach assumes that variants with large AlphaGenome scores are more likely to be the causal variant for the GWAS trait.

This analysis relies on the following simplifying assumptions:

- The putative causal gene identified by Open Targets’ predictive model is correct.
- The (unknown) GWAS causal variant exerts its phenotypic effect primarily through changes in the expression of this putative causal gene, detectable within one of the GTEx tissues or cell types for which AlphaGenome provides predictions.
- **3. Calibrating to a specific sign accuracy** Many credible sets may have an AlphaGenome score that is non-zero. However, some of these may be close to zero, and not be predicting the sign reliably. We can use known expression-modulating variants (eQTLs) to pick a score threshold, 𝑇_𝑘_, that yields a particular

accuracy (e.g. 𝑘 = 80%) for predicting the correct sign. We can then ask how many GWAS credible sets

(and their corresponding putative causal genes) have an absolute score greater than a threshold, 𝑇_𝑘_.

To determine 𝑇_𝑘_ we used the same set of fine-mapped eQTLs as in the ‘eQTL Sign’ evaluation, where we have AlphaGenome scores and strong evidence (PIP>0.9) that the variants are causal (for gene expression) in a given gene, as well as their observed direction of effect, with one change to account for the uncertainty in the causal tissue in the GWAS setting. That is, for a given variant/gene pair in the evaluation dataset, we define the ground truth sign as the sign of the tissue with the largest maximum absolute observed effect size, but we use the sign of the tissue with the largest maximum AlphaGenome score across *all* GTEx tissues in order to calculate accuracy. In Fig. 4h the calibrated score thresholds used for 80% (90%) accuracy are 𝑇_80_ = 0.015 (𝑇_90_ = 0.107).

1. **4. Identification of eQTL-Colocalizing GWAS Loci** We identified GWAS credible sets likely sharing a causal variant with an eQTL by utilizing Open Targets’ colocalization data (*variant_disease_coloc* table). Credible sets demonstrating a COLOC^35^ posterior probability of a shared causal variant (H4) greater than 0.95 were classified as colocalizing. Out of the 18,537 curated GWAS credible sets, 3,132 (17%) satisfied this colocalization criterion.
2. **5. Stratification of GWAS Loci for Analysis** To explore how sign prediction coverage by AlphaGenome and eQTL colocalization varied with the characteristics of GWAS signals, credible sets were stratified based on the following criteria:
3. *Single-variant credible sets:* These are sets that contain only one variant, thereby having little to no ambiguity regarding the identity of the potential causal variant.
4. *Minor Allele Frequency (MAF):* Credible sets were categorized by the PIP-weighted average MAF of their constituent variants. Sets falling into the lowest quintile of these weighted average MAFs were labeled ‘small MAF,’ and those in the highest quintile were labeled ‘large MAF.’ MAF values for Non-Finnish Europeans were used, as supplied in the Open Targets’ *variants* table (originally sourced from gnomAD).
5. *GWAS effect size*: A similar quintile-based categorization was applied using the PIP-weighted average of the absolute value of each variant’s estimated GWAS effect size (beta coefficient, from the *tag_beta* column in the Open Targets credible set table), defining ‘small effect size’ and ‘large effect size’ subsets.
6. *High Causal Gene Probability:* Credible sets where the putative causal gene – as assigned by

Open Targets’ predictive causal gene model (detailed in their *locus2gene* table) – had an estimated probability (y_proba_full_model column) greater than 0.8.

#### Polyadenylation variants

We evaluate the model’s ability to predict the effects of polyadenylation-altering variants using the tissue-pooled paQTL variant set from Borzoi, which contains fine-mapped 3’ QTLs from the eQTL Catalog^87^. For all paQTL metrics, we used variants only in the ‘test’ set chromosomes (see ‘Chromosome splits for variant benchmarks’ above). We follow Borzoi paQTL processing steps to generate a dataset of causal paQTLs (n=613) that affect mRNA isoform abundance by altering polyadenylation (pA) site usage, together with a negative set (n=1950) controlled for distance to the nearest 3’ cleavage site and for similar expression levels. We applied Borzoi’s approach to calculating AUPRC by performing 100 permutations of randomly matching each positive SNP to one of multiple potential negative SNP matches and calculating the AUPRC. We then average AUPRC performance across all permutations to get our final metric. SNPs that are not scored because they do not have > 80% of PAS within the input interval or only have one PAS site are imputed as 0 scores for metric calculations.

Following Borzoi’s methodology, we score paQTLs by comparing predicted RNA-seq coverage between the reference and alternative alleles at 3’ cleavage sites i.e. polyadenylation signals (PASs). The scoring approach quantifies the log fold change in expression between the set of proximal PASs vs. the set of distal PASs, where the sets of proximal and distal PASs are the PASs upstream or downstream a given 3’ cleavage site respectively. For each gene within the input window, and for both REF and ALT predictions:

1. Each PASs is extended with 400 bp of upstream context.
2. Predicted RNA-seq coverage is summed spatially to produce a single aggregated value per PASs per RNA-seq track.
3. For each possible split of PASs into proximal and distal sets, we compute the absolute log fold change: *|* log(ALT/REF)*|*.
4. The maximum value over all possible splits is taken for each track, producing one score per track.
5. The average score across tracks is then used as the final score for metric calculations.

#### Chromatin accessibility variants & bQTLs

We sourced the following chromatin accessibility variants - DNase-seq quantitative trait loci (dsQTLs) and chromatin accessibility QTL (caQTL) - evaluations from ChromBPNet paper.

We evaluate the models on their ability to distinguish causal versus non-causal variants, and the effect size of causal variants, on the following datasets:

- *dsQTL Yoruba*. Yoruba African ancestry dsQTLs from 70 lymphoblastoid cell-lines (LCLs).
- *caQTL European.* European ancestry caQTLs from ATAC-seq profiling in ∼100 LCLs.
- *caQTL African.* Representing African ancestry of six ancestry subgroups caQTLs from ATAC-seq profiling in ∼100 LCLs.

As per ChromBPNet, we further evaluate the models on their ability to predict the reported effect size of significant caQTLs in 2 specific cell types *microglia* and *coronary smooth muscle cells (SMCs)* by indexing into closest matching tracks. We also evaluate the model on their ability to predict binding QTL (bQTL) variants for the SPI1 pioneer TF identified from 60 Yoruban LCLs by indexing into SPI1 ChIP-seq tracks for Borzoi and AlphaGenome.

Although ChromBPNet trained specialized cell type-specific models for these tissues, we evaluate AlphaGenome and Borzoi zero-shot on these tasks by indexing into the closest matching tracks. As with all variant evals, we assign a VALID/TEST chromosome split and only report the TEST numbers.

- *Cell type matching for GM12878.* For the Yoruba dsQTL, European caQTL, and African caQTL tasks (causality and effect size prediction), these variants are from LCLs. The ChromBPNet model

was trained on a single reference LCL (GM12878). We zero-shot extract AlphaGenome GM12878 predictions. AlphaGenome contains GM12878 predictions for both ATAC and DNase outputs, and we find that DNase GM12878 predictions perform slightly better. For Borzoi, GM12878 was only available for DNase, there are 2 tracks and we take the mean of these. We use the same recommended scorer for both (center mask, width=501, aggregation DIFF_LOG2_SUM which is LFC).

- *Cell type matching for microglia*. On the validation set, we examined which AlphaGenome tracks performed best for predicting microglia caQTL effect sizes and found that it was Microglia ATAC for Borzoi (kai182) and suppressor macrophage DNase (CL:0000862) for AlphaGenome (microglia are a type of macrophage). We then used these columns for predictions on the test set.
- *Cell type matching for coronary smooth muscle cells*. Similarly, we found that AlphaGenome’s “Left cardiac atrium” (UBERON:0002079) ATAC track and Borzoi’s “Vascular Smooth Muscle 2” (kai222) ATAC track performed best on the validation set.
- *Cell type matching for SPI1 bQTL tasks.* For AlphaGenome, we use ChIP-seq predictions for the SPI1 TF in GM12878 (EFO:0002784). Borzoi did not have SPI1 GM12878 ChIP-seq, the closest match was SPI1 GM12891.

#### Prioritization of Trait-Associated GWAS Variants

To assess AlphaGenome’s ability to prioritize functional variants from GWAS, we first curated a benchmark dataset using credible set data from Open Targets^99^ (sourced from bigquery-public-data.open_ targets_genetics, release 22_09). This initial dataset contains variants statistically associated with a wide range of diseases and complex traits such as height and BMI. We applied stringent filtering to retain only high-confidence credible sets, requiring that at least one variant within each set demonstrated genome-wide significance for both marginal and conditional associations (marginal ‘tag_pval’ 𝑃 < 5 × 10*^−^*^8^ and conditional ‘tag_pval_cond’ 𝑃 < 5 *×* 10*^−^*^8^), and that the sum of Posterior Inclusion Probabilities (PIPs) for variants within any given set did not exceed 1.

To focus on potentially non-coding regulatory variants, we then excluded variants whose most severe consequence, as annotated by the Ensembl Variant Effect Predictor (VEP)^98^, fell into protein-altering categories. The excluded categories comprised: ‘missense_variant’, ‘frameshift_variant’, ‘stop_gained’, ‘inframe_deletion’, ‘inframe_insertion’, ‘start_lost’, ‘stop_lost’, ‘coding_sequence_variant’, and ‘pro- tein_altering_variant’. Following this, variants were deduplicated by selecting their maximum PIP value across all associated phenotypes. For the benchmark, variants with a final PIP *≥* 0.9 were labeled as ‘causal’ (positive examples), while those with PIP < 0.01 were labeled as ‘non-causal’ (negative examples). This dataset was subsequently rebalanced at the variant ID level to achieve an equal 50/50 ratio of causal to non-causal variants.

Given that GWAS variants can influence phenotypes through diverse molecular mechanisms affecting multiple genomic features, we trained a Random Forest (RF) classifier to integrate information from model-derived variant scores across modalities. This allowed us to evaluate the collective power of these scores in distinguishing causal from non-causal GWAS variants. The RF models were trained and evaluated using a 5-fold cross-validation scheme on the curated GWAS variant dataset, with average auROC performance reported. Input features were as follows:

- For AlphaGenome, the features input to the classifier consisted of a comprehensive set of variant scores derived from all its output modalities, using scorer configurations previously optimized on modality-specific evaluations (see Variant Scoring section).
- For Borzoi, features were generated using its published gene-agnostic variant scoring strategy^3^, which involves calculating the L2 norm of log1p-transformed differences over a 524,288 bp window across all its track outputs.

In all cases we used the absolute values of scores for this task. The Random Forest classifier was implemented using the RandomForestClassifier function from Scikit-learn (version 1.0.2)^93^, with the max_depth parameter set to 5 and other hyperparameters kept at their default values without further optimization. Note that this benchmark does not use the chromosome split described previously, as we did not use it to optimize our variant scorers.

The features used for the ‘baseline’ analysis were derived from publicly-available sources. We used the same set of features as listed in *eQTL functional segmentation*, except for those related to a specific gene (last to bullets).

#### TraitGym

To train and evaluate models using a random forest on the Traitgym complex variant benchmark, we followed the same approach as above.

To evaluate the ability of sequence-to-function models to predict trait-affecting variants (mendelian and complex) in a zero-shot fashion, we followed Benegas et al^47^. Specifically, for each track predicted by a model, we first computed the predicted log-fold change in activity per position (or bin) due to the variant and then calculated the L2 norm across the sequence. For simplicity, we excluded tracks which were not easily amenable to this score, either due to their sparsity (splice-site location and usage) or because they involved an additional dimension (contact maps and splice junctions). As a result, Borzoi and AlphaGenome have access to the same modalities in this evaluation. As in Benegas et al., we averaged effect predictions from the forward and reverse strand sequence for all models (see ‘Test-time augmentation’).

To aggregate variant scores across modalities, Benegas et al. applied another L2 norm along the track dimension. However, the models we analyzed differ in the number of tracks they predict, how these tracks are scaled and at which resolution they are predicted. Therefore, a single aggregation strategy may not perform well for all tested models. To account for this possibility, we evaluated each model under three different aggregations (L2, mean and max) and took the respective best. We found that AlphaGenome always performed best when taking the max across tracks. For mendelian traits, the max across tracks also performed best for Enformer and the Borzoi ensemble, but not for Borzoi *fold-0*, which exhibited a slight boost from using the L2. For complex traits, both Borzoi and Enformer performed best under the L2 norm. Note that this benchmark does not use the chromosome split described previously, as we did not use it to optimize our variant scorers.

#### Enhancer-gene linking

Our evaluations are based on the dataset of CRISPRi-validated enhancer-gene pairs assembled in the ENCODE-rE2G study^17^. We followed the authors’ approach of filtering for genes present in their annotations, resulting in 471 positives and 10356 pairs in total. We needed to also drop 3 additional element-gene pairs whose gene ids were not present in our annotation file (GENCODE v46), resulting in a final dataset of 471 positives and 10353 examples in total.

**Zero-shot evaluation** Each element-gene pair was scored using an established input gradient scoring method, derived from K562 RNA-seq data as previously described^3^, to directly assess sequence importance. That is, we centered the reference input sequence on the target gene midpoint and computed gradients with respect to the input. The scalar used for gradient computation was the average predicted gene expression (log sum across exons) across K562 tracks. We then computed each element-gene pair score by taking a weighted average of absolute input gradient contribution scores in the local window centered at the element. A Gaussian kernel was used to compute the weighted average, with window size 2400 and standard deviation 300. Element-gene pair scores were normalized by dividing by the mean absolute input gradient across the input sequence, to account for genes being expressed at different levels. Putative enhancers falling outside a model’s input context window (i.e. too far from the target gene) were imputed with zero scores.

The above strategy was applied to AlphaGenome and Borzoi-ensemble models. We benchmarked the models by comparing the area under the Precision-Recall curve for selected TSS distance bins, borrowing the ones reported in the ENCODE-rE2G paper. We used the TSS annotations provided by ENCODE-rE2G for binning the elements. As simple baselines, we have included random predictions and the inverse distance to TSS.

Complementing this, we also scored enhancer-gene pairs using our variant scorers by permuting the 2kb interval centered on the enhancer element 10 times and taking the mean effect on a specified track as the score (Extended Data Fig. 7a).

**Supervised evaluation** The ENCODE-rE2G models^17^ are logistic regression models trained on bundles of features, and evaluated on out-of-fold predictions. For ENCODE-rE2G and ENCODE-rE2G extended, we used the precomputed cross-validated scores on the K562 cell line provided by the authors. We evaluated the effect of including AlphaGenome-based features in their feature bundles by including the K562 RNA-seq input x gradient score from AlphaGenome as a single additional feature into the full ENCODE-rE2G extended feature set (Extended Data Fig. 7b) and re-running their full training pipeline (forked from their repository). We also evaluated the performance of this input x gradient feature alone, and in combination with the TSS distance feature.

Subsequently, we used a more comprehensive set of AlphaGenome variant scorer-based features from K562 including Allele-Specific Activity Scores (AAS) and differential variant effect scores for RNA-seq of the target gene, ChIP-seq for EP300 and H3K27ac, CAGE, Pro-cap, as well as H1-ESC contact maps (K562 contact maps were not in our model outputs). These variant scores were combined with the existing ENCODE features for training the supervised regression model (Extended Data Fig. 7c).

#### Benchmarking on the CAGI5 MPRA challenge

AlphaGenome’s performance variant effect prediction was further evaluated on the experimental massively parallel reporter assays (MPRA) data from the CAGI5 challenge. All predictions were made using the native genomic sequence surrounding each variant, using hg19 (human) reference genome and GTF V19 annotations to extract input sequences to match vcf file coordinates. Variant effect scores were computed following either using our recommended variant scorers or the Enformer or Borzoi strategies as described below. Zero-shot performance was computed by taking the Pearson *r* correlation between DNASE variant effect scores averaged across cell type-matched DNase tracks and observed CAGI5 effects. Following our previous work, for the lasso regression we scaled test set features using scaling factors from the training set such that the training features had a mean and standard deviation of 0 and 1 respectively. We then applied sklearn.LassoCV to train a model for each locus using the corresponding CAGI5 challenge training set with 10-fold cross validation and n_alphas = 100. Enformer Lasso comparisons were cell type-agnostic, using the full set of cell type tracks for CAGE and DNase. Borzoi comparisons used either DNase alone, or jointly with ChIP-Histone and RNA output types, in combination with either a cell type-agnostic or cell type-matched strategies described below.

For comparisons against Enformer, we used the previously defined variant scoring strategy of using the predicted difference in coverage between the reference and alternative allele summed across a 512 bp mask centered on the variant (CAGE and DNase modalities). All CAGE features had a 1 pseudocount added and then were log transformed before computing this difference. For cell type-matched comparisons, we used the same gene name to substring mapping as defined by the paper with the following modifications for when there was no available match. (1) The additional inclusion of ‘melanocyte’ for *IRF4*, (2) The addition of ‘adrenal gland’ for *TERT* performed in GBM cells.

For Borzoi comparisons, we followed their strategy of using the log-fold changes of total coverage summed across a 4000 bp window centered on the variant (DNase, CHIP-HISTONE) or from the log fold changes of summed exon coverage of the target gene (RNA-SEQ). For cell type-matched comparisons, we used the same gene name to substring mapping as defined in their report with the following modifications. (1) The additional inclusion of ‘melanocyte’ for IRF4, (2) the substitution of the substring ‘adrenal gland’ for the kidney tubule cell ontology CURIE ‘CL:1000507’ for the loci *LDLR, HNF4A, MSMB, TERT, MYC* for subsetting DNASE and CHIP-HISTONE predictions. RNA predictions were still subsetted using ‘adrenal gland’ for these loci. See Supplementary Table 10 for complete list of ontology CURIEs extracted from AlphaGenome predictions.

For average comparisons described in Figure 5j, the following loci are excluded to ensure a fair comparison: TERT-GBM (TERT variants tested in GBM cells) because ChromBPNet only reports TERT-HEK293T variant performance, and MYC due to non-convergence of LASSO regression for both AlphaGenome and Borzoi. F9 is excluded from comparisons of LASSO regressions using only DNASE features due to non-convergence when using cell type-matched DNase features for both AlphaGenome and Borzoi. The F9 Pearson r performances using cell type-agnostic DNase features for AlphaGenome and Borzoi were 0.62 and 0.54 respectively. When cell type-matched comparisons are specified, AlphaGenome used the modified Borzoi cell type-matching strategy as described above.

#### Multimodal variant example: TAL1

We gathered clinically observed variants from published literature reporting T-ALL noncoding variants around *TAL1* (Supplementary Table 11): a cluster of 5’ neo-enhancer mutations upstream of the *TAL1* TSS^42,43^; an intronic single nucleotide variant^44^; and a 3’ neo-enhancer^45^.

To predict the impact of mutations on TAL1 gene expression and local epigenetic changes, we used AlphaGenome’s variant scorers. Variant effects of cancer-associated mutations were compared against background variants (length-matched randomly shuffled insertions at the loci). Background variants were either saturating (all possible combinations at that length), otherwise 100 shuffles were used. Variant scores in CD34+ common myeloid progenitors from TAL1 RNA-seq, DNAase, histone modifications such as H3K4me1/3, H3K27ac, H3K27me3, H3K9me3, H3K36me3) were collected. There is no tissue matched contact map, therefore we reported the mean effect across all contact map tracks.

To visualize a heatmap of these predicted effects, variant effects for each track were min-max scaled, grouped by their insertion length and position (as grouped in Fig. 6c).

#### Model Ablations

The following sections detail the specific experimental setups for the ablation studies presented in Fig. 7. Unless otherwise specified, models were trained using the same architecture, pre-training data, optimization procedures and evaluation metrics as described throughout the Methods section. All ablation experiments were conducted using four independent training runs with different random seeds, unless stated otherwise.

**Impact of Target Resolution** To evaluate the impact of prediction resolution, models were trained to predict targets for DNA accessibility (ATAC-seq, DNase-seq), gene expression (RNA-Seq, CAGE-seq, PRO-cap), and splicing (splice site classification, splice site usage, splice junction counts) at varying output resolutions (1 bp, 2 bp, 8 bp, 32 bp, and 128 bp).

This required different treatment depending on the specific output:

- For track modalities (RNA-Seq, ATAC-seq, DNase-seq, CAGE-seq), lower-resolution targets were generated by summing the 1 bp resolution ground truth counts across consecutive bins corresponding to the target resolution after augmentation was applied at base resolution.
- For splice site classification and usage, which are per-base predictions, the 1 bp resolution ground truth labels (positive classes) were max-pooled to the target resolution.
- For splice junction count prediction, which relies on base-resolution positions of donor and acceptor sites, lower resolution embeddings from the Output Embedder were upsampled by repeating along the sequence axis (e.g., 4 times for 4 bp embedding), and used by the junction prediction head as if at base-resolution.

For these ablations, the core model architecture, including the decoder structure up to the point of embedding extraction, remained unchanged; only the extraction point of the final output embedder and the resolution of the target data and corresponding loss calculations were modified.

**Impact of Sequence Length during Training and Inference** This ablation explored the effect of varying input DNA sequence length (8kb, 32kb, 131kb, 512kb, and 1 Mb) on model performance. AlphaGenome’s architecture is designed to be trained or evaluated at any sequence length that is a multiple of 2048 (the resolution of the pair activations modules), and the training and evaluation sequence lengths can differ. Three scenarios were tested: Three scenarios were tested:

- *Fixed Training Length, Variable Evaluation Length (Blue Series):* A single set of models, pre-trained using 1 Mb input sequences, was evaluated using varying input sequence lengths at inference time.
- *Variable Training Length, Fixed Evaluation Length (Purple Series):* Models were trained using input sequences of varying lengths. During training, the batch size was adjusted inversely to the input sequence length to maintain a constant cumulative sequence length processed per gradient step across runs. All models in this series were subsequently evaluated using a fixed 1 Mb input sequence length during inference.
- *Matched Training and Evaluation Length (Green Series):* Models were trained and evaluated using the same matched input sequence length.

**Impact of Ensembling and Distillation** To assess the benefits of ensembling and distillation, 64 independent models were pre-trained using the Fold 0 data partition. The performance of mean ensembles was evaluated by averaging the predictions of randomly selected subsets of these 64 pre-trained models, with ensemble sizes ranging from 1 to 4. Single student models (with the same architecture) were produced by distilling knowledge from ensembles of these 64 Fold 0 pre-trained teacher models, using 1, 4, or 64 unique teachers in the distillation process, following the procedure described in the main “Distillation” methods.

**Impact of Multimodal Learning (Modality Ablations)** To investigate the contribution of different data types to learning shared representations and to overall predictive performance, models were trained with gradients restricted to specific modality groups. For each ablated model, the standard loss functions were applied normally to all prediction heads. However, for prediction heads outside this target group, stop-gradients were applied between the model embeddings and the head linear layer for both mouse and human tracks within each modality group. This ensured that these other heads were still trained (allowing their accuracy to be evaluated) but did not contribute to the learning of the shared model trunk representations.

The performance of these modality-specific models (n=8 seeds per group) was compared against the fully trained multimodal model (n=4 seeds). The modality groups investigated were: accessibility (ATAC-seq, DNase-seq, and chromatin contact maps), expression (RNA-Seq, CAGE-seq, and PRO-cap), splicing (splice site classification, splice site usage, and splice junction counts), and Histone ChIP-seq.

**Evaluation Metrics for Ablation Studies** The performance across these ablation studies was assessed using a consistent set of metrics. These include: RNA-Seq Pearson correlation (log1p transformed counts), RNA-Seq gene-level Pearson correlation (Log Fold Change), Junction Counts Ratio Pearson correlation, ATAC-seq Pearson correlation, Histone ChIP-seq Pearson correlation, Contact Maps Pearson correlation (computed at 2048 bp resolution and averaged over all 28 human datasets), eQTL Sign prediction auROC, eQTL Causality prediction auROC (gene-balanced dataset), sQTL Causality prediction auPRC, Splicing Outlier prediction auPRC, Chromatin Accessibility Variants Causality auPRC (average of dsQTL Yoruba, European caQTL, and African caQTL tasks), and paQTL prediction auPRC. These evaluations were done on human tracks only.

#### Model performance analysis and visualization

Data analysis used Python v.3.11.8 (https://www.python.org/), NumPy v2.2.5 (https://github.com/ numpy/numpy), SciPy v.1.14.1 (https://www.scipy.org/), seaborn v.0.12.2 (https://github.com/ mwaskom/seaborn), Matplotlib v.3.9.1 (https://github.com/matplotlib/matplotlib), Pandas v.2.2.3 (https://github.com/pandas-dev/pandas), anndata v0.11.4 (https://github.com/scverse/anndata), Scikit-learn v1.6.1, (https://github.com/scikit-learn/scikit-learn), InterpretML v0.6.10 (https://github.com/interpretml/interpret), and Colab (https://research.google.com/colaboratory).

## Acknowledgements

We thank Dhavanthi Hariharan, Charlie Taylor, and Ottavia Bertolli for product and program management support, and Molly Beck and Uchechi Okereke for legal counsel. We are grateful to Yannis Assael and Alex Botev for contributions to modeling and engineering; and to Anna Trostanetski, Lucas Tenório, Victoria Johnston, Richard Green and Sarah Chakera for their work on the API release. We also thank Kathryn Tunyasuvunakool for feedback on the manuscript, Rachael Tremlett for assistance with graphic design and figure preparation, and former interns Ibrahim I. Taskiran, Andreea-Alexandra Muşat, Raiyan Khan, and Ren Yi for their contributions to earlier stages of this research. Gemini (Google) was used for assistance with language editing and improving the clarity of the manuscript. This work extensively utilized publicly available datasets. We extend our sincere gratitude to the numerous research consortia (including, but not limited to, ENCODE, GTEx, FANTOM5, and the 4D Nucleome Project), their members, data contributors, and all study participants whose efforts in generating and openly sharing these foundational genomic resources were essential for this study.

## Author Contributions

Z.A. conceptualized the study with input from N.L., J.C., C.B., T.W., G.N., K.R.T, P.K and D.H. Z.A. led the project with help from N.L. leading manuscript preparation, G.N. leading overall model development, J.C. conceiving and leading the splicing-related research, and K.R.T., T.W leading engineering. Z.A., G.N. conceived the device-distributed model architecture with input from R.T.. G.N. conceived the contact-maps pair stack. J.C., G.N. conceived and developed splice junction prediction head with contributions from Z.A.. Z.A., G.N. conceived model distillation with input from J.C., V.D., K.R.T., G.N. developed the model with contributions from Z.A., J.C., K.R.T., T.W., E.A., V.D., S.D.. A.G., A.M., A.K., A.S.. Z.A., J.C. and G.N., curated, processed and prepared the training data with contributions from L.N., K.R.T, T.W., J.P., M.P., T.S., A.M.. T.W, T.S., developed the data pipeline with contributions from Z.A., J.C., G.N., K.R.T., L.N.. Z.A., J.C., K.R.T., T.W., L.N., E.A designed variant scoring infrastructure with contributions from N.L., C.B., R.T., T.A. G.N., K.R.T., T.W., L.N., E.A., V.D. optimised the model and variant scoring runtime. C.B. conceived and developed variant score normalization with contributions from Z.A. and T.W.. T.W. conceived and developed model serialization and inference infrastructure with contributions from Z.A., K.R.T., L.N.. L.N. developed the evaluation infrastructure with contributions from N.L., K.R.T., T.W., R.T., V.D., and A.M. Z.A., N.L., J.C., G.N., C.B., L.N., E.A., J.P., R.T., V.D., M.P., A.K., A.G., A.M., P.D. evaluated the model. Specific evaluation areas were covered as follows: Z.A. and V.D. covered track evaluation, J.C. and E.A. covered splicing track and variant evaluation; N.L. and C.B. covered eQTLs, E.A. and J.P. covered enhancer to gene linking, A.Ka. covered eQTL indel effects; N.L. covered accessibility QTLs; J.P. covered multimodal examples, N.L. and G.N. covered contact maps; C.B. covered GWAS and complex traits analysis with help from A.Ka., J.P., and T.A.; R.T. and M.P. covered polyadenylation QTLs and CAGI5; A.Ka. covered TraitGym. Z.A., N.L., J.C., G.N., K.R.T., T.W., C.B., L.N., E.A., J.P., R.T., V.D., M.P., A.Ka. analyzed data and prepared figures. Z.A., N.L., J.C., G.N., K.R.T., T.W., C.B., L.N., E.A., J.P., R.T., V.D., M.P., A.Ka., A.G., S.B. wrote the manuscript. L.H.W. managed and coordinated the project planning and execution Z.A. and P.K. managed the project. All authors reviewed the manuscript.

## Data Availability

All primary experimental datasets utilized for the training and evaluation of AlphaGenome in this study were obtained from publicly accessible sources. A comprehensive manifest detailing these data sources including specific repositories (e.g., ENCODE portal, GTEx portal, 4D Nucleome portal, ClinVar, gnomAD), individual accession numbers, relevant version information, and direct URLs where applicable is provided in Supplementary Table 2. This study did not generate new primary experimental data requiring deposition.

## Code Availability

AlphaGenome will be available for non-commercial use via an online API at http://deepmind. google.com/science/alphagenome, with an accompanying Python software development kit (SDK) provided to interact with the model. The model source code and weights will also be provided upon final publication.

We also provide a genome interpretation suite to facilitate the exploration and interpretation of AlphaGenome. This offers a range of functionalities, for example streamlined variant scoring with quantile calibration, and identification of critical sequence regions via contribution scores from ISM-based experiments. Both the model and interpretation SDK are available at https://github.com/google-deepmind/alphagenome.

## Supplementary Information

## Supplementary Note: AlphaGenome shows competitive performance on the CAGI5 MPRA bench- mark

Massively parallel reporter assays (MPRAs) provide high-throughput functional readouts, experimentally measuring the impact of sequence variation on gene regulation, albeit in artificial constructs lacking native chromatin context. Building upon previous work demonstrating strong performance by models like Enformer on the CAGI5 saturation mutagenesis MPRA challenge^25,26^, we evaluated AlphaGenome on the same dataset, comparing against Enformer, Borzoi, and ChromBPNet (Extended Data Fig. 8). Using a zero-shot approach with cell-type matched scores derived from accessibility predictions alone (DNase), AlphaGenome performed comparably to both ChromBPNet and the Borzoi Ensemble (overall Pearson 𝑟 *≈* 0.54-0.56; Extended Data Fig. 8A). Furthermore, employing a cell-type agnostic strategy and using LASSO regression to aggregate features derived from multiple AlphaGenome tracks (e.g., DNase combined with CAGE), AlphaGenome achieved strong correlations (𝑟 *≈*0.61) comparing favorably to both Enformer (𝑟 *≈* 0.548) and Borzoi (𝑟 *≈* 0.549) across different feature sets (Extended Data Fig. 8B). We then explored the use of LASSO regression on other modalities. Using Borzoi’s variant scoring strategy of ensembling over cell-type matched tracks from multiple modalities (DNASE and CHIP and RNA or only DNASE and CHIP, Extended Data Fig. 8C) improved both AlphaGenome (𝑟 *≈* 0.609 - 0.635) and Borzoi’s (𝑟 *≈* 0.599 - 0.635) performances. Finally, we tried a cell type agnostic LASSO regression across these combined modalities using recommended scorers for each modality (Extended Data Fig. 8D) and saw AlphaGenome’s highest performance (𝑟 *≈* 0.648 - 0.662) and (Borzoi 𝑟 *≈* 0.584 - 0.601), indicating that specific cell type matching may not be necessary, and can hinder performance if they are not optimized properly. These results on an independent MPRA benchmark further validate AlphaGenome’s ability to predict the effects of regulatory variants on gene expression, achieving performance competitive with state-of-the-art models.

## Supplementary Note: Trait-altering variants

Linking genetic variants to their phenotypic consequences, particularly complex traits and diseases, remains a central challenge in genomics. This task is complicated by the difficulty in establishing gold- standard causal variants for benchmarking, often relying on fine-mapped GWAS signals or curated databases like OMIM. Such limitations frequently necessitate aggregating data across diverse traits, potentially introducing heterogeneity. Furthermore, the optimal strategy for selecting negative variants in these benchmarks is often unclear, yet it critically influences performance assessment. To evaluate AlphaGenome’s utility in prioritizing trait-associated variants, we explored two benchmarking approaches. The first was an in-house benchmark using Open Targets data, where positives were fine-mapped GWAS variants (PIP >= 0.9) and negatives were sampled from low-PIP variants (PIP < 0.01) to achieve a 50:50 class balance. The second, TraitGym, also uses fine-mapped GWAS variants as positives but employs a negative set matched for baseline features like TSS distance, MAF, and linkage disequilibrium (LD), with a 9:1 negative-to-positive class balance. A key question is how to best leverage sequence- to-function models like AlphaGenome for this task, as they do not directly predict traits or pathogenicity. Intuitively, non-coding trait-associated variants should alter regulatory outcomes detectable by such models. However, the specific gene, tissue, and regulatory modality affected are often unknown, necessitating methods to aggregate multiple model predictions. We explored both training random forests for this aggregation and a simpler zero-shot approach (e.g., taking the maximum predicted effect). Our benchmarking revealed that the choice of negative set heavily influences apparent predictor performance. When negative variants were not matched for baseline features, these baseline features themselves showed predictive power, although AlphaGenome modestly outperformed them and Borzoi (Supplementary Fig. 10a). In this scenario, modalities detecting variant location within cis-regulatory elements were more predictive than actual variant effect scores. Conversely, when negatives were matched for baseline features (as in TraitGym), these features were, unsurprisingly, not predictive (Supplementary Fig. 10b). Here, variant effect scores became more informative, though the performance differences between models using either random forest aggregation or zero-shot approaches were not substantial (Supplementary Fig. 10c). We found no evidence that explicitly adding baseline features improved AlphaGenome’s predictions, suggesting it inherently captures such information. Overall, however, robustly solving the general variant-to-trait prioritization problem remains challenging. GWAS signals are often distally located, posing difficulties for current sequence-based models, and complex traits likely require integrating information beyond cis-regulatory effects, such as gene function and network interactions. Even perfect prediction of steady-state gene expression changes due to cis- regulatory variants would likely be insufficient for many traits. TraitGym also provides a benchmark for distinguishing Mendelian disease variants (typically rare and monogenic) from TSS-distance and consequence matched common variants. On this task, where mechanisms might be simpler and more directly tied to strong regulatory feature changes, sequence-to-function models, including AlphaGenome, generally performed better than on the GWAS benchmark (Supplementary Fig. 10d). AlphaGenome offered a slight improvement over previous sequence-based models. However, broad-spectrum predictors like CADD16, which leverage conservation, remain highly competitive, partly because conservation implicitly accounts for gene essentiality and function – factors that sequence-to-function models focusing purely on regulatory impact by design will not capture. We also note that in the TraitGym Mendelian benchmark, due to matching criteria, about 7% of negative control variants had a high probability (PIP > 0.5) of being eQTLs in at least one study in the eQTL catalogue, which sequence-to-function models might understandably rank highly. Overall, it will likely be necessary to integrate sequence-to-function models with other measures, such as conservation and estimates of gene function, to enable direct predictions of variant deleteriousness.

---

**Extended Data Figure 1.**
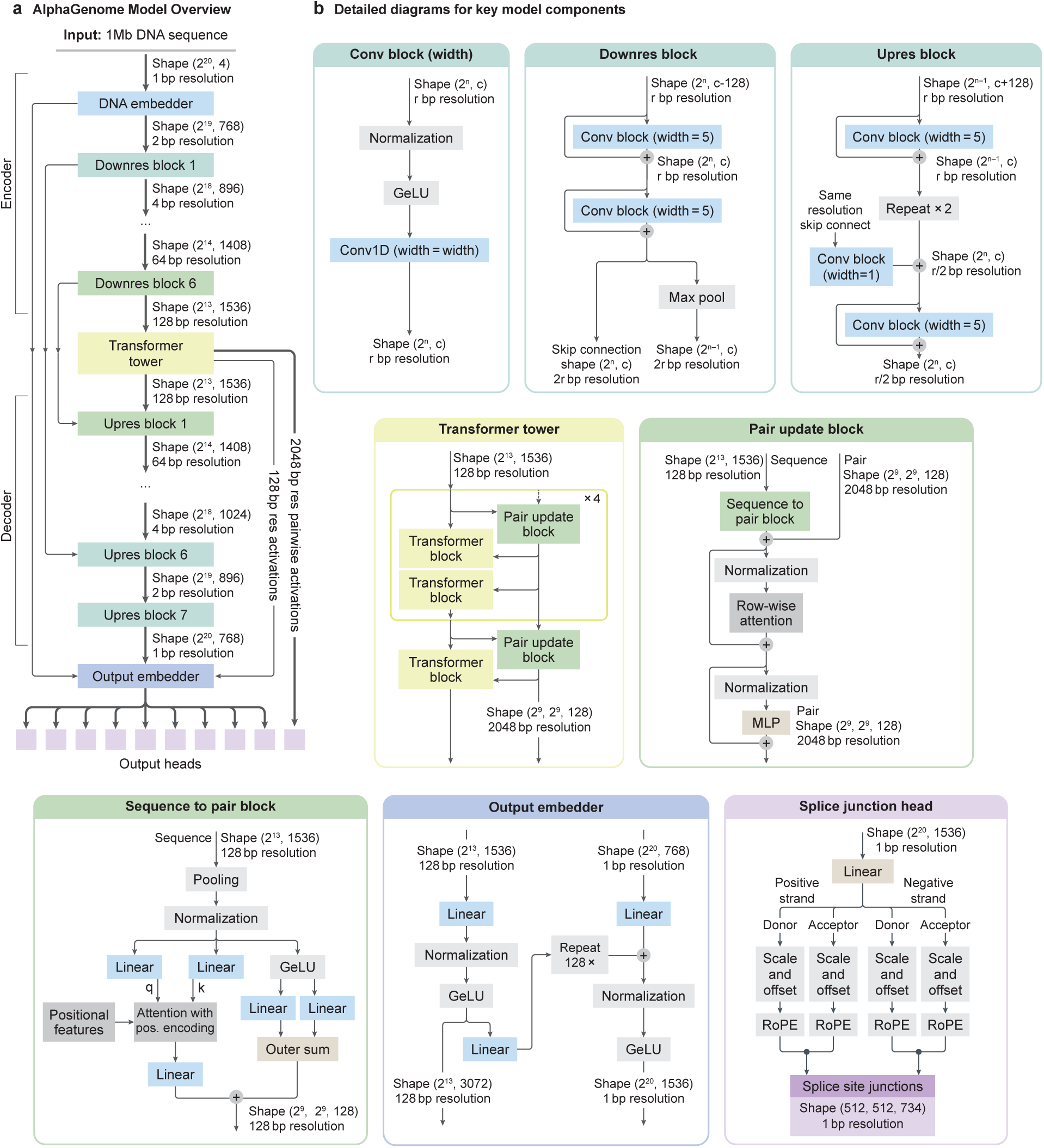
AlphaGenome model architecture. **(a)** Overview schematic illustrating the flow of activations through the model. The architecture follows a U-Net-like structure with an Encoder, a central Transformer Tower, and a Decoder processing a 1Mb DNA input sequence. The Encoder uses convolutional blocks and max pooling to progressively downsample the sequence resolution (from 1 bp to 128 bp) while increasing feature channels. The Transformer Tower operates at 128 bp resolution, iteratively refining sequence representations and generating pairwise (2D) representations. The Decoder uses convolutional blocks and upsampling, incorporating skip connections (dashed lines) from corresponding Encoder stages, to restore sequence resolution up to 1 bp. An Output Embedder performs final processing before feeding representations to task-specific output heads. **(b)** Internal structure of key component blocks used repeatedly within the architecture overview shown in (a). Diagrams detail the layers within the convolutional blocks (Conv block, Upres block), the Transformer blocks, and the blocks responsible for generating and updating pairwise representations (Pair update block, Sequence to pair block). Tensor shapes are shown excluding the batch dimension. Abbreviations: r = log-resolution, c = channels.

**Extended Data Figure 2.**
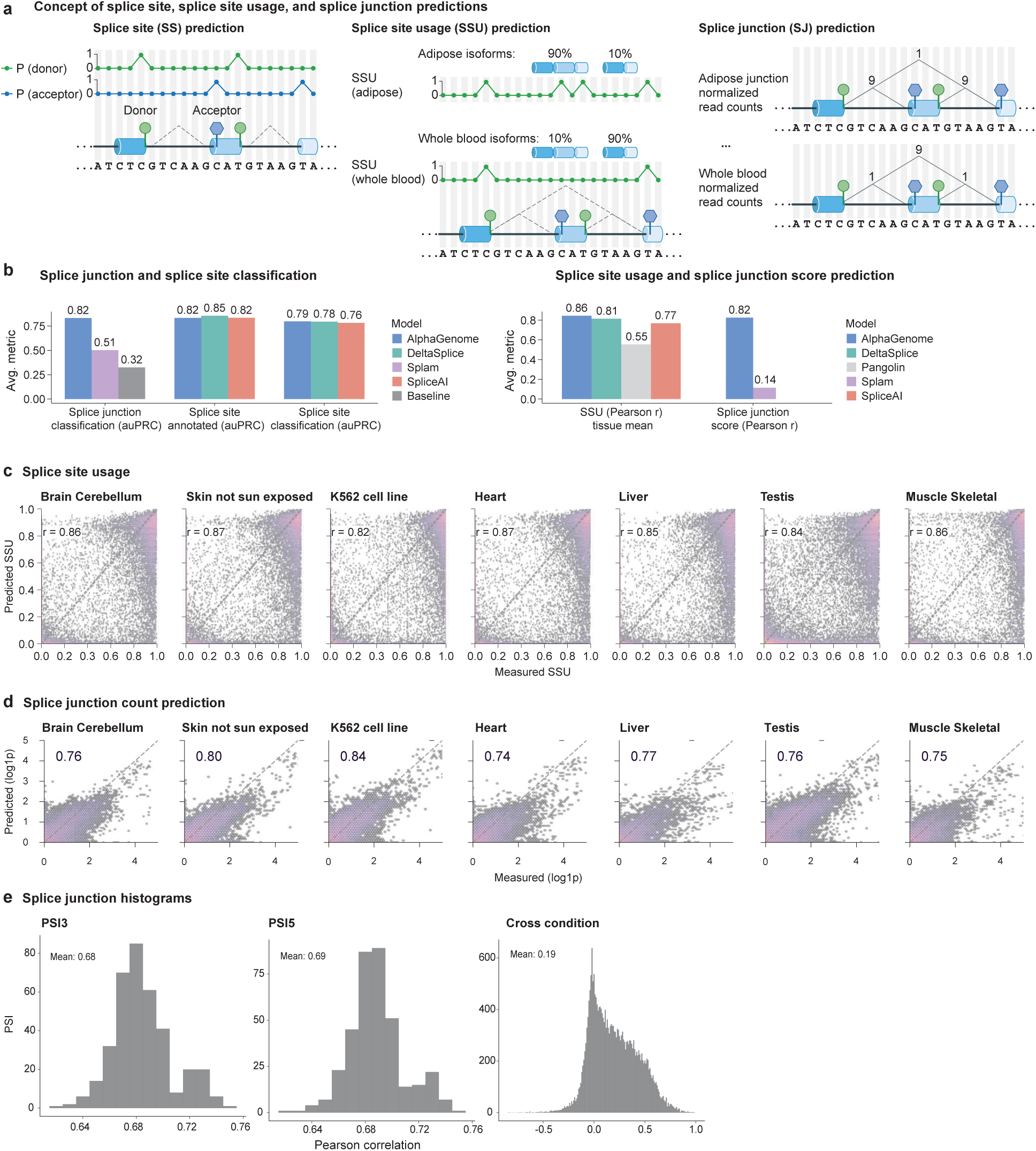
Splicing track performance. **(a)** Schematic overview of splice site (SS) classification, splice site usage (SSU) prediction, and splice junction read count (SJ) prediction tasks. **(b)** (left) Performance comparison (AUPRC) of SS classification and SJ classification against reference methods. ‘Baseline’ means fraction of positive splice junctions in the evaluated data. Splice site classification is evaluated with both GTF (GENCODE v46) annotated splice sites only, and also splice sites derived from GTEx RNA-seq data (Methods). Splice junction classification is to classify true splice junctions observed from RNA-seq versus false junctions not observed from RNA-seq (but the splice sites are observed). Splice junction classification was evaluated per tissue and then the mean AUPRC across tissues were reported. (right) Performance comparison (Pearson r) of predicted vs. measured SSU and SJ counts (log(1 + 𝑥) transformed). This metric is computed per tissue and the mean across tissues were reported. **(c)** Scatter plot between predicted and measured donor SSU across seven example human tissues (from GTEx). Pearson r in each tissue is displayed as text. **(d)** Scatter plot between predicted and measured splice junction counts across seven human tissues (from GTEx). Pearson r in each tissue is displayed as text. **(e)** Distribution of Pearson correlation coefficients between predicted and measured PSI3 per tissue (left), PSI5 per tissue (middle), and junction counts across tissues (measure tissue specificity of splice junction predictions).

**Extended Data Figure 3.**
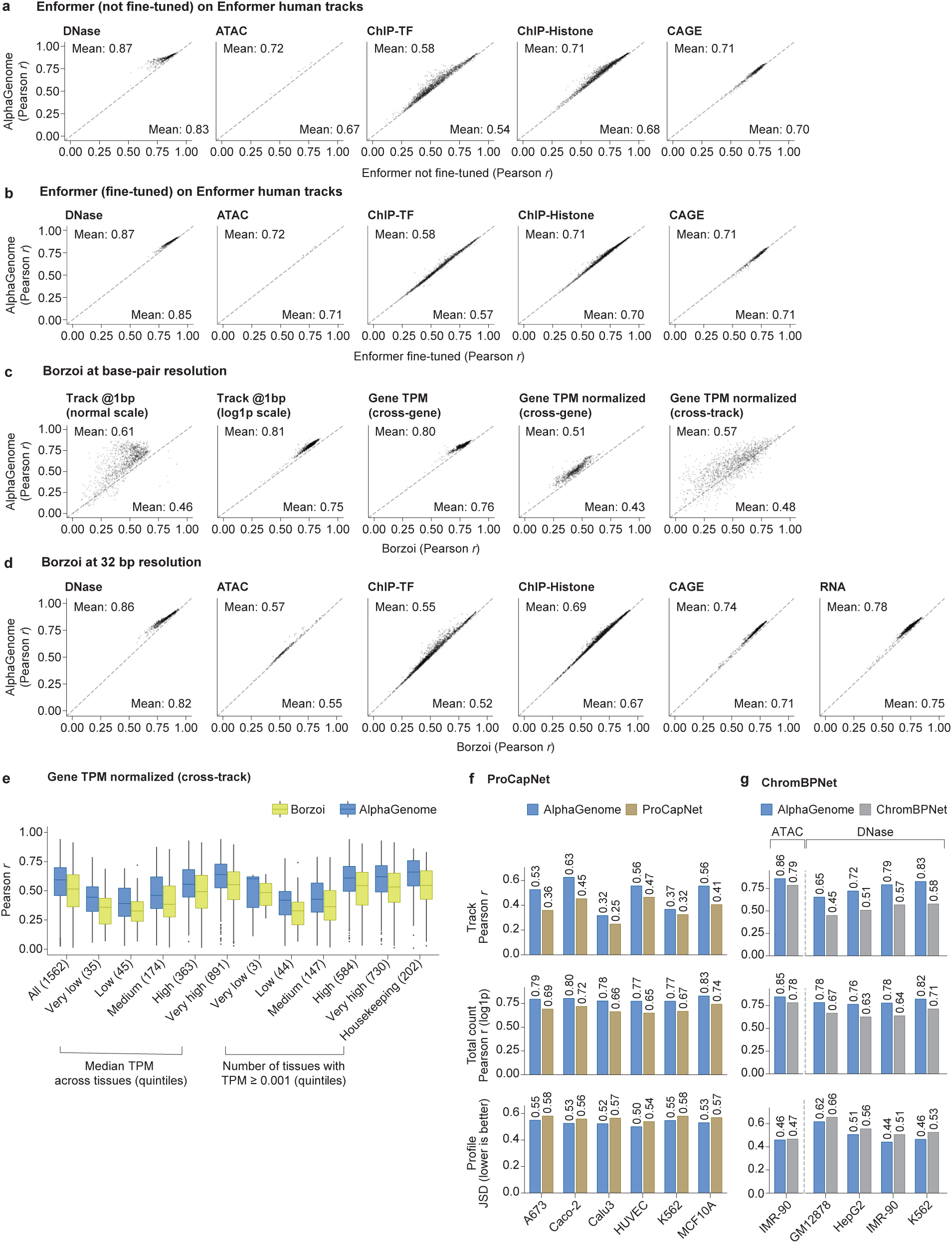
Track-level performance benchmarking. Performance comparison of AlphaGenome with Enformer and Borzoi on held-out genomic track prediction. **(a, b)** Comparison of AlphaGenome test set performance on Enformer human tracks (each dot is one track) against Enformer models either (a) not fine-tuned or (b) fine-tuned on human data (the main released Enformer version). AlphaGenome model was re-trained for direct comparability using matched training intervals and an additional Enformer prediction head (see Methods). **(c)** Evaluation of RNA-seq prediction performance at base and gene resolution using the same source of RNA-seq data as Borzoi, but processed at base-resolution and not scaled (see Methods). Borzoi’s 32 bp RNA-seq predictions were upsampled and unscaled to the original scale for comparison. The larger performance difference observed on the normal scale (first column) likely reflects resolution differences at exon-intron boundaries. This difference decreased when using log(1 + 𝑥) transformed values (second column), suggesting better agreement on overall gene expression levels. A similar trend was observed when aggregating expression per gene (average exon coverage, third column). Cell-type specificity was evaluated by correlating quantile-normalized, mean-subtracted expression profiles across genes (fourth column) and across tracks (fifth column). **(d)** Test set performance comparison of AlphaGenome against Borzoi (fold 1) on Borzoi track data at 32 bp resolution (each dot is one track). AlphaGenome was fine-tuned with an additional ‘Borzoi head’ at matched resolution (see Methods). **(e)** Stratification of cell-type specific prediction accuracy. The per-gene log-fold change correlation performance (from panel c, fourth column) was stratified by gene characteristics: median expression level across tissues (Median TPM; quintile breakpoints: 5.5 *×* 10*^−^*^9^, 4.1 *×* 10*^−^*^4^, 8.1 *×* 10*^−^*^4^, 0.17, 4.1, 3.6 *×* 10^4^ TPM), number of tissues with the gene expressed (TPM *≥* 0.001; quintile breakpoints: 9.4 *×* 10*^−^*^8^, 9.4 *×* 10*^−^*^4^, 8.0, 52, 54, 54 tissues), and housekeeping gene status. **(f, g)** Performance comparison of AlphaGenome against (f) ProCapNet (on PRO-Cap data) and (g) ChromBPNet (on ATAC and DNase). Evaluation was performed on ProCapNet fold 5 and ChromBPNet fold 0 test peak regions, respectively, where regions overlapping with AlphaGenome fold 0 training intervals were excluded. Performance is quantified by track Pearson r, Pearson r on the log total count, and Jensen-Shannon distance (JSD; lower indicates better performance). AlphaGenome outperforms the baselines across all metrics, modalities and cell-lines. For (g), only tracks with matching experiment accessions between AlphaGenome and ChromBPNet training sets were considered.

**Extended Data Figure 4.**
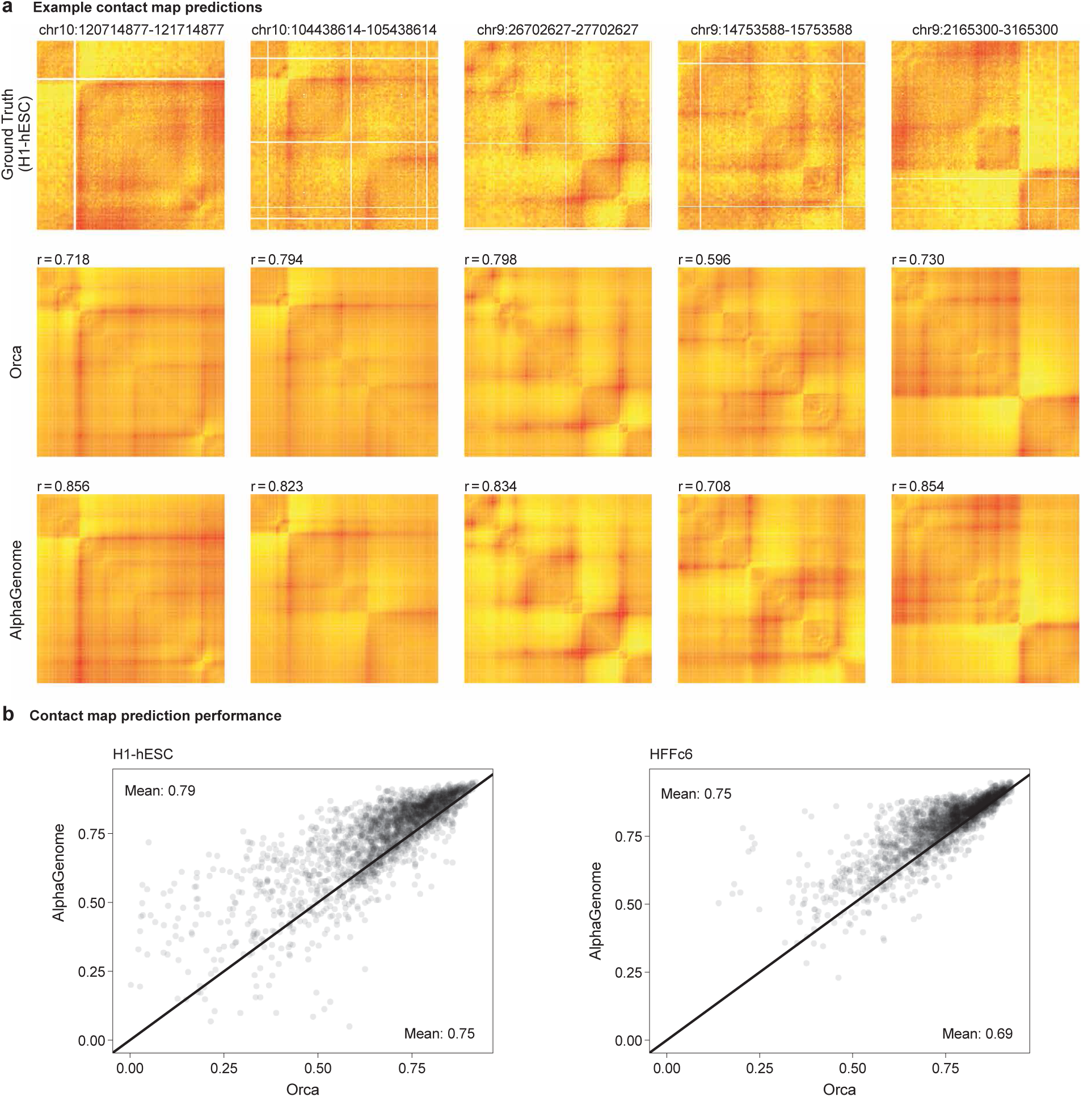
Contact map prediction and comparison with Orca baseline. **(a)** Comparison of predicted chromatin contact maps for five randomly selected 1Mb intervals from the validation set. Rows show ground truth experimental data (H1-hESC Micro-C), predictions from the baseline model Orca, and predictions from AlphaGenome. Pearson correlation coefficients (R) between prediction and ground truth are displayed in the upper right corner of each corresponding predicted map. **(b)** Scatterplot comparing overall contact map prediction performance of AlphaGenome and Orca across all 1Mb test intervals, for the two cell-types predicted by Orca. Each point represents an interval, plotting the Pearson R achieved by AlphaGenome (y-axis) versus Orca (x-axis) when comparing predictions to the ground truth. Points above the diagonal indicate intervals where AlphaGenome achieved higher correlation than Orca.

**Extended Data Figure 5.**
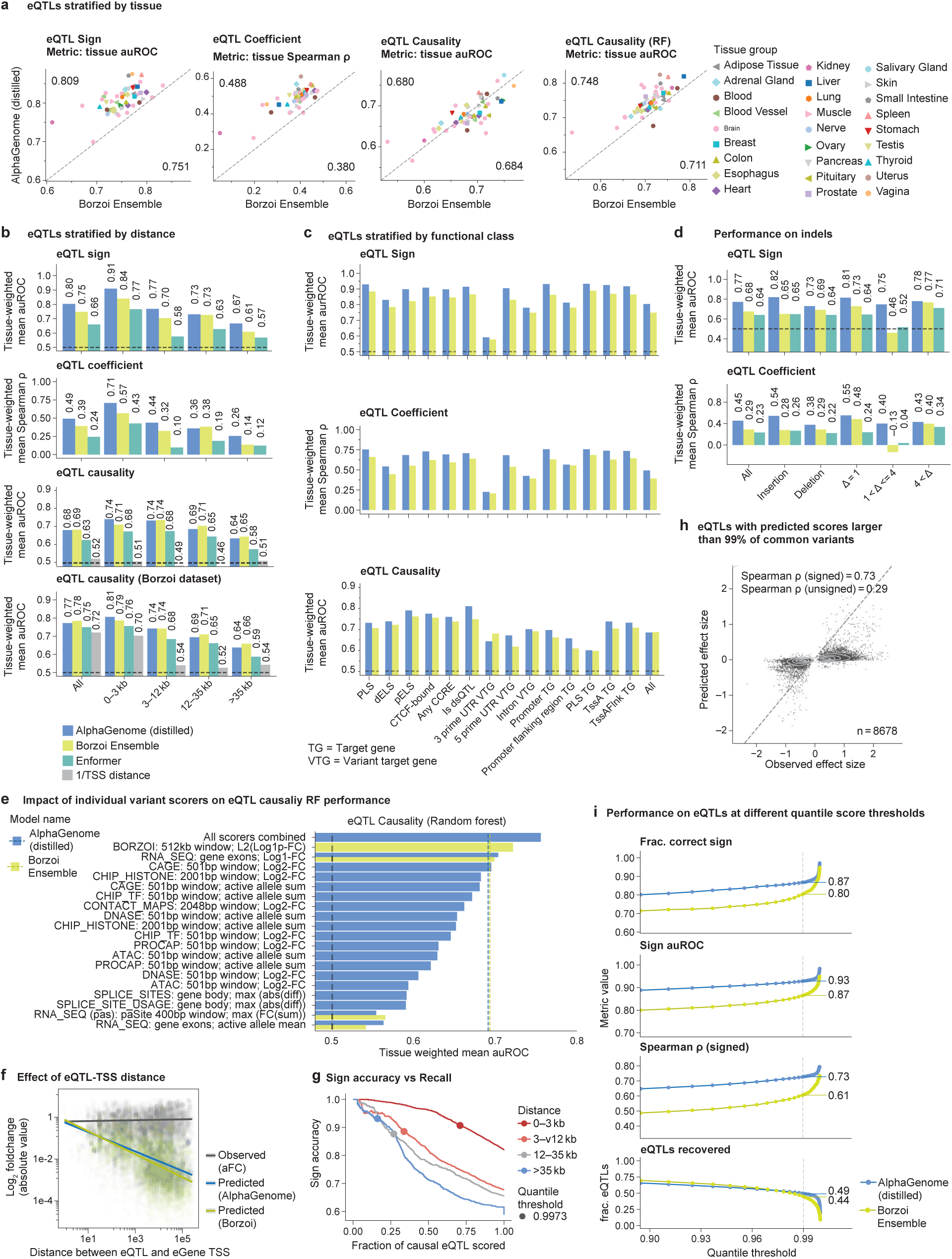
Additional eQTL analysis. Further characterization and stratification of AlphaGenome’s eQTL prediction performance. **(a)** Performance comparison on eQTL tasks (Coefficient Spearman R, Sign auROC, Causality auROC zero-shot, Causality RF auROC) across individual GTEx tissues (see Methods for specific scorer configurations). For zero-shot tasks, AlphaGenome’s default variant scorer is used (Methods; Fig. 4a). For the supervised causality task (’eQTL Causality (RF)’), results use the best-performing feature set (’All scorers combined’; (e)). For Borzoi comparison, the published L2_DIFF scorer across all output tracks is used. **(b)** Performance on eQTL tasks (Coefficient, Sign, Causality) stratified by distance from variant to target gene TSS, compared to Borzoi and a 1/distance baseline for the causality task. **(c)** Performance on eQTL tasks stratified by broad functional classes based on variant location. TG = Target Gene; VTG = Variant target gene. **(d)** Performance on indel eQTLs for the Sign and Coefficient tasks stratified by indel size, comparing AlphaGenome and Borzoi. **(e)** Random forest performance on the eQTL causality task using different feature sets. Horizontal bar plot compares RF performance (auROC from Fig. 4g) when trained using features derived from all scorers combined or individual modality scorers (also comparing AlphaGenome to Borzoi outputs). **(f)** Observed versus predicted eQTL log2 fold change plotted against variant-TSS distance (within 262 kb, ensuring target TSS is within receptive field context). Observed values are allelic log-fold changes from^1^ were used to allow direct comparison with the scale of AlphaGenome’s predicted log-fold changes (Methods). **(g)** Sign accuracy as well as the fraction of eQTLs that pass a certain quantile-score cutoff visualized for all possible thresholds and four eQTL distance categories. The quantile score cutoff which leads to an overall (across distance-bins) sign accuracy of 90% is highlighted. At this cutoff, sign accuracy is similar across distance categories, although recall is notably lower for distal variants. **(h)** Scatterplot comparing observed versus predicted effect sizes for 8,678 fine-mapped eQTLs filtered for high AlphaGenome prediction scores (>99th percentile of common variants). **(i)** Relationship between AlphaGenome’s quantile score threshold (x-axis; Methods) and the resulting performance (y-axis) for eQTL Sign accuracy and Coefficient Spearman R tasks.

**Extended Data Figure 6.**
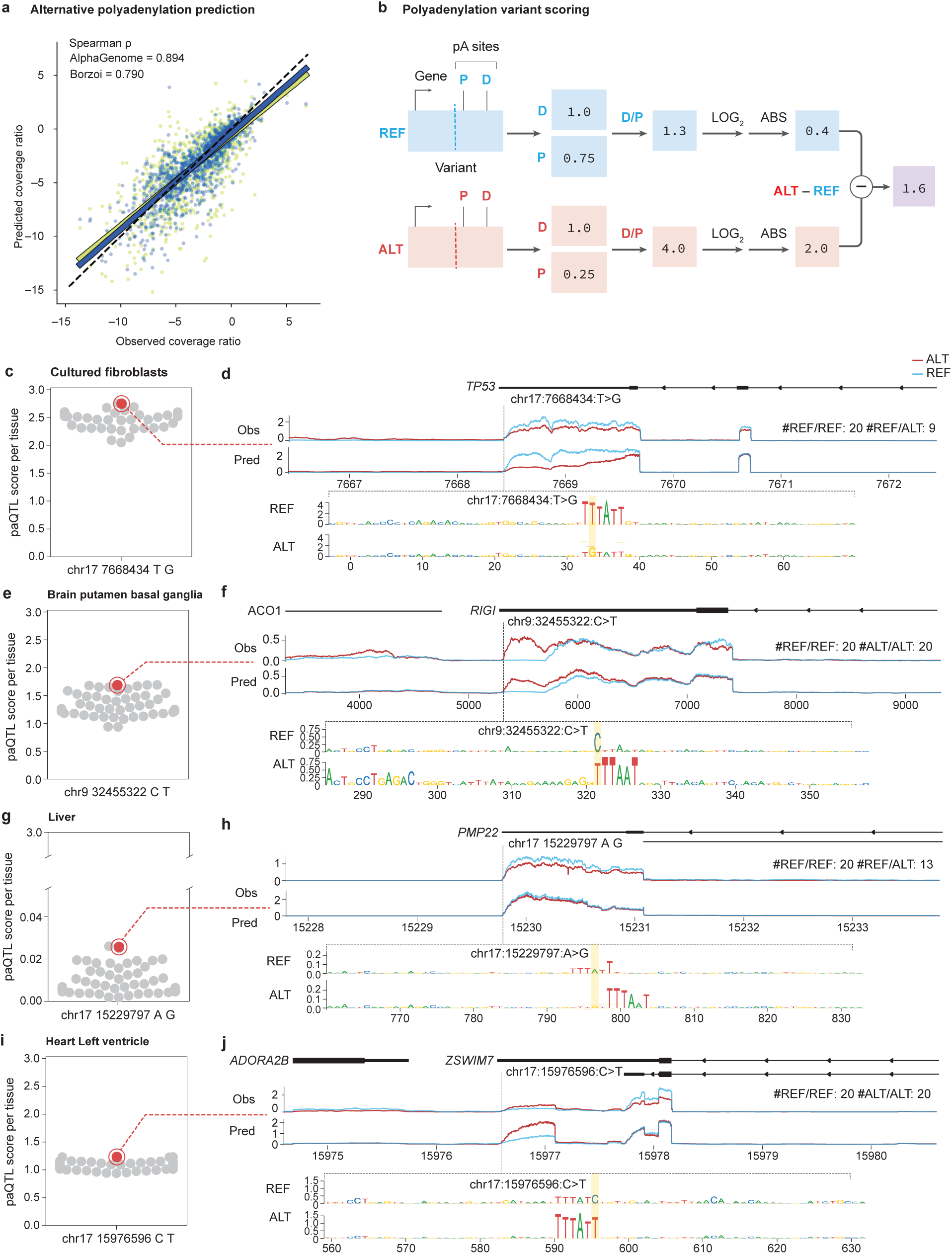
Improved prediction of 3’ UTR polyadenylation and paQTLs with AlphaGenome. Polyadenylation predicted vs observed coverage ratio (COVR) for each gene’s most distal and proximal PAS for Borzoi and AlphaGenome. Polyadenylation variants scoring scheme. **(d,e)** Example of variant disrupting a polyadenylation motif in the TP53 gene. (d) Distribution of scores for the indicated variant across GTEx tissues (gray swarmplot) with the highest and lowest scoring tissues and average score highlighted (red dots). (e) Observed and predicted RNAseq for the tissues highlighted in (d), and in silico saturation mutagenesis scores (ISM) of the 72 bp flanking the variant. ISM clearly highlights the relevance of the polyadenylation motif exclusively in the reference background, where the variant does not compromise the motif. **(f,g)** As (d) and (e) but for a variant generating a new polyadenylation motif in the RIGI gene. **(h,i)** As (d) and (e) but for a paQTL with a low variant score. The variant disrupts the PAS but a cryptic one emerges nearby with limited effect on gene expression. **(j,k)** As (d) and (e) but for a failure case. AlphaGenome correctly identifies the emergence of a novel PAS but fails to correctly predict the effect on gene expression.

**Extended Data Figure 7.**
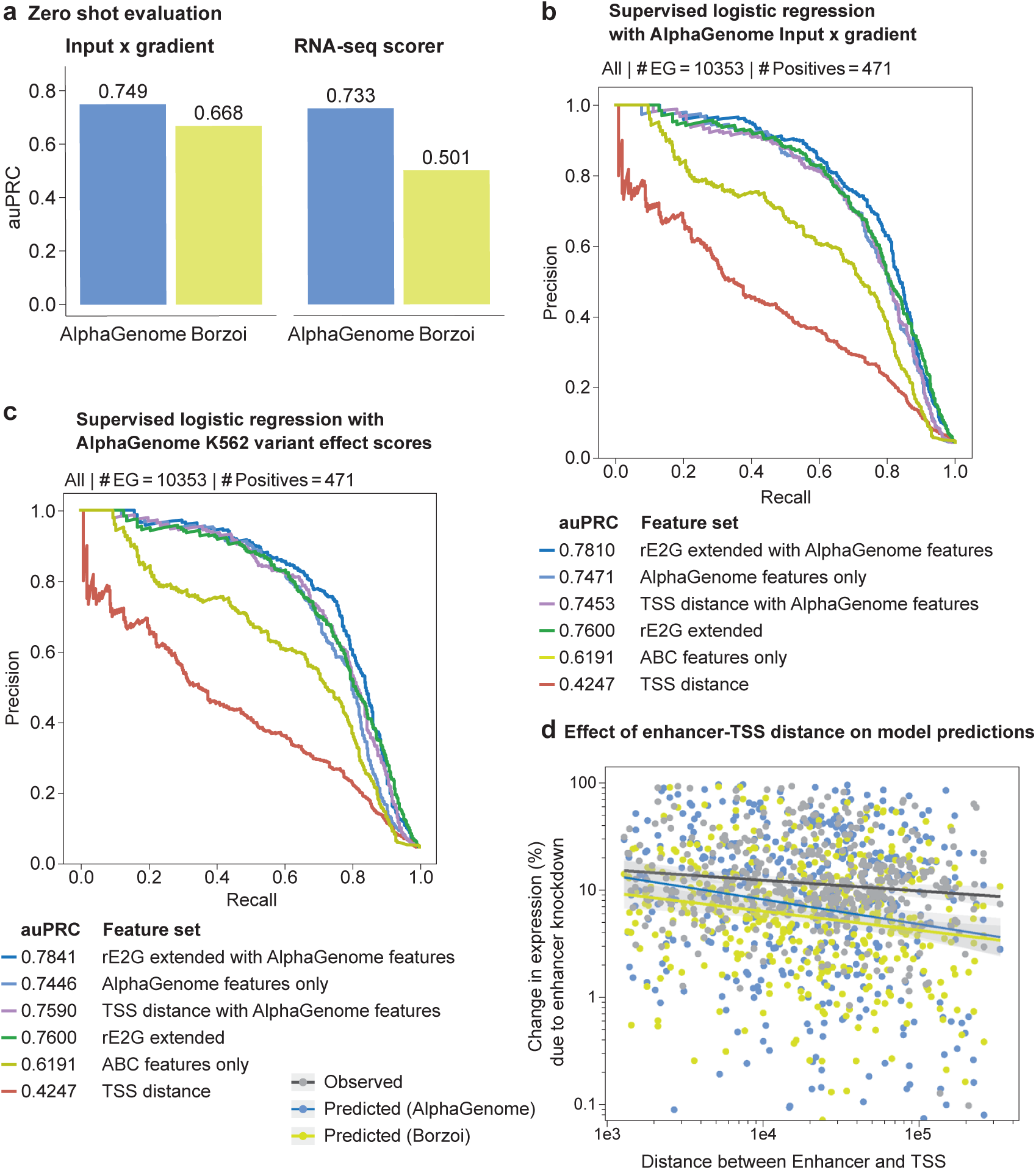
AlphaGenome improves enhancer-gene linking using input gradients and shows enhanced sensitivity to distal enhancers. **(a)** Zero-shot performance of AlphaGenome and Borzoi on the ENCODE-rE2G benchmark. Bars indicate the area under the precision-recall curve (auPRC) for predicting enhancer-gene links. Two scoring methods derived from each model were evaluated: input gradient scores and RNA-seq variant effect scores. **(b)** Impact of incorporating AlphaGenome’s input gradient score as a feature in the ENCODE-rE2G extended logistic regression model, evaluated on the ENCODE-rE2G benchmark. ENCODE-rE2G is a logistic regression model trained to predict enhancer-gene interactions from features^2^. Precision-recall curves are shown, colored by the feature sets used for training the regression model (auPRC values indicated in the legend). Feature sets are: • *rE2G extended with AlphaGenome features*: All ENCODE-rE2G extended model features plus a single AlphaGenome’s input x gradient score. • *AlphaGenome features only* : The AlphaGenome input x gradient score alone. • *TSS distance with AlphaGenome features*: AlphaGenome input x gradient score plus the distance to TSS feature. • *rE2G extended* : All features from the ENCODE-rE2G extended model^2^. • *TSS distance*: Distance to TSS feature from^2^. • *ABC features only* : Subset of ’rE2g extended’, with only features related to the Activity-By-Contact (ABC) model^2^. **(a)** Precision-recall curves for the ENCODE-rE2G benchmark, similar to panel (b), evaluating the ENCODE-rE2G extended regression model with different feature sets. Area under the precision-recall curve (auPRC) values for the different feature sets are indicated in the legend. In this configuration, ‘AlphaGenome features’ consist of a more comprehensive set of K562 cell line-specific variant effect scores. These include Allele-Specific Activity Scores (AAS) and variant effect scores calculated as the difference between alternate (ALT) and reference (REF) allele predictions (ALT-REF Diff scores). These scores were derived from AlphaGenome for the following genomic assays: • RNA-seq of the target gene • ChIP-TF EP300 • ChIP-Histone H3K27ac • CAGE • PRO-cap • H1-ESC contact maps **(b)** Relationship between enhancer perturbation effects (ENCODE-rE2G dataset^2^) and enhancer-promoter distance. The scatter plot shows experimentally observed percentage changes in gene expression upon enhancer knockout (grey points and trend line) versus the genomic distance between the enhancer and the target gene’s Transcription Start Site (TSS). Overlaid are trend lines for AlphaGenome’s (AG, dark blue) and Borzoi’s (green) predictions of these expression changes, derived from their respective model input gradient scores. Each point corresponds to a validated enhancer-gene pair

**Extended Data Figure 8.**
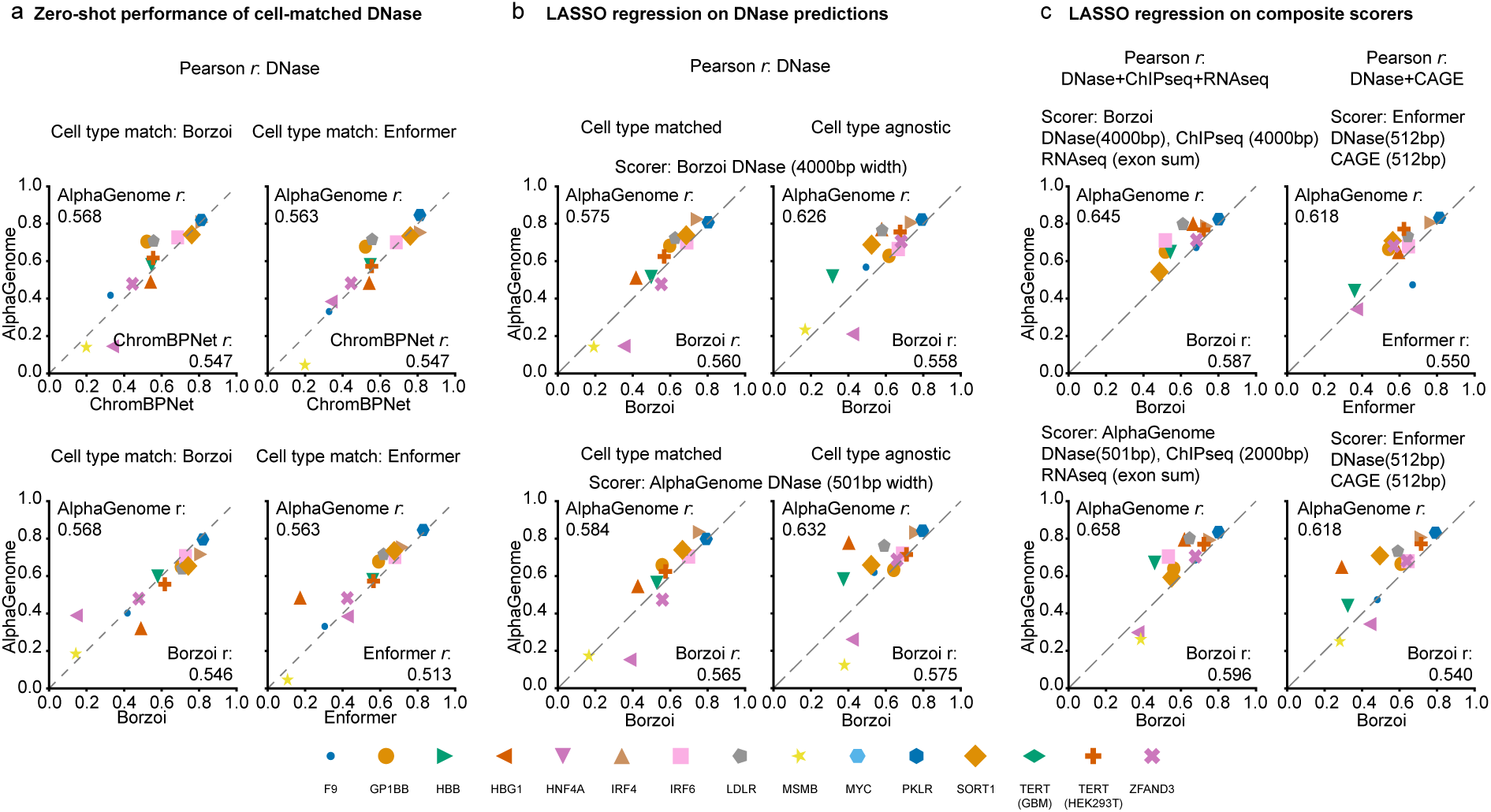
AlphaGenome performance on the CAGI5 MPRA challenge. Comparison of models predicting the effects of regulatory variants on gene expression using CAGI5 MPRA data. **(a)** Pairwise comparisons of zero-shot, cell-type matched performance using raw DNase model outputs. Scatter plots show per-gene or per-context Pearson correlations. AlphaGenome shows comparable or slightly improved performance compared to Enformer, ChromBPNet, and Borzoi Ensemble based on overall correlations across compared loci (details in plot annotations; cell-type matching strategy adapted from Enformer/Borzoi papers, see Methods). Note that ChromBPNet only reports performance for TERT-HEK293T, thus TERT-GBM is excluded in the AlphaGenome vs ChromBPNet comparison. **(b)** Pairwise comparisons of cell-type matched (see a) and cell type agnostic performance using LASSO regression on spatially summed DNase features (Methods). Comparison of the original Borzoi scorer (upper row) and AlphaGenome recommended scoring strategy (bottom row). AlphaGenome outperforms Borzoi in all settings and shows the lack of need for cell type matching when applying lasso regression. **(c)** Pairwise comparisons of cell-type agnostic performance using LASSO regression with different input features: left column shows the composite scorer described in Borzoi (DNase + Histone ChIPseq + RNA), right column sows the composite scorer described in Enformer (DNase + CAGE). AlphaGenome generally outperforms Enformer and Borzoi for all input feature settings.

**Extended Data Figure 9.**
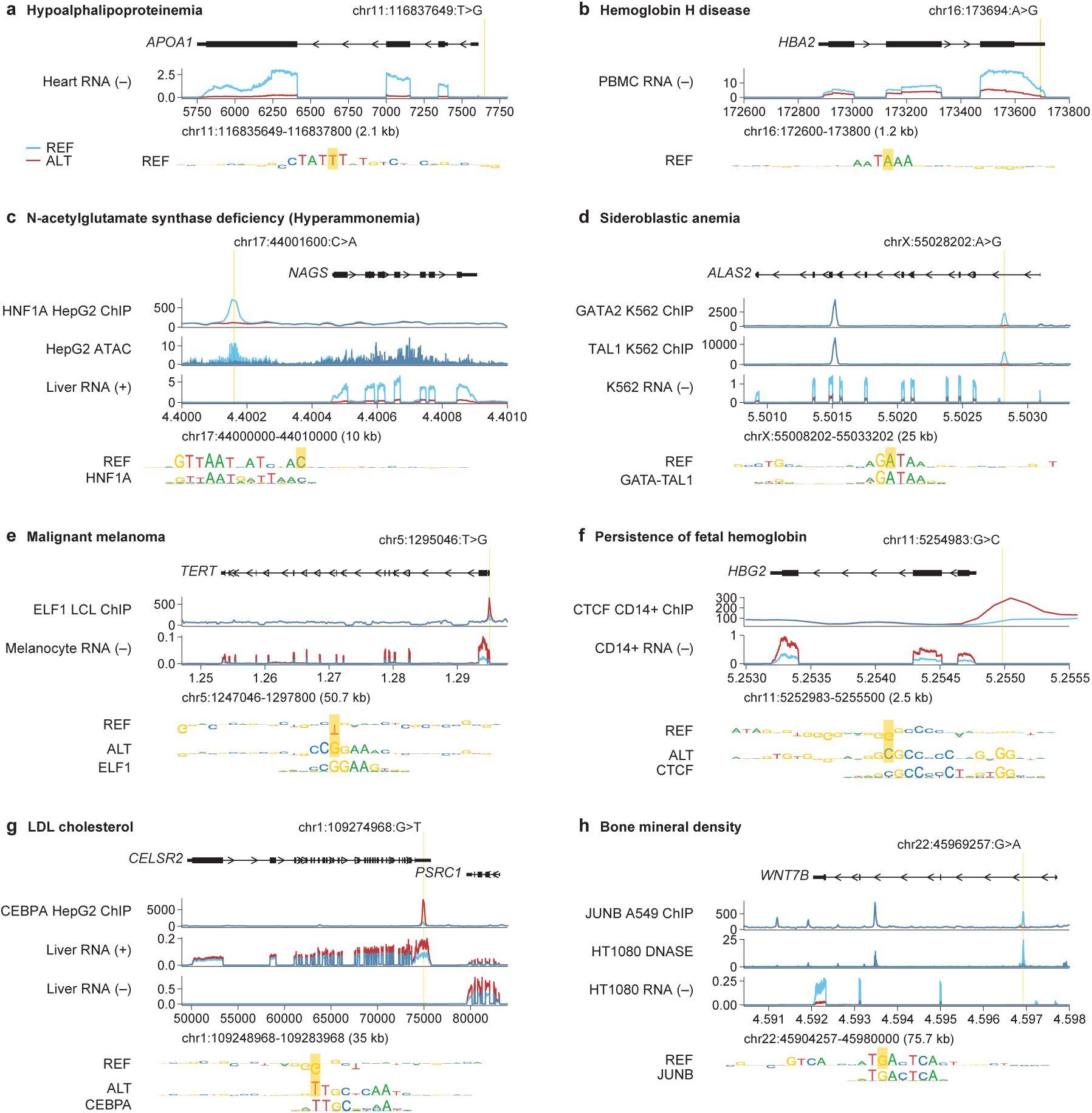
Interpreting trait-affecting variants using AlphaGenome. **(a)** AlphaGenome predictions for the variant chr11:116837649:T>G, previously reported to result in underexpression of APOA1 and Hypoalphalipoproteinemia^3^ . The variant strongly reduces the predicted expression of the APOA1 transcript in heart tissue. ISM of the reference sequence suggests that the variant disrupts a TATA-like motif. **(b)** AlphaGenome predictions for the variant chr16:173692:A>G, previously reported to cause Hemoglobin H disease^4^. The model predicts that the variant strongly reduces the predicted expression of the HBA2 transcript. ISM of the reference sequence suggests that the variant disrupts the polyadenylation hexamer. **(c)** AlphaGenome predictions for the variant chr17:44001600:C>A, previously reported to result in reduced binding of HNF1, underexpression of NAGS and ultimately N-acetylglutamate Synthase Deficiency^5^. AlphaGenome predicts reduced binding of HNF1A, local DNA accessibility, as well as a strong reduction in expression of the NAGS transcript in liver and related cell types. ISM of the reference sequence suggests that the variant disrupts a HNF1 motif. **(d)** AlphaGenome predictions for the variant chrX:55028202:A>G, previously reported to result in reduced GATA binding and reduced expression of ALAS2, thereby possibly causing Sideroblastic Anemia^6^. The model predicts reduced binding of GATA factors (GATA2 shown), as well as TAL1, and drastically reduced expression of the ALAS2 transcript in K562. ISM of the reference sequence suggests that the variant disrupts a GATA-TAL composite motif. **(c)** AlphaGenome predictions for the variant chr5:1295046:T>G, previously reported to cause TERT overexpression associated with malignant melanoma^7^. AlphaGenome predicts increased binding of ETS factors (ELF1 shown) and a significant increase in TERT expression in melanocytes. Comparing the ISM around the reference and alternative alleles suggests that the variant creates an ETS factor binding motif. **(d)** AlphaGenome predictions for the variant chr11:5254983:G>C, previously implicated in hereditary persistence of fetal hemoglobin^8^. The model predicts increased CTCF binding, as well as increased HBG2 expression (shown for CD14+ monocytes). Comparing the ISM around the reference and alternative alleles suggests that the variant creates a CTCF binding motif. **(g)** AlphaGenome predictions for the variant rs12740374, previously found to be associated with low-density lipoprotein cholesterol levels^9^. Additionally, the variant is a plasma pQTL for CELSR2, as well as GTEx eQTL for CELSR2 and PSRC1^10,11^. Consistent with the observed molecular QTL effects, AlphaGenome predicts increased expression of both of these genes in liver tissue. Moreover the model predicts increased CEBP binding on the alternative allele. Comparing the ISM around the reference and alternative alleles suggests that the variant creates a CEBP binding motif. **(h)** AlphaGenome predictions for the variant, rs570639864 previously found to be associated with reduced bone mineral density^12^. AlphaGenome predicts decreased binding of JUN/FOS factors (JUNB shown), as well as a strong reduction in expression of WNT7B, a gene implicated in bone formation^13^. ISM of the reference sequence suggests that the variant disrupts a JUN/FOS motif.

**Supplementary Figure 1.**
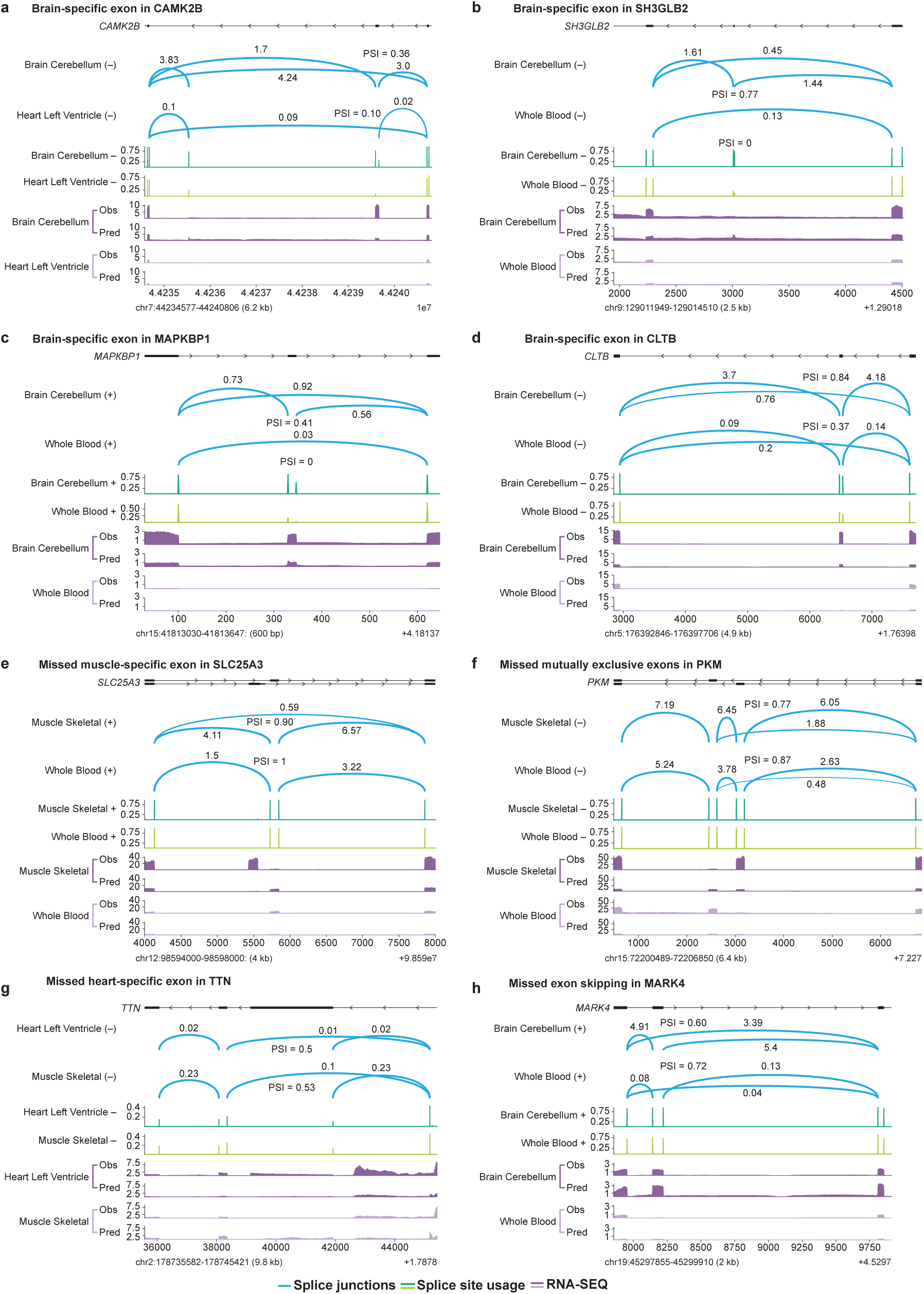
Examples of AlphaGenome predictions for tissue-specific alternative splicing. **(a)** Panels illustrate both successful predictions and specific challenges/failure modes. Coordinates are hg38. AlphaGenome generally predicts differential splice site usage and splice junction patterns across tissues, with failures typically involving missed predictions of specific tissue-regulated exon inclusion or skipping events. Measured data from GTEx. Predicted and observed RNA-seq, predicted splice site usage, and splice junctions surrounding an alternative exon (chr7:44239589-44239663) in CAMK2B, exhibiting differential splicing between neuronal and non-neuronal tissues^14^. **(b-d)** Additional examples of predicted splicing patterns around alternative exons exhibiting neuronal vs non-neuronal splicing^14^ in: (b) SH3GLB2 (chr9:129013008-129013019), (c) MAPKBP1 (chr15:41813330-41813347), and (d) CLTB (chr5:176396479- 176396532). **(e)** Failure case: Predictions surrounding an alternatively spliced exon in SLC25A3. AlphaGenome fails to predict the observed inclusion of the muscle-specific exon. **(f)** Failure case: Predictions surrounding mutually exclusive exons in PKM. AlphaGenome erroneously predicts the inclusion of both exons in Muscle and Blood tissues where only one is expected. **(g)** Failure case: Predictions surrounding exons 49 and 50 of TTN. AlphaGenome fails to predict the observed inclusion of the heart-specific exon 49. **(h)** Failure case: Predictions surrounding an alternative exon (chr19:45298143-45298222) in MARK4. AlphaGenome fails to predict the observed exon skipping event in Blood relative to neuronal tissues^14^.

**Supplementary Figure 2.**
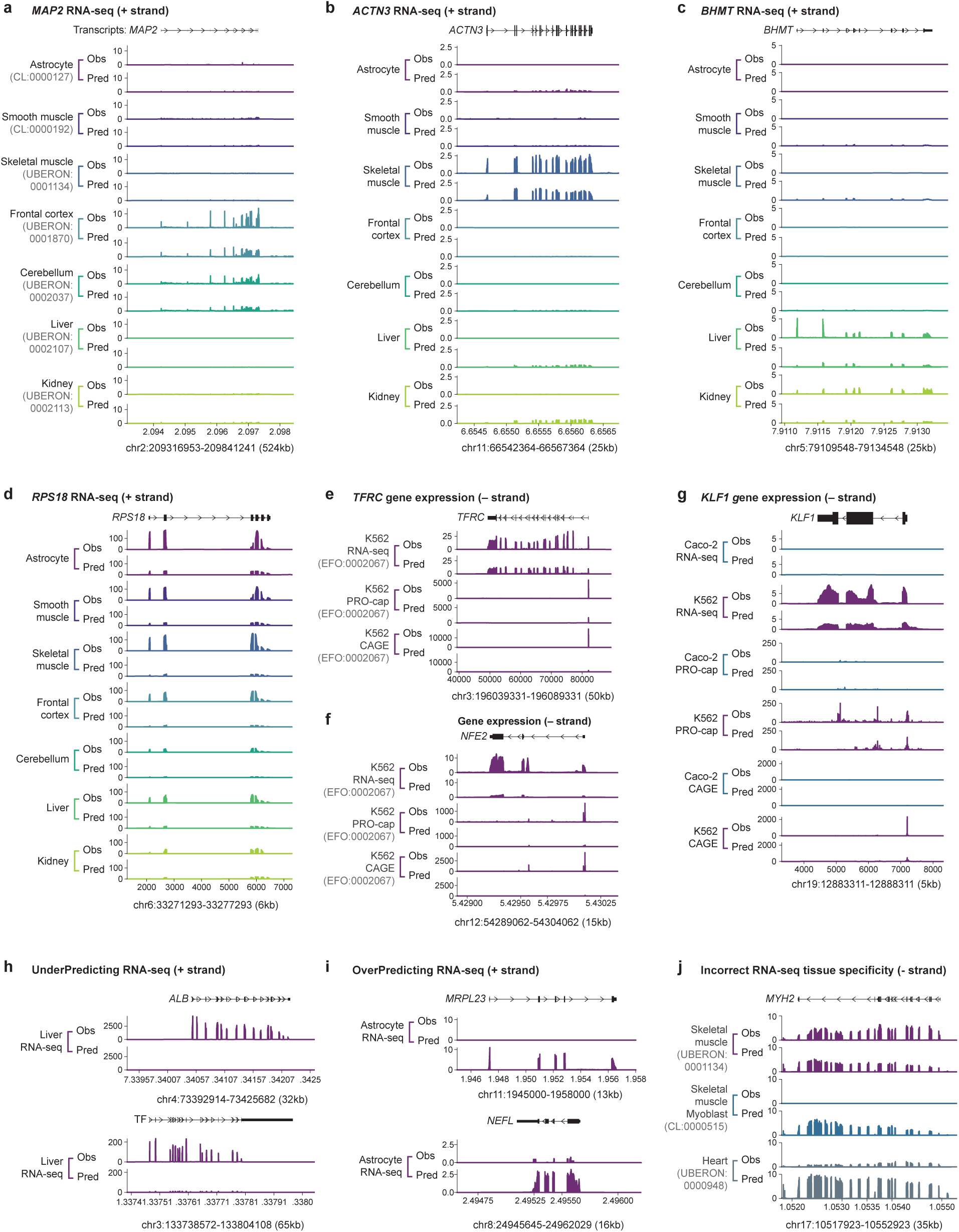
Examples of AlphaGenome gene expression predictions. Observed (obs) versus predicted (pred) signals for various expression-related assays across different genomic loci (hg38 coordinates) and tissues/cell types, illustrating both successful predictions and limitations. AlphaGenome generally captures expression patterns across multiple expression assays (RNA-seq, CAGE, PRO-cap), with failures typically involving underprediction of signal magnitude and errors in tissue specificities. **(a)** Accurate prediction of *MAP2* (Microtubule Associated Protein 2) expression in brain tissues (frontal cortex, cerebellum) but absence in astrocytes. **(b)** Accurate prediction of *ACTN3* (Actinin Alpha 3) expression specifically in skeletal muscle tissue. **(c)** Accurate prediction of *BHMT* (Betaine-Homocysteine S-Methyltransferase) expression in liver and kidney tissues. **(d)** Prediction for the housekeeping gene *RPS18* (Ribosomal Protein S18). The RNA-seq expression patterns are largely correct, but signal magnitude is underestimated across tissues. **(e)** Example of multi-output type expression analysis (RNA-seq, CAGE, PRO-cap shown) for *TFRC* (Transferrin Receptor) in K562 cells. Predicted signal output shapes are accurate but magnitudes are underestimated. **(f)** Additional multi-output type expression analysis example for *NFE2* (Nuclear Factor, Erythroid 2) in K562 cells. **(g)** Multi-output and multi-tissue expression analysis for *KLF1* (KLF Transcription Factor 1), correctly predicting high expression in K562 but absence in Caco-2 cells. **(h)** Failure cases: Significant underprediction of RNA-seq signal for liver-specific genes *ALB* (Albumin) and *TF* (Transferrin). **(i)** Failure cases: Significant overprediction of *MRPL23* (Mitochondrial Ribosomal Protein L23) and *NEFL* (Neurofilament Light Chain) RNA-seq signal in astrocytes. **(j)** Failure case: Incorrect tissue-specificity predicted for *MYH2* (Myosin Heavy Chain 2), showing slight underprediction in skeletal muscle alongside overprediction in skeletal muscle myoblasts and heart tissue.

**Supplementary Figure 3.**
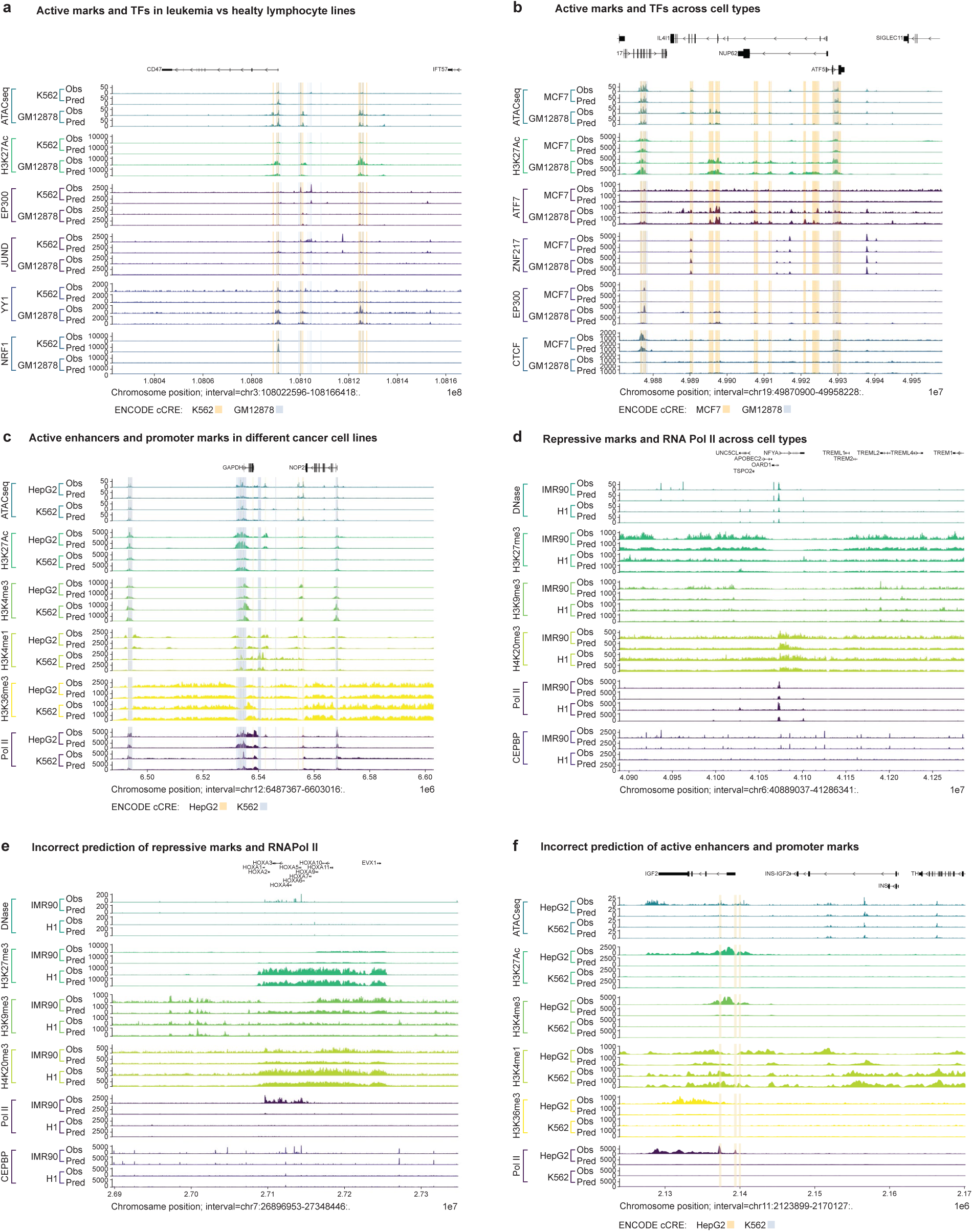
AlphaGenome prediction examples for chromatin tracks. Comparison of predicted (pred) and observed (obs) epigenetic features and transcription factor (TF) binding profiles on held-out genomic locations (hg38), illustrating capabilities and limitations. AlphaGenome is broadly accurate at predicting diverse chromatin landscapes, including accessibility, histone modifications, and protein binding, often capturing cell-type specific patterns seen across lineages. Observed prediction failures typically involve underpredicting expected signal magnitude or presence in specific contexts, rather than placing marks in incorrect locations. **(a)** Differential regulatory landscape at the *CD47* locus between K562 (leukemia) and GM12878 (lymphoblastoid) cell lines. Despite shared lymphoid origin and *CD47* expression, AlphaGenome correctly captures cell-line-specific differences in TF binding, EP300 DNA-binding cofactor occupancy, and histone modifications in the proximal regulatory region. **(b)** Cell-type-specific predictions at the *IL4I1* (Interleukin 4 Induced 1) locus. AlphaGenome accurately recapitulates chromatin accessibility and TF binding in GM12878 versus MCF7 (breast cancer) cells. **(c)** Active transcription marks at the ubiquitously expressed housekeeping gene *GAPDH* (Glyceraldehyde-3-Phosphate Dehydrogenase). AlphaGenome correctly predicts patterns for both broad and sharp active chromatin marks, as well as RNA Polymerase II binding. **(d)** Prediction of an active *NFYA* (Nuclear Transcription Factor Y Subunit Alpha) promoter region embedded within heterochromatin. AlphaGenome accurately predicts the overall chromatin landscape, including broad flanking repressive marks. **(e)** Failure case: Developmentally regulated *HoxA* (Homeobox A) cluster. AlphaGenome correctly predicts the repressed state in H1 human embryonic stem cells but fails to capture the activation of the anterior part of the cluster in IMR90 fibroblasts. **(f)** Failure case: Liver-specific gene regulation at the *IGF2* (Insulin Like Growth Factor 2) locus. While correctly predicting inactivity in non-expressing K562 cells, AlphaGenome fails to predict expression-associated histone modifications and RNA Polymerase II binding in expressing HepG2 liver cancer cells.

**Supplementary Figure 4.**
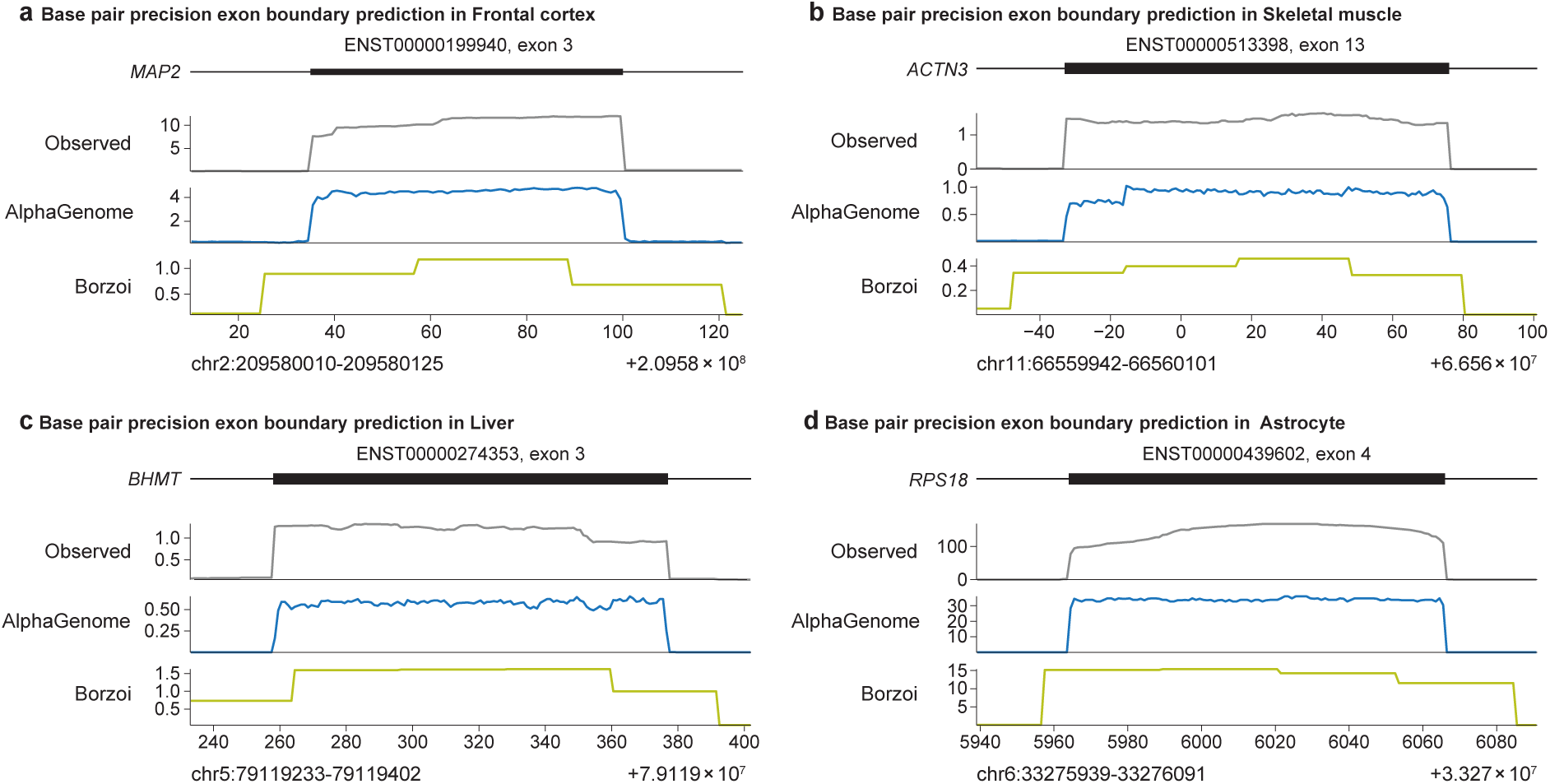
AlphaGenome enables single base-pair predictions of RNA-seq coverage at exon-boundaries. **(a)** Observed and predicted RNA-seq coverage for exon 3 of the *MAP2* gene in frontal cortex tissue (same gene as in S2.2a). AlphaGenome correctly predicts the exon boundaries. Borzoi predicts RNA-seq coverage in 32 bp bins, preventing exact inference of the beginning and end of the exon. **(b)** Observed and predicted RNA-seq coverage for exon 13 of the *ACTN3* gene in skeletal muscle tissue (c.f. S2.2b). **(c)** Observed and predicted RNA-seq coverage for exon 3 of the *BHMT* gene in liver tissue (c.f. S2.2c). **(d)** Observed and predicted RNA-seq coverage for exon 4 of the *RPS18* gene in astrocytes (c.f. S2.2d).

**Supplementary Figure 5.**
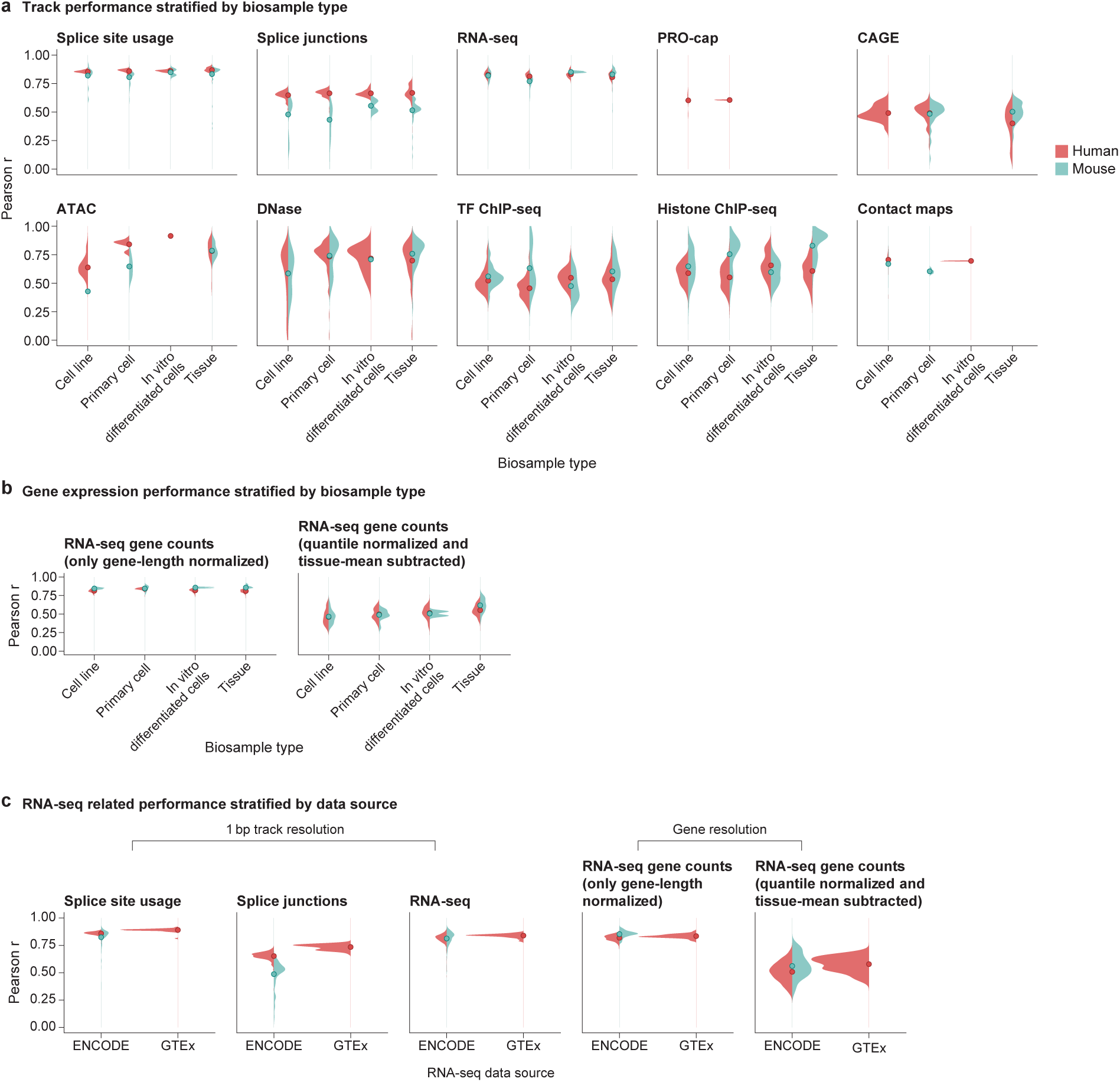
Stratification of model performance supplementary to main. Fig. 2c**,d**. Model performance evaluation metrics stratified by metadata variables. **(a)** Track correlation performance (Pearson r, related to Fig. 2c) stratified by biosample type (cell line, primary cell, in vitro differentiated cells, tissue) for Human (red) and Mouse (blue). Performance is broadly consistent across types for most assays, with notable exceptions including lower performance observed for splice junctions in mouse primary cells, and for DNase-seq and ATAC-seq in cell lines compared to tissues. **(b)** Gene expression correlation performance (Pearson r, related to Fig. 2d) stratified by biosample type. Performance on both raw and normalized/mean-subtracted gene expression metrics remains generally consistent across different biosample types for both species. **(c)** Performance comparison for RNA-seq derived metrics (Pearson r, related to Fig. 2c,d) stratified by data source (ENCODE vs GTEx). Only minor performance differences are observed between the two data sources, such as slightly lower correlation for splice junctions derived from ENCODE data compared to GTEx.

**Supplementary Figure 6.**
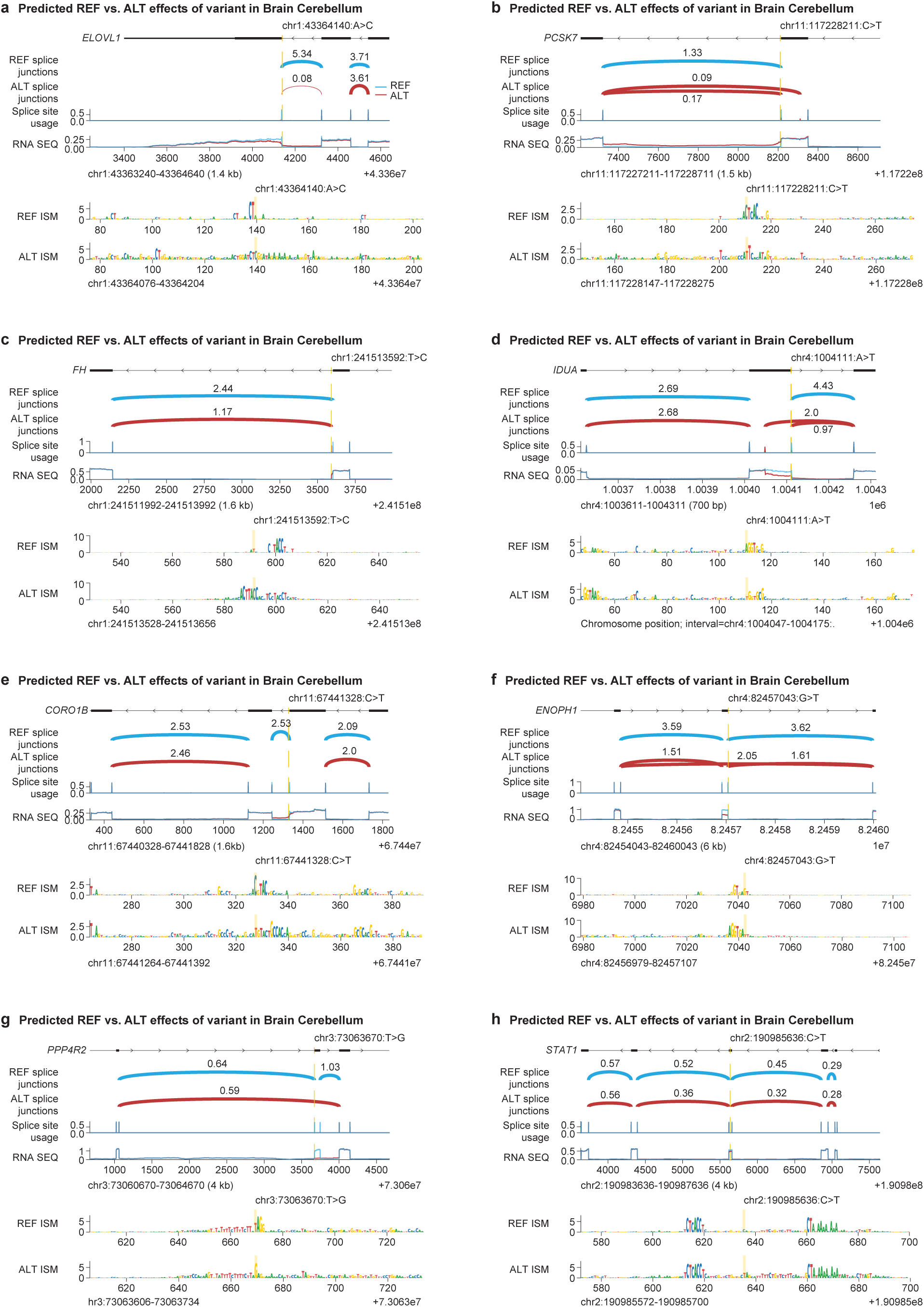
**AlphaGenome predictions on validated de novo cryptic splice mutations in autism spectrum disorder (ASD) individuals from the SpliceAI study**^15^**. (a-b)** Reference and Alternate Splice Junction, Splice Site Usage and RNA-seq AlphaGenome predictions on cases with validated intron retention with the variant. ISM on both the Reference and Alternate sequences is also shown. **(c-e)** Example predictions where the variant results in the creation of novel junctions. In examples (c) and (d), AlphaGenome clearly predicts the novel junctions, whereas in example (e) the model predicts an intron retention event. **(f-h)** Example predictions where the variant results in exon skipping. In examples (f) and (g), AlphaGenome correctly predicts the exon skipping event, whereas in example (h) it misses it.

**Supplementary Figure 7.**
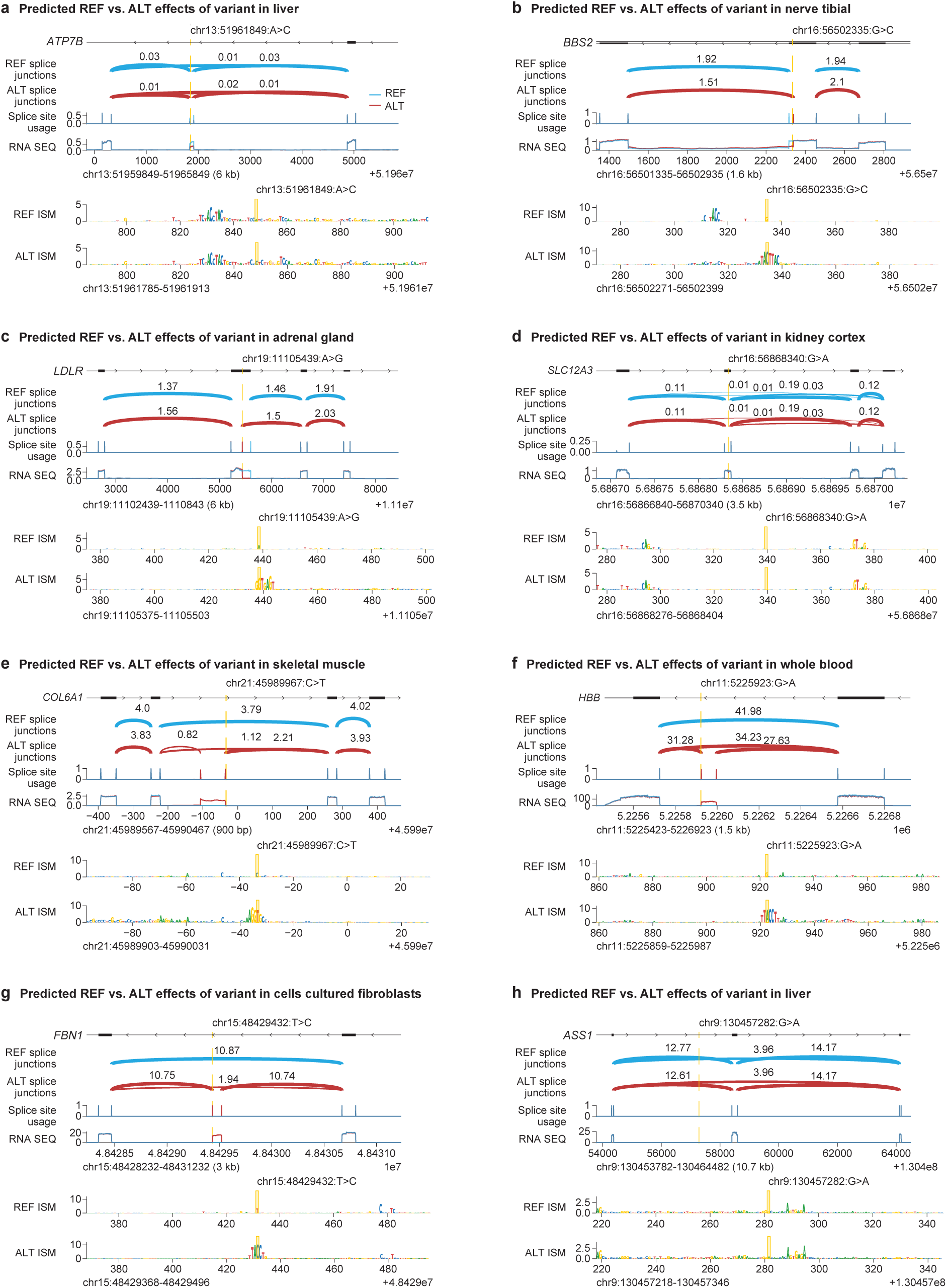
AlphaGenome predictions for missense and deep intronic variant examples from ClinVar. Success and failure examples. **(a)** Reference and Alternate Splice Junction, Splice Site Usage and RNA-seq AlphaGenome predictions on a missense variant associated with Wilson disease^16^. ISM on both the reference (REF) and alternate (ALT) sequences is also shown. AlphaGenome predicts an exon skipping event in the presence of the variant (𝑅𝑒𝑓 𝑃 𝑆𝐼 = 0.75, 𝐴𝑙𝑡 𝑃 𝑆𝐼 = 0.33). **(b-c)** AlphaGenome predictions indicate the creation of novel junctions for two missense variants associated with (b) Bardet-Biedl syndrome^16,17^ and (c) familial hypercholesterolemia5. **(d)** AlphaGenome fails to predict a splicing effect for a missense variant associated with Gitelman syndrome^18^. **(e-g)** AlphaGenome predictions on ClinVar pathogenic deep intronic (distance to closest splice junction > 50 bp) variants. Predictions indicate exon inclusion events due to intronic variants associated with (e) Bethlem myopathy^19^ and Ullrich congenital muscular dystrophy^20^, (f) beta-thalassemia^21,22^, and (g) Marfan syndrome^21^.

**Supplementary Figure 8.**
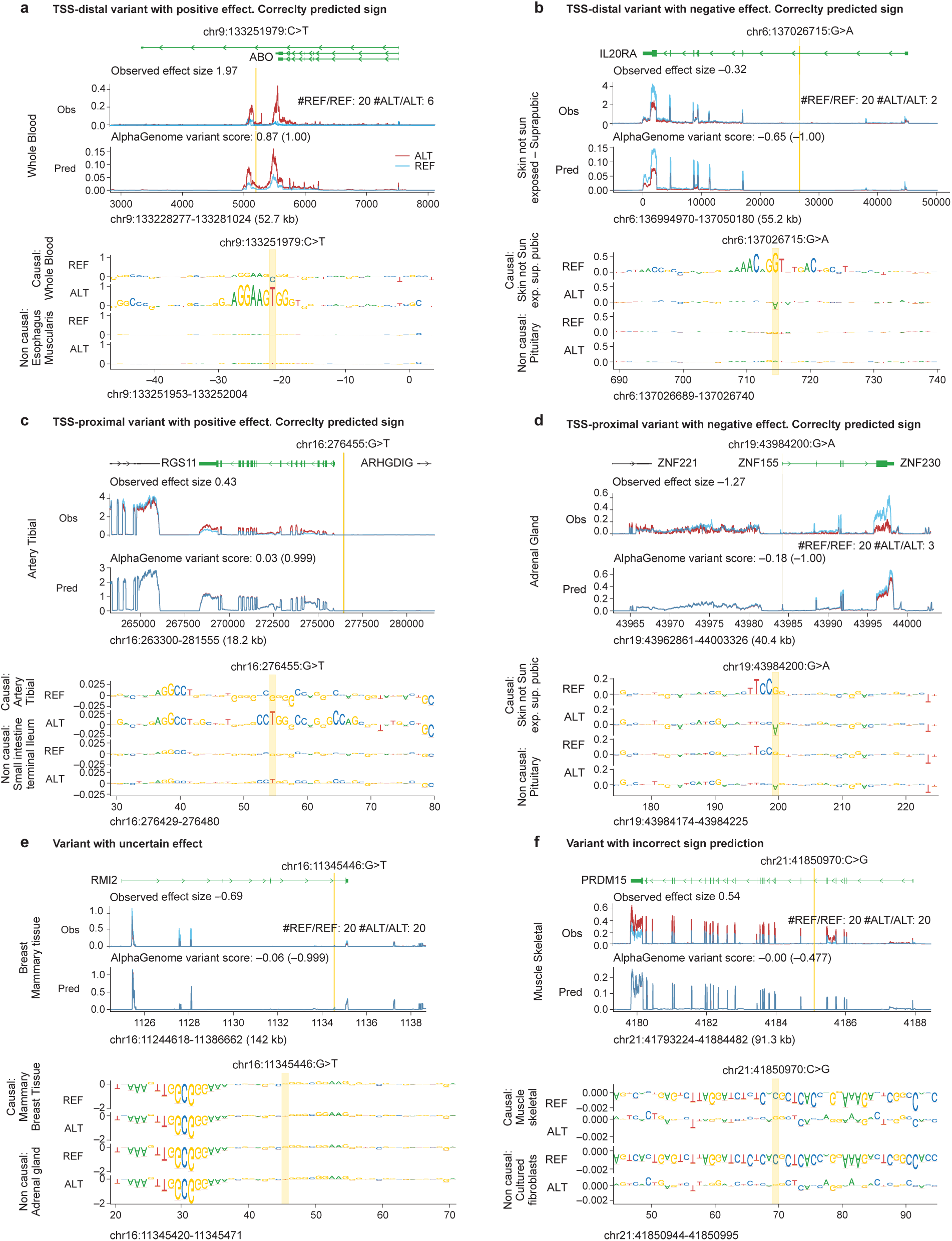
AlphaGenome eQTL effect predictions. **(a)** Example of distal variant (12-35kb from the target gene TSS) with a positive effect and correctly predicted sign. The variant generates a new ETS-like binding site resembling a known binding site (Jaspar matrix #MA0080.2) for SPI1, a critical transcription factor for myeloid and lymphoid cells development. ISM scores highlight how the motif is only relevant in the GTex tissue where this eQTL was identified, Whole Blood. **(b)** Example of distal variant with a negative effect and correctly predicted sign, with the variant leading to reduced gene expression. **(c)** Example of proximal variant with positive effect and correctly predicted sign. The variant generates a CCTGG sequence, a known non-CpG target for DNA methylation associated with gene repression. **(d)** Example of proximal variant with negative effect and correctly predicted sign. The variant disrupts the edge of an ETS1-like motif (Jaspar matrix #MA0098.1) leading to strongly reduced motif score and correctly predicted reduction in gene expression. **(e)** Example of an ambiguous case for a distal variant with correctly predicted sign. ISM scores do not highlight any relevant motif for either alleles overlapping the variant. Only a tissue non-specific motif is evident nearby. **(f)** Example of failure case on a variant with incorrectly predicted effect and very low ISM scores.

**Supplementary Figure 9.**
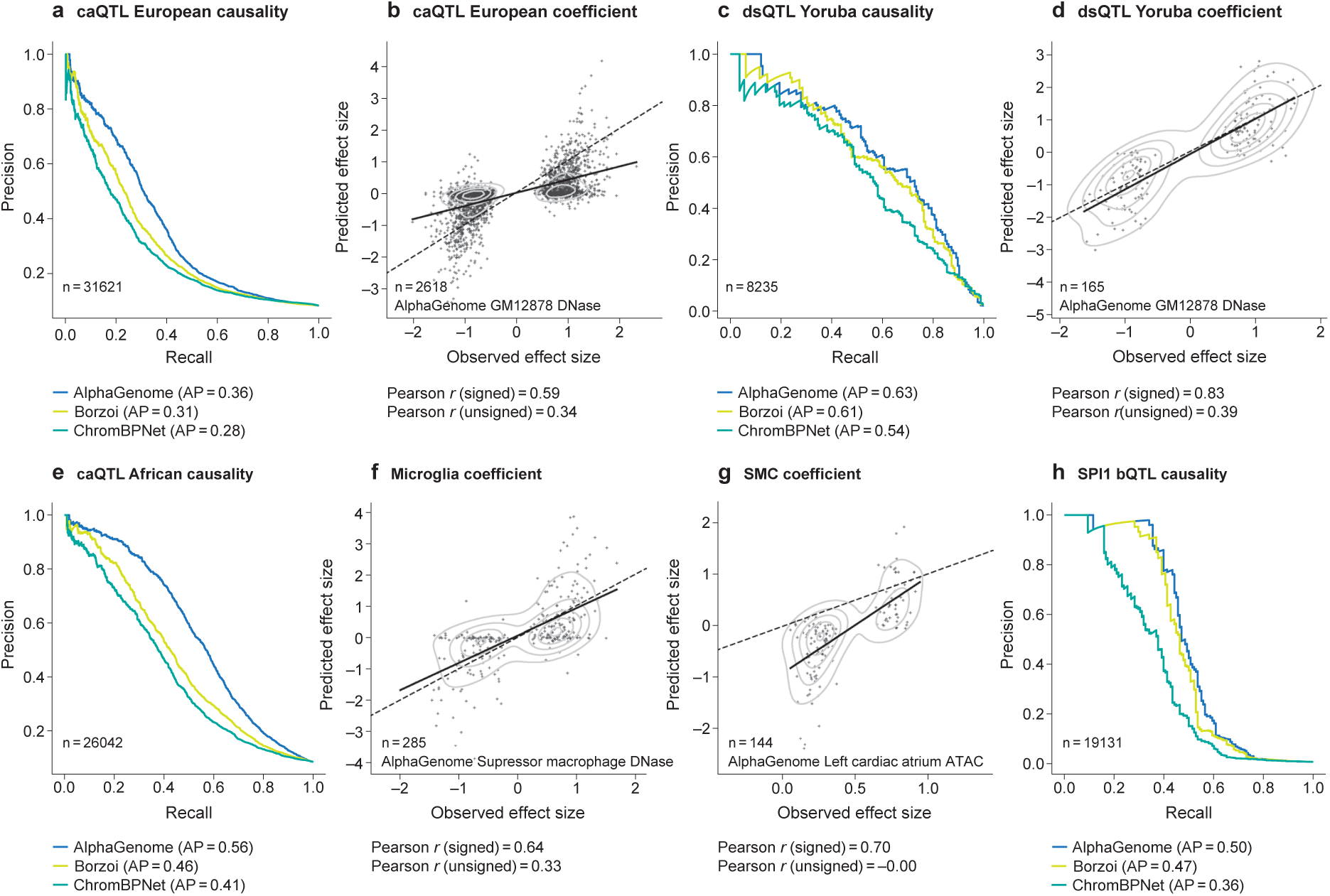
Additional accessibility variant analysis. Extended evaluation of variant effect prediction on chromatin accessibility across diverse contexts. AP = average precision (auPRC). Signed Pearson R correlation uses raw values; unsigned Pearson R uses absolute values first. **(a)** Precision-Recall curves comparing AlphaGenome, Borzoi, and ChromBPNet performance on caQTL causality prediction in European ancestry. **(b)** Scatterplot comparing AlphaGenome’s predicted versus observed effect sizes (Coefficient) for causal caQTL variants in European ancestry. **(c)** Precision-Recall curves comparing AlphaGenome, Borzoi, and ChromBPNet performance on dsQTL causality prediction in Yoruba ancestry. **(d)** Scatterplot comparing AlphaGenome’s predicted versus observed effect sizes (Coefficient) for causal dsQTL variants in Yoruba ancestry. **(e)** Precision-Recall curves comparing model performance for caQTL causality prediction (African ancestry). **(f)** Effect size prediction for microglia causal caQTL variants. Scatterplot compares observed effects versus AlphaGenome’s predicted DNase effects in a closely-related available cell type (suppressor macrophage). **(g)** Effect size prediction for cardiac smooth muscle cell (SMC) causal caQTL variants. Scatterplot compares observed effects versus AlphaGenome’s predicted ATAC effects in a closely-related available cell type (left cardiac atrium ATAC). **(h)** Precision-Recall curves comparing model performance for SPI1 bQTL causality prediction.

**Supplementary Figure 10.**
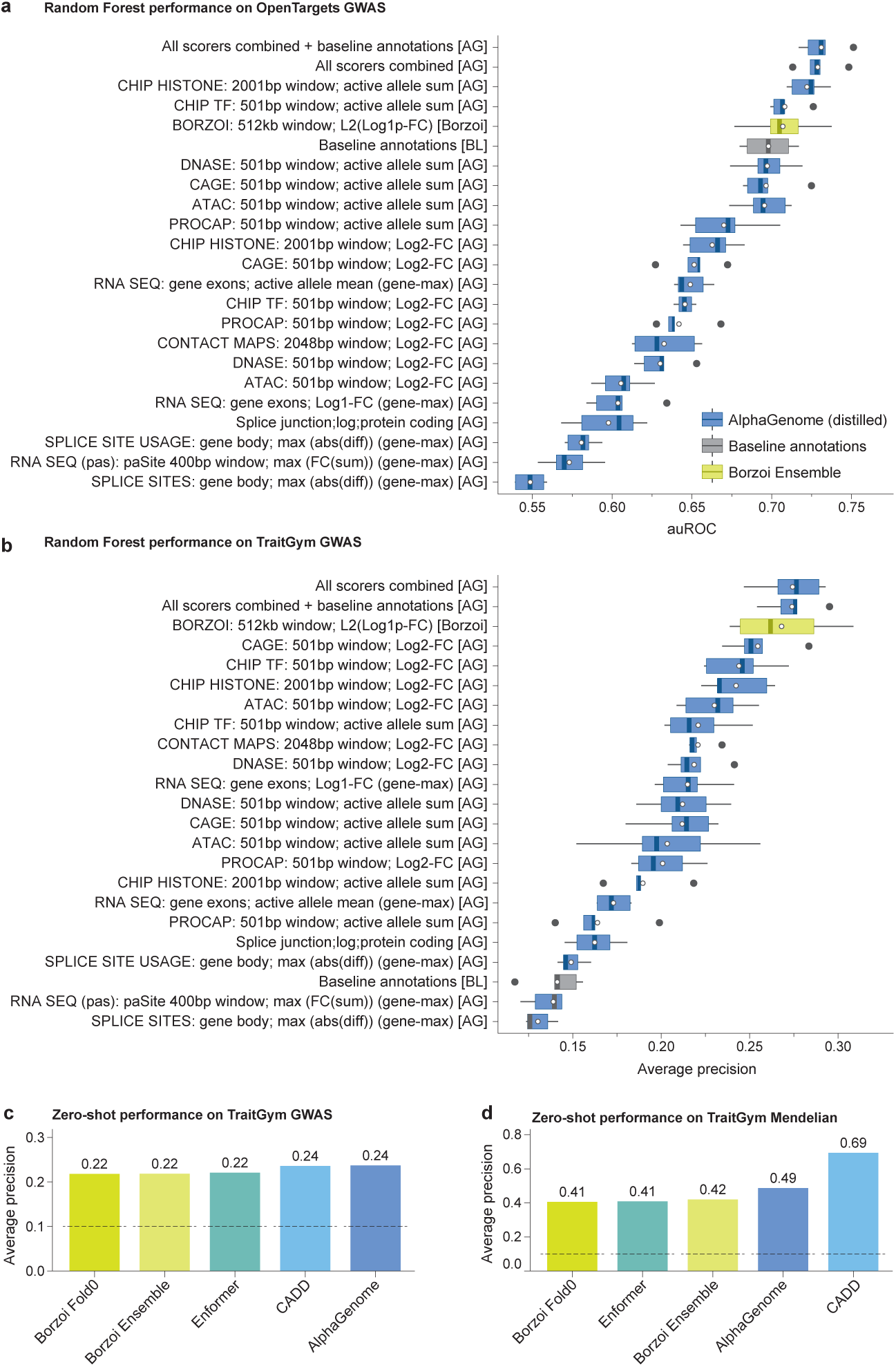
Evaluating models on complex and mendelian traits. **(a)** AUROC for classifying whether variants are implicated in complex traits using random forests trained on model predictions or baseline features (𝑛 = 7636). This evaluation uses positive and negative variants derived from the OpenTargets dataset^23^ (Methods). Random forests were evaluated using five-fold cross-validation, white dot indicates the mean across folds. Whiskers represent 1.5 times the interquartile range. **(b)** Average Precision (auPRC) for classifying whether variants are implicated in complex traits using random forests trained on model predictions or baseline features (𝑛 = 11400). This evaluation uses causal and control variants as defined in Traitgym^24^. Random forests were evaluated using five-fold cross-validation, white dot indicates the mean across folds. Whiskers represent 1.5 times the interquartile range. **(c)** Average Precision (auPRC) for classifying whether variants are implicated in complex traits using aggregated zero-shot model predictions (Methods, 𝑛 = 11400). This evaluation uses the same variants as in (b). However, as no random forest is trained, no cross-validation is performed. Grey line indicates random performance (class balance). All methods, including CADD, exhibit moderate predictive power. **(d)** Average Precision (auPRC) for zero-shot classification of whether a variant is implicated in a mendelian trait or is a matched negative control, sampled from common variants (𝑛 = 3380). Causal and control variants are taken from TraitGym. AlphaGenome outperforms previous sequence-to-function models. However, CADD strongly outperforms all other methods, likely because it can leverage conservation information to separate rare from common variants. Grey line and error bars as in (c).

**Supplementary Figure 11.**
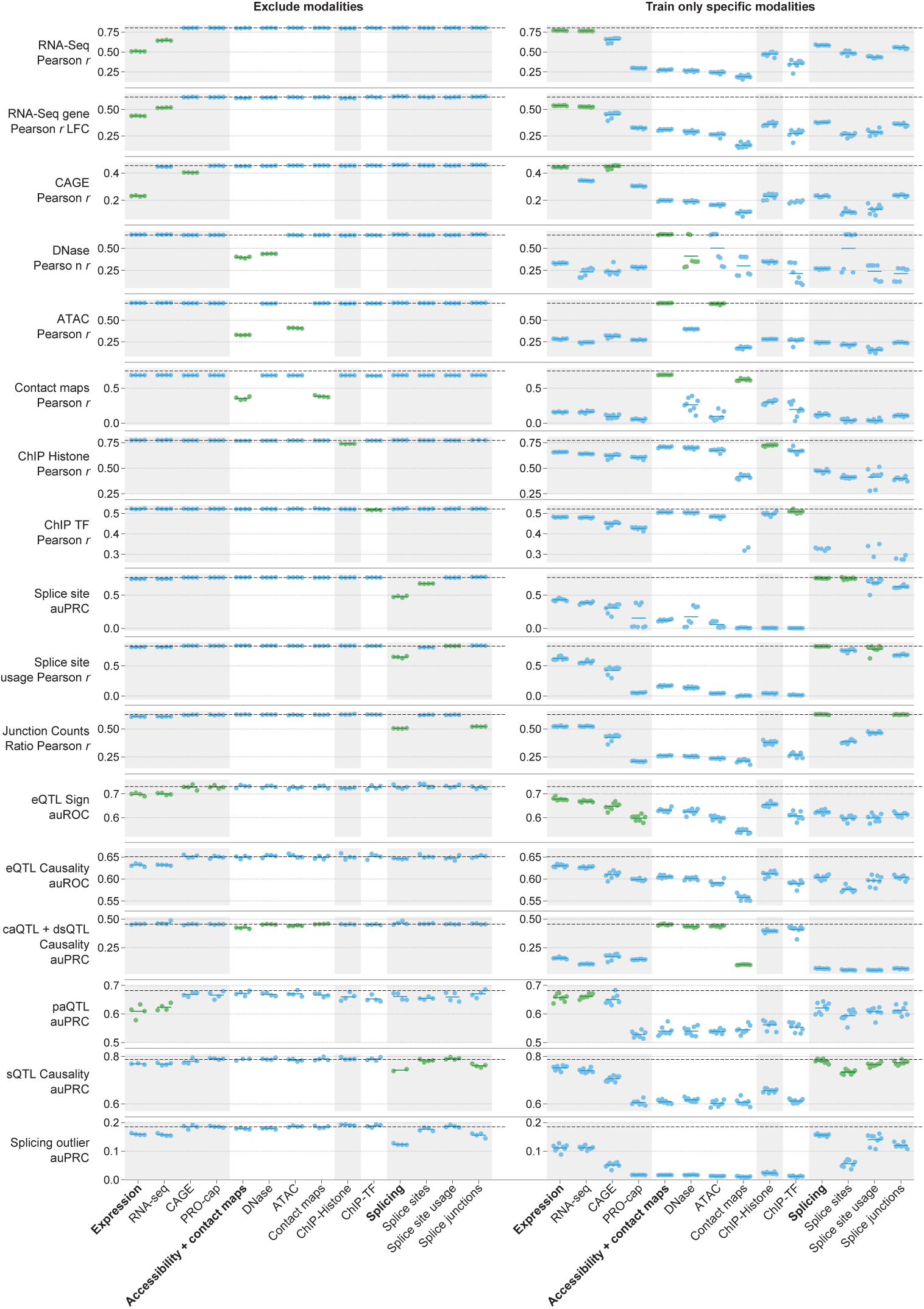
Impact of excluding or isolating single modalities on track and variant tasks. Comparison of AlphaGenome performance on variant and track prediction tasks when training excludes a single output/modality group versus when training only a single modality group. The x-axis indicates the specific output head (e.g., ’CAGE’; regular size label) or the broader modality group (Expression, Accessibility + contact maps, and Splicing; bolded label) targeted by the ablation. Each dot represents an independent training run (n=4 for ’Exclude Modalities’ and n=8 for ’Train only Specific Modalities’ random seeds); black dashed lines indicate the mean performance of the full multi-task model for reference; green dots indicate when the tasks is related to the ablated modality and blue is the default color.

**Supplementary Figure 12.**
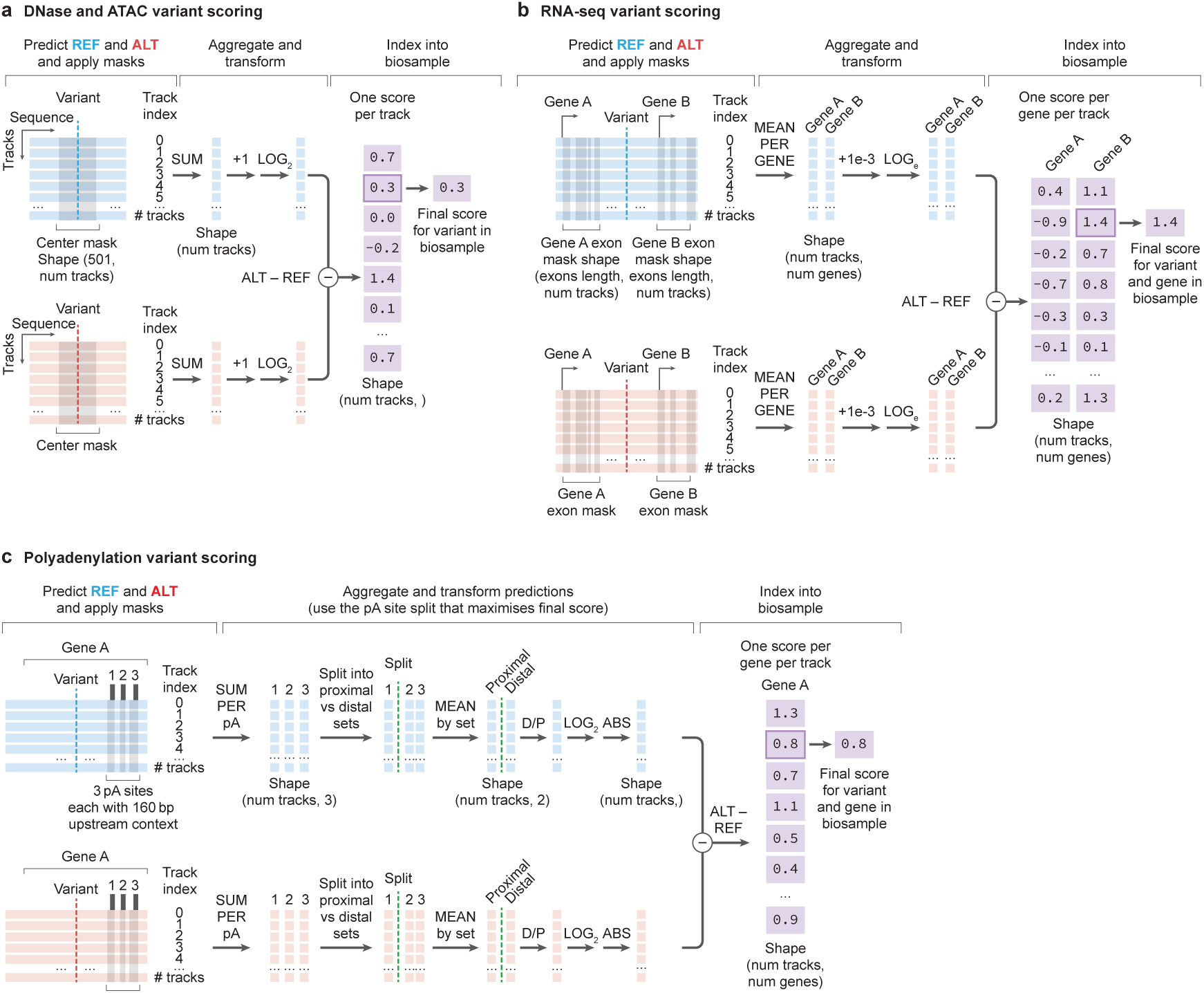
**Illustrative workflows for selected variant scoring mechanisms. (a) Chromatin accessibility variant scoring (i.e. DNase/ATAC)**. Schematic illustrating a center-mask based approach. Signals from reference (REF) and alternative (ALT) allele predictions are summed within a window, log2-transformed, and then subtracted to quantify variant impact on local chromatin accessibility. **(b) RNA-seq gene expression variant scoring**. Schematic for gene-based scoring of expression changes. REF and ALT predictions are averaged within gene boundaries, log-transformed, and then subtracted to estimate a log-fold change per gene. **(c) Polyadenylation variant scoring** Schematic for assessing variant effects on polyadenylation site usage. The workflow involves comparing REF and ALT predictions at potential polyA sites to determine changes in relative proximal versus distal site usage, summarized as a maximum log-fold change of isoform ratios.

**Supplementary Figure 13.**
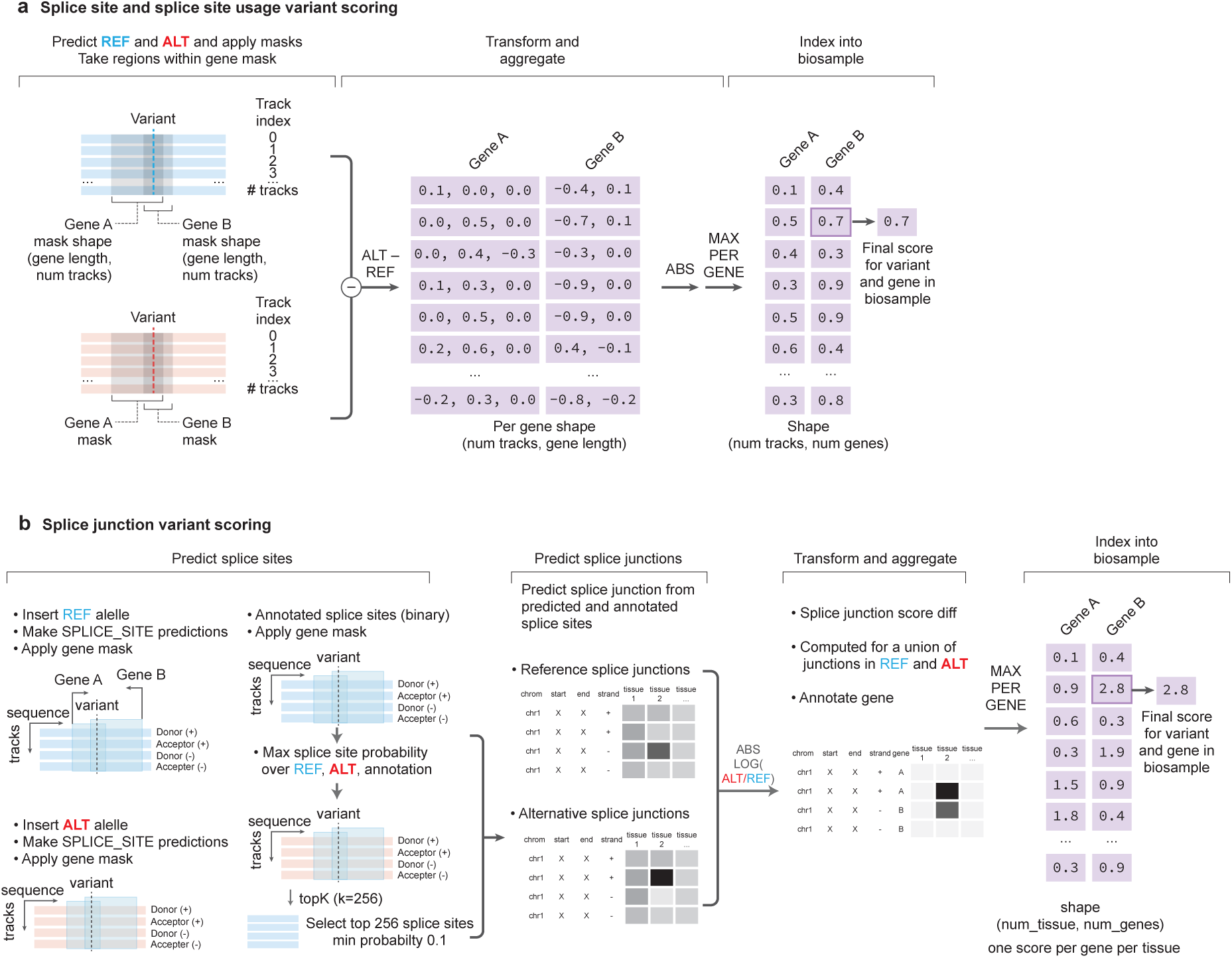
**Illustrative workflows for splicing variant scoring mechanisms. (a) Splice site and splice site usage variant scoring**. Schematic for gene-based splicing impact. Differences between REF and ALT splicing predictions are calculated within gene body regions, an absolute value is taken, and the maximum absolute difference across relevant sites forms the variant score. **(a) Splice junction variant scoring**. Schematic for gene-based splicing impact scored with the splice junction head. Only predicted and annotated splice sites from the genes overlapping the variant are considered for splice junction predictions. Absolute log fold change on splice junction scores is computed for all predicted junctions. A maximum score per tissue is used as the variant score for that tissue.

